# PANCS-Inhibitors: A rapid method to directly select for protein-protein interaction inhibitors

**DOI:** 10.64898/2026.08.05.743105

**Authors:** Matthew J. Styles, Victoria Cochran Xie, Sandrine Legault, Joshua A. Pixley, Bryan C. Dickinson

## Abstract

Aberrant protein-protein interactions (PPIs) drive myriad diseases. Inhibiting these PPIs often relies on discovering molecules that bind to one of the proteins and hoping that this binding inhibits the PPI. Molecular binder discovery often takes months, but a discovery process that ensures that the resulting molecule not only binds a target protein, but selectively inhibits a target PPI, could dramatically accelerate these endeavors. Here, we develop Phage-Assisted Non-Continuous Selection of PPI Inhibitors (PANCS-Inhibitors): a rapid screening platform that directly selects for molecules capable of disrupting a pre-formed PPI. We demonstrate this new platform using three clinically relevant oncogenic PPIs: KRas-Raf, Mdm2-p53, and Myc-Max. PANCS-Inhibitors can be used to both improve known PPI inhibitors and for *de novo* discovery of mini-protein PPI inhibitors that function in mammalian cells. This platform has the potential to rapidly generate inhibitors for many clinically relevant PPIs, which can be used as starting points for therapeutic development.

## Introduction

An estimated 130,000 human protein-protein interactions (PPIs)^1^ regulate virtually every cellular process, including replication,^2,3^ translation,^4^ and signal transduction.^5–7^ Advances in unbiased proteome-wide PPI mapping methods such as 2-hybrid screens,^8–10^ proximity labeling technologies,^11,12^ CRISPR base-editing screening,^13^ and advanced computational predictions^14,15^ have enabled extensive mapping of human PPIs. Dysregulation of specific PPIs drive pathology in humans,^16–18^ and therefore represent critical therapeutic targets for disease intervention.^19–22^ Despite their clear importance, tools for precisely ablating a specific PPI are lacking: simple genetic knockouts involves additional, confounding changes as most proteins engage in multiple PPIs and can have moonlighting functions.^23^ As such, molecules that specifically disrupt target PPIs are critical for both assigning the functional significance of mapped PPIs as well as for creating therapeutics.

Discovery and development of PPI inhibitors is inherently challenging.^6^ The most common approach for PPI inhibitor discovery entails finding a competitive binding partner to one of the proteins involved in the interaction, usually through binding at the PPI interface. This can be done via high-throughput screening,^24–26^ rational design,^27^ computational modeling,^28,29^ and binding-based directed evolution platforms.^30–33^ However, the underlying issue for all binding-based discovery campaigns is that binding does not necessarily confer inhibition; the function selected for is not the desired end function. Competitive PPI inhibition can be obtained by directing the binding epitope of a *de novo* or designed binder.^28,34^ Conversely, allosteric inhibitors can be difficult to discover from binder-first approaches, as allosterically regulated sites can be difficult to predict *a priori*. An optimal strategy to discover inhibitors would be to directly select for inhibition of a preformed PPI in a mechanistically agnostic manner. The reverse 2-hybrid selection design directly links PPI disruption to selection pressure, but has yielded only a few successes.^33,35–40^ We recently reported Phage-Assisted Non-Continuous Selection of Binders (PANCS-Binders)^41^ as a rapid, high fidelity means of screening up to 10^10^ variant libraries for *de novo* protein binders. Yang *et al.* adapted this platform for the discovery of aggregation inhibitors (aggregation of one half of a split RNAP impeded its function, inhibition of aggregation restored function).^33^ Additionally, we have demonstrated that our split RNAP can be used for the direct selection of PPI inducers.^42^ We sought to adapt our inducer biosensor and PANCS protocols for direct, unbiased selection of PPI inhibitors in a generalizable format to rapidly select for PPI inhibitors in a mechanistically agnostic manner.

In this work, we develop Phage-Assisted Non-Continuous Selection of PPI inhibitors (PANCS-Inhibitors), which directly links the life cycle of an M13 bacteriophage to the selective disruption of a target PPI. After optimizing PANCS-Inhibitors to detect inhibitors of three cancer related PPIs (KRas-Raf, Mdm2-p53, and Myc-Max) in mock selections, we demonstrate the capabilities of PANCS-Inhibitors in a series of selections. First, because of the simultaneous high fidelity and large library sizes, we sought to perform a selection in which the end goal was not discovery, but to characterize the underlying fitness landscape of a PPI inhibitor. Thus, we performed a deep mutational scan (DMS), comprehensively covering four randomized positions within the Ras binding domain of cRAF (Raf) to understand how well single and double mutations capture the relevant epistasis needed to predict multi-mutant fitness.^43,44^ Second, we used PANCS-Inhibitor selections to identify *de novo* Mdm2-p53 and Myc-Max protein-based inhibitors that function in mammalian cells. This work showcases the capacity of PANCS-Inhibitors for robust, rapid, and mechanistically agnostic study and discovery of PPI inhibitors.

## Results

### Adaptation of the split RNAP biosensor to detect PPI inhibition in *E. coli*

In *de novo* binder selections, binders might potentially bind at one of many “hot-spots” on the target protein. These sites, while often sites of natural PPIs, may not disrupt the desired PPI (**Fig. 1a**). The oncogenic signaling protein, KRas, exemplifies this trend. Ras isoforms have been the subject of numerous binder discovery campaigns and numerous hot-spots have been identified (**Fig. S1**); however, most binders do not inhibit the KRas-Raf interaction, either directly or allosterically. Further highlighting this point, each KRas-Raf inhibitor (**Fig. S1**) was discovered through extensive screening of binding variants from selections for binding to KRas, not direct screening for inhibiting the KRas-Raf interaction. We recently reported a high affinity (∼0.2 nM K_d_) KRas binder^41^ which, based on the crystal structure of this PPI^45^, we did not anticipate would inhibit the KRas-Raf interaction (**Fig. S1**); we confirmed this (*see below*). As binder identification takes significant effort and resources, discovery and evaluation of binders that do not confer inhibition of the desired PPI is a hinderance to inhibitor development.

**Fig. 1:**
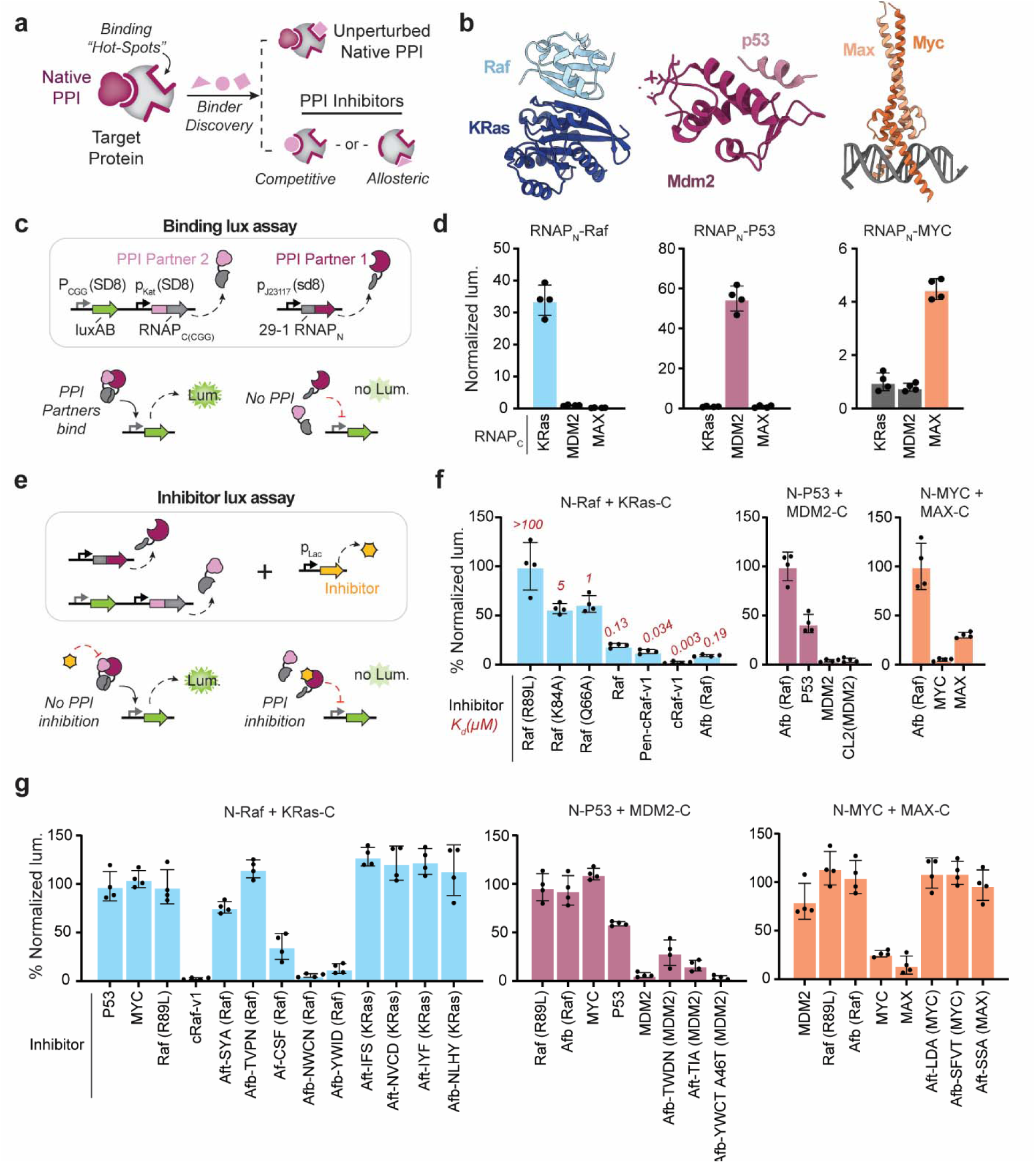
Detecting protein-protein interaction inhibition in *E. coli*. **a,** Binding does not necessarily confer inhibition as the most likely site for a *de novo* binder, a binding hot-spot, is not necessarily at a site that directly competes with binding or at a site for allosteric regulation. **b**, Structures of target PPIs for this study: KRas-Raf (PDB:6XHB), Mdm2-p53 (PDB:4HFZ), Myc-Max (PDB:1NKP). **c**, Proximity dependent split-RNA polymerase (RNAP) complementation assay for the detection of binding: PPI Partner 1 is fused to the RNAP_N_ fragment and PPI Partner 2 is fused to the RNAP_C_ fragment, each is expressed from a separate promoter system, and RNAP complementation drives LuxAB expression. **d**, Binding luciferase assays for on- and off- target PPIs (n=4; error bars indicate SD; see **Table S1** for protein sequences) after optimization of linker length and orientation (**Fig. S2**). **e**, Proximity-dependent split-RNA polymerase (RNAP) complementation assay for the detection of protein-protein interaction inhibition in *E. coli*: to the optimized PPI detection pairings (**c**), a potential inhibitor is expressed from a third plasmid. **f**, Detection of PPI inhibition in our *E. coli* reporter (n=4; error bars indicate SD) for various on- and off- target PPI partners and known inhibitors (affinities reported previously).^30,49–51^ **g**, Testing of previously identified binders for inhibition of our target PPIs. (n=4; error bars indicate SD; see **Table S1** for protein sequences and information on prior binder discovery).

We sought to develop a screening assay to test if our previously reported binders^41^ functioned as PPI inhibitors. We selected three biophysically distinct, but clinically important, PPIs to focus on: KRas-Raf, Myc-Max, and Mdm2-p53 (**Fig. 1b**; see **Table S1** for sequences/details of prior binder discovery campaigns). We monitored binder-target interactions using our previously reported split RNAP *E. coli* luciferase binding reporter assay^41,46^. In this assay, a proximity-dependent complementation between the split RNAP controls the expression of a luciferase reporter gene wherein protein binding drives the degree of luciferase signal (**Fig. 1c**). After optimizing linker lengths and orientation of the PPI across the split RNAP components, we observed robust PPI detection for each of our PPI pairs in comparison to off-target pairings (**Fig. 1d; see Fig. S2 for additional negative controls**). Next, we adapted our previously reported PPI inducer *E. coli* luciferase assay^42^ to monitor PPI inhibition. To our optimized binding *E. coli* luciferase PPI/split RNAP combinations, we additionally expressed a potential inhibitor from a third plasmid (**Fig. 1e**). Expression of either partner of a PPI pair or known inhibitors resulted in significant decreases in luciferase signal in an affinity dependent manner (**Fig. 1f**). Our previously reported Mdm2 binders were expected to inhibit the Mdm2-p53 interaction based on the presence of a known Mdm2 binding motif, AlphaFold3^47^ predictions (**Fig. S3**), and our phenotypic demonstration that these binders inhibit the Mdm2-p53 interaction in mammalian cells.^41^ None of our other binders were tested for PPI inhibition; however, seven out of 15 have high confidence (iPTM >0.5) AlphaFold3 predictions, four of which indicate likely competitive inhibition (for low confidence predictions, six out of eight are predicted to inhibit; **Fig. S3**). We tested our binders for KRas, Raf, Mdm2, Myc, and Max (no p53 binders were identified) in our inhibitor luciferase assay (**Fig. 1g**). As expected, each Mdm2 binder tested inhibited the Mdm2-p53 interaction; however, none of the KRas binders inhibited the KRas-Raf interaction, none of the Myc or Max binders inhibited the Myc-Max interaction, and only a subset of Raf binders (three out of five) inhibited the KRas-Raf interaction (**Fig. 1g**). Notably, only Raf binders identified from large library selections (10^10^ affibody variants) were inhibitors - those identified from smaller, 10^8^ libraries were not - suggesting that the binding motif that engenders KRas-Raf inhibition may be a rare binding motif.^48^ As PPI inhibition can be monitored with our split RNAP and our binder selections do not generally result in inhibitors, we sought to modify our PANCS-Binders split RNAP biosensor for direct selection of PPI inhibitors.

### Development of the PANCS-Inhibitors platform

Our PANCS-Binders platform,^41^ which is based on the principles of Phage-Assisted Continuous Evolution (PACE),^52^ uses replication deficient M13 bacteriophage (the essential *gIII* is removed from the phage genome) and *E. coli* selection strains which use a biosensor to control *gIII* expression. Phage that encode a gene variant that activates the biosensor are able to replicate, while phage that encode variants that do not activate the biosensor are unable to replicate (**Fig. S4**). This links sequence (phage encoded variant) to fitness (phage replication). In PANCS-Binders (**Fig. S5**), phage encode the N-terminal half of the split RNAP (RNAP_N_) fused to a potential binding variant in the phage genome (SP), and the *E. coli* selection strain encodes the C-terminal half of the split RNAP (RNAP_C_) fused to the target protein of interest and RNAP dependent control of *gIII* on a plasmid (+AP). To prevent replication of variants that simply bind the RNAP_C_, we also include in the *E. coli* selection strain an orthogonal RNAP_C_ fused to an off-target protein and orthogonal RNAP-dependent control of a dominant negative variant of *gIII* (*gIII_neg_*), which poisons phage replication when expressed. We sought to modify this system such that phage replication would be dependent on selective inhibition of the desired PPI.

After attempting several simple but unsuccessful strategies to modify the split-RNAP biosensor (see **Supplementary Note 1**), we developed Phage-Assisted Non-Continuous Selection of Inhibitors (PANCS-Inhibitors), which successfully links the production of an inhibitor for a preformed target PPI (PPI Partner 1 and PPI Partner 2) to phage replication (**Fig. 2a**). A first accessory plasmid (AP1) encodes: (i) a constitutively expressed zipper peptide 2 (ZP2) fused to zipper peptide A (ZA) and PPI partner 1; (ii) a constitutively expressed zipper peptide B (ZB) fused to the C-terminal half of the split CGG RNAP (RNAP_C(CGG)_), which transcribes from the CGG promoter (P_CGG_) when activated; and (iii) P_CGG_-driven *gIII*. A second accessory plasmid (AP2) encodes: (i) a constitutively expressed PPI partner 2 fused to the C-terminal half of the split T7 RNAP (RNAP_C(T7)_), which transcribes from the T7 promoter (P_T7_) when activated and (ii) P_T7_-driven *gIII_neg_*. Prior to phage infection, ZA and ZB form a PPI, and PPI Partner 1 and PPI Partner 2 form a PPI, but neither *gIII* nor *gIII_neg_*are expressed, because RNAP_N_ (which can complement either RNAP_C_) is not present in the system. We then engineered M13 bacteriophage that encodes: (i) RNAP_N_ fused to zipper peptide 1 (ZP1), which binds ZP2, and (ii) a potential inhibitor molecule of the target PPI. Upon phage infection, ZP1 binds to the ZP2 on both the on-target (PPI Partner 1 and PPI Partner 2) and off-target (ZA-ZB) PPI complexes, triggering both *gIII* and *gIII_neg_* production and preventing phage replication. However, if the inhibitor encoded by the phage selectively disrupts the pre-formed on-target PPI, then *gIII_neg_*production is blocked, allowing that phage variant to replicate. This construct prevents expression of *gIII* or *gIII_neg_* prior to phage infection (which is necessary as *gIII* expression inhibits phage infection) and minimizes possible routes by which mutations in the phage could ‘cheat’ the biosensor (mutations to the RNAP_N_-ZP1 cannot selectively favor complementation with RNAP_C (CGG)_ for *gIII* production). Critically, the only difference between the two trimolecular complexes formed in the absence of an inhibitor is the target PPI versus the ZA-ZB PPI, so inhibitors of the split RNAP, ZP1-ZP2, or ZA-ZB will prevent phage growth by not allowing the expression of *gIII*; therefore, the selection pressure is entirely focused on selective disruption of the target PPI. Having established the conceptual framework of PANCS-Inhibitors, we next sought to optimize the system and assess its performance characteristics as a selection platform.

**Fig. 2:**
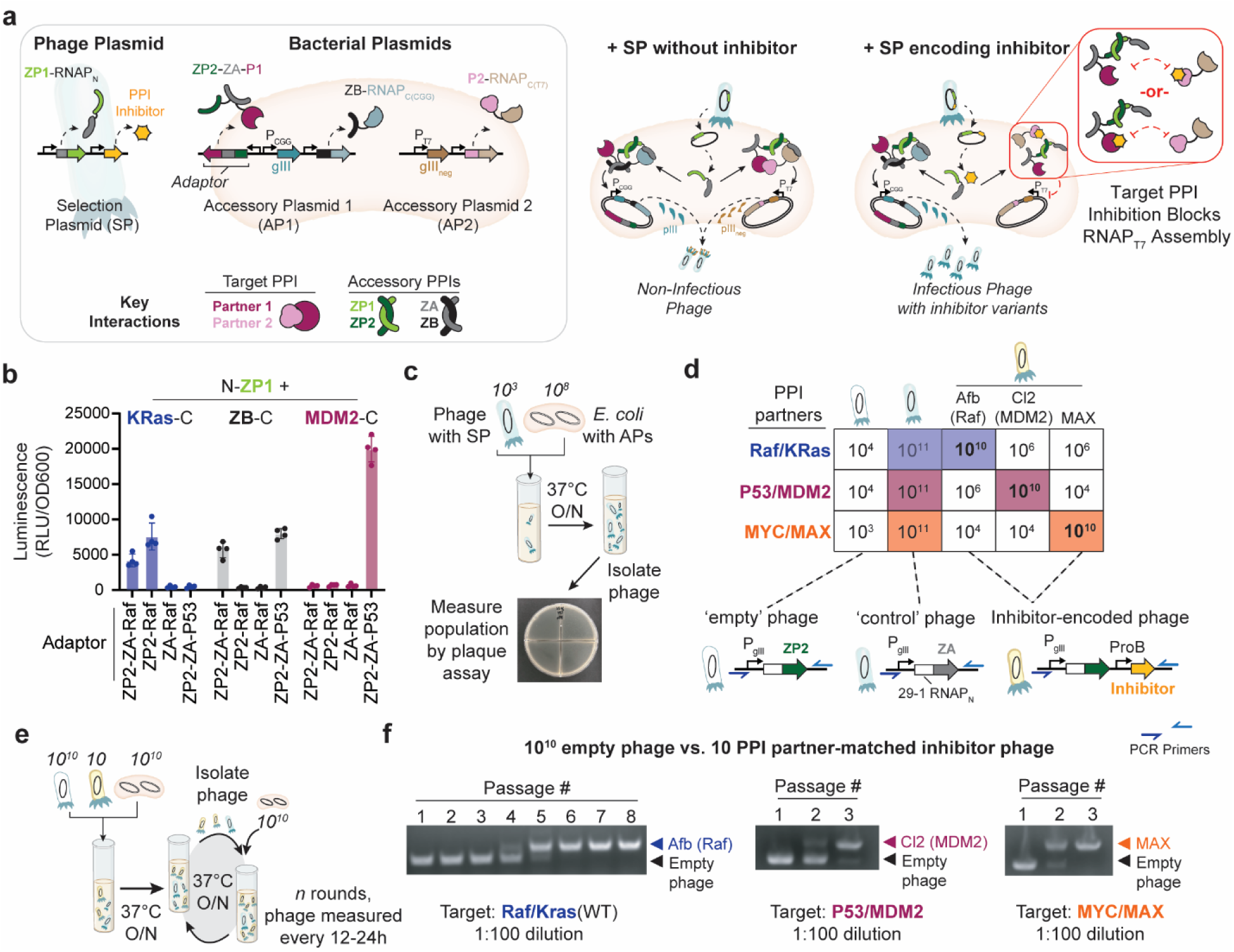
Developing the PANCS-Inhibitors platform. **a,** PANCS-Inhibitors biosensor concept: three genetic constructs (AP1, AP2, and SP). AP1 encodes three components: ZB fused to RNAP_C (CGG)_, RNAP_CGG_ dependent *gIII*, and an adaptor (a fusion of PPI partner 1 (e.g. Raf), ZA, and ZP2. AP2 encodes two components: PPI Partner 2 (e.g. KRas) fused to RNAP_C (T7)_ and RNAP_T7_ dependent *gIII_neg_* expression. The SP encodes two components: a potential inhibitor and RNAP_N_ fused to ZP1. There are several key interactions: ZA binds ZB, ZP1 binds ZP2, and PPI Partner 1 (e.g., Raf) binds PPI Partner 2 (e.g., KRas). Prior to phage infection, the ZA-ZB interaction complex forms and the target PPI (e.g., Raf- KRas) interaction complex forms, but neither RNAP_C_ can induce expression as RNAP_N_ is absent. Upon phage infection, RNAP_N_ is brought to both the ZA-ZB and target PPI interaction complex via the ZP1-ZP2 interaction, thus activating RNAP complementation and associated transcription of *gIII* and *gIII_neg_* inhibiting phage replication. However, in the case where an inhibitor was encoded in the phage, the target PPI is disrupted preventing expression of *gIII_neg_* allowing for robust phage replication (right). **b**, *E. coli* inducer luciferase reporter assay demonstrating that the adaptor can mediate the relevant PPIs and RNAP activation of RNAP_N_-ZP1 and PPI Partner 1-RNAP_C_ or ZB-RNAP_C_ with associated negative controls (n=4, error bars indicate SD). **c**, Schematic of the overnight phage replication assay: 1,000 PFU phage are allowed to replicate on 1 mL of selection strain culture overnight prior to determining the phage titer by activity independent plaque assay. **d**, Replication rate on optimized AP2 expression levels (**Fig. S6**); assay performed as a single replicate as depicted in **c**. (Below) Schematics of the three phage types: empty (RNAP_N_-ZP1), control (RNAP_N_-ZA), and inhibitor (RNAP_N_-ZP1, inhibitor); blue arrows indicate the primers used in amplifying phage by PCR in **f** (resulting in different band sizes). **e**, Schematic of mock library PANCS-Inhibitors selections: a mock library is prepared by mixing 10 PFU inhibitor phage and 10^10^ PFU empty phage, this mock library is allowed to replicate on 1 mL of *E. coli* selection strain overnight (12-24 hours), before the phage are isolated by centrifugation and a fraction of this phage is used to seed a fresh *E. coli* selection strain culture to begin the next passage. This cycle is repeated for up to eight total overnight growths (Passages). **f**, Mock selections for inhibitors of KRas-Raf, Mdm2-p53, and Myc-Max were monitored by PCR to identify when the empty phage had de-enriched and the inhibitor phage had de-enriched (PCR bands expected to be observable at ∼10^5^ PFU/mL). Passaging was stopped after Passage 3 for Mdm2-p53 and Myc-Max based on the observed enrichment of inhibitor phage and de-enrichment of empty phage.

First, we demonstrated that the appropriate trimolecular complexes assemble in *E. coli*using our PPI inducer version of our luciferase reporter^42^: RNAP_N_-ZP1, ZP2-ZA-Partner 1, and ZB-RNAP_C_ or Partner 2-RNAP_C_ (**Fig. 2b**). We then cloned and optimized the components of our PANCS-Inhibitor system for each of our target PPIs including positive and negative control phage (**Fig. 2a**). The rate of phage replication is highly dependent on proper tuning of the expression levels of components of the system. Ideally, expression levels are tuned such that phage that do not encode an inhibitor are unable to replicate, but phage that do contain an inhibitor are able to replicate in a gradient fashion (the stronger the inhibition, the greater the replication). To measure phage replication rates, we performed overnight phage replication assays: 1,000 plaque forming units (PFU) of phage stock was added to a 1 mL subculture of selection strain *E. coli* and incubated overnight prior to determining the titer via activity-independent plaque assays (**Fig. 2c**). We varied the expression of each component of our system by altering the strength of the ribosome binding site (RBS) sequence and determined the phage replication rates with various positive control phage (cheater phage which encode RNAP_N_-ZA rather than RNAP_N_-ZP1), on-target inhibitor phage (encodes a known inhibitor), off-target inhibitor phage (encode a mis-matched inhibitor), and a negative control phage (“empty phage” that do not encode an inhibitor; **Fig. 2d**, **Fig. S6**). Under optimized expression levels, positive control phage could replicate 10^8^-fold, empty phage replicate ∼10-fold, on-target inhibitors replicated 10^7^-fold, and off-target inhibitors replicated 1000-fold or less (**Fig. 2d**). In each case, we needed to tune only the AP2 RBS strengths to achieve selective replication rates. For PPIs beyond the scope of this work, tuning the RBS strengths and/or origin of replication on AP1 might be necessary. With these two levers, likely a very wide range of target PPI affinities can be tuned for selective pressure in this biosensor design.

Given the observed selective pressure for phage replication, we used mock selections to demonstrate that the system was tuned appropriately for PANCS. As we can easily generate libraries with 10^9+^ unique variants, we created a mock library by mixing 10 PFU of an active inhibitor phage and 10^10^ PFU of empty phage, to replicate having a single active variant in a 10^9^ library (separate mock libraries were made for each PPI). We conducted our PANCS-Inhibitor selections (**Fig. 2e**) in a manner analogous to PANCS-Binders selections.^41^ Mock library was added to subcultures of our selection strains, followed by overnight growth, pelleting of the cells to isolate the phage containing supernatant, and then seeding a fresh selection strain subculture with a fraction of the supernatant. This process was repeated for eight total growths (passages). We monitored the phage population at the end of each passage by PCR using primers that land up- and downstream of the inhibitor (bands for active inhibitor phage and empty phage have different sizes). The initial input library only shows empty phage by PCR, as the ten active phage present are below the limit of detection (∼10^5^). However, after just 3-8 rounds of serial dilutions, the empty phage populations de-enrich for each target PPI, and the active, inhibitor-encoding, phage appear (**Fig. 2f**; **Fig. S7**), representing at least a 10^14^-fold relative enrichment of inhibitor-encoding phage over non-inhibitor control phage in just a few days. Given the strong performance of PANCS-Inhibitors in these mock selections, we next aimed to challenge the system with libraries of PPI inhibitors.

### PANCS-Inhibitors selections as a deep-mutational scanning tool

In our PANCS-Binders study, we observed high fidelity between rank-order and function: phage with the highest fraction of the endpoint population had the highest affinity.^41^ We expect that PANCS-Inhibitors selections will similarly maintain this rank-order relationship, where relative enrichment strongly correlates to relative inhibition. High fidelity selections enable a critical tool for assessing the fitness landscape of an inhibitor via deep mutational scanning (DMS). Thus, as a proof-of-concept for this new platform, and to assess reproducibility and fidelity, we decided to perform DMS on an inhibitor of the KRas-Raf interaction. Few KRas binders have been shown to inhibit the KRas-Raf interaction (**Fig. S1**). In contrast, the Ras binding domain of Raf, which inhibits the KRas-Raf PPI, has been the focus of several affinity maturation studies,^53–55^ including a high diversity phage display selection that identified a heavily mutated, high affinity Raf variant, cRAFv1).^30^ The KRas-Raf interaction is mediated by intermolecular interaction of an antiparallel β-sheet established by β1-2 of Raf and β2-3 of Ras and an interaction between the α-helix of Raf with the switch 1 loop of Ras. Prior targeted mutational studies have identified two positions on the α-helix of Raf (**Fig. 3a**) where mutations significantly improve affinity (A85K and V88R or Y); however, epistatic effects have been observed where the effect of these mutations are not additive (A85K V88R is lower affinity than either A85K or V88R alone).^56^ Along this helix, mutations likely displace several high energy waters that fill a pocket between KRas and Raf centered around Raf A85 (**Fig. S8**), and engage in additional KRas-Raf interactions. While these efforts have demonstrated that high affinity Raf variants can be discovered, none have done so in a manner that enables a systematic understanding of the epistatic relationships between positions within this helix.

**Figure 3:**
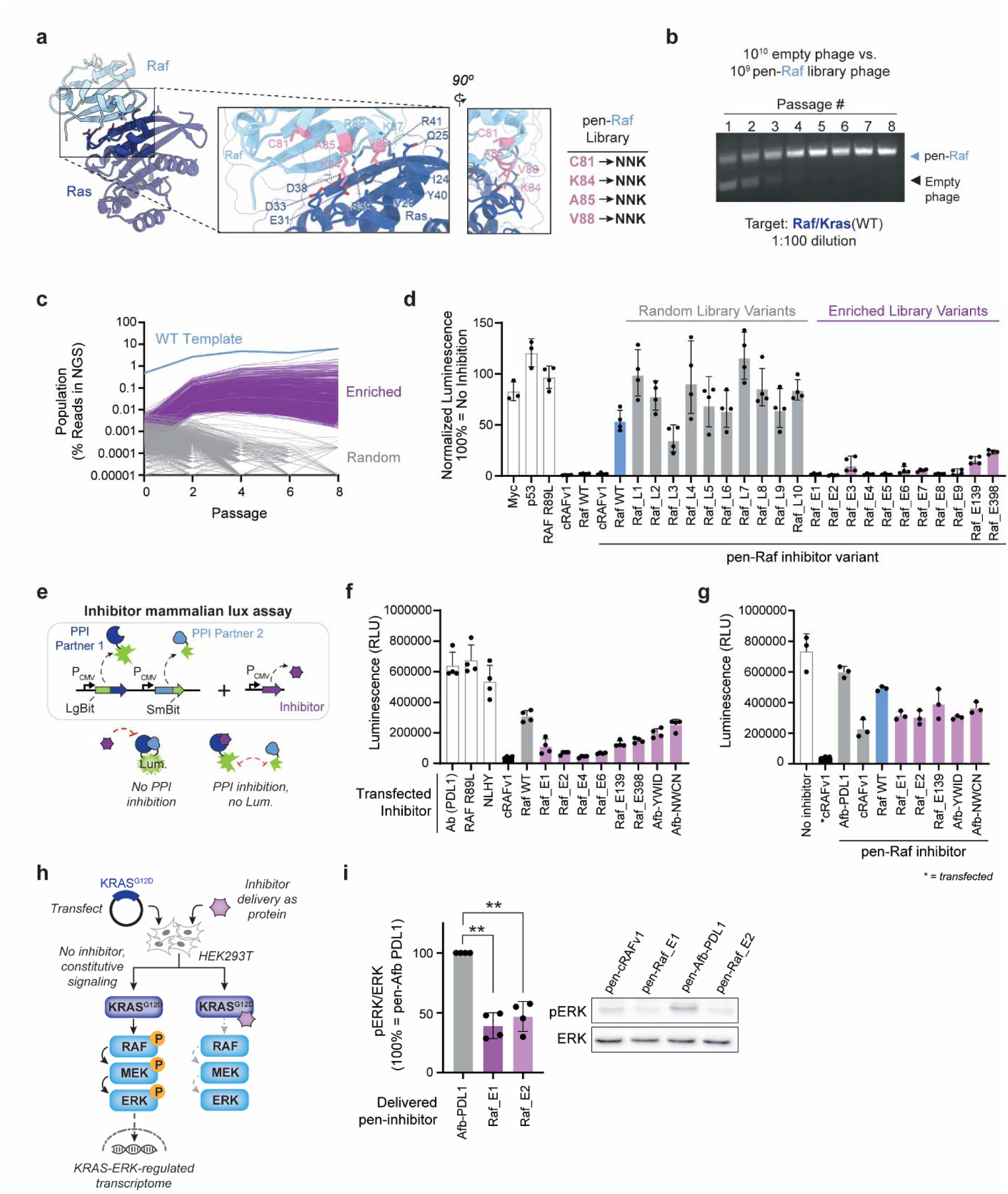
pen-Raf deep-mutational scanning of KRas-Raf inhibition. **a,** Structure of the KRas-Raf PPI (PDB: 6VJJ) and the residues selected for complete (NNK) randomization (pink). **b**, PANCS-Inhibitor pen-Raf 4xNNK library with 1:1 empty phage spike-in selection for KRas-Raf inhibition monitored by PCR. **c**, The 1000 most enriched library variants in Passage 8 (with spike-in) over Passages 0 (library) to 8 (purple curves) and 1000 random library members (grey; if NGS had 0 reads, population set to 0.00001% for plotting on log scale). The template pen-Raf sequence (not codable by NNK) used to construct the randomized library is shown as well (blue). **d**, PPI inhibitor E. coli luciferase assay (n = 4, error bars indicate SD) on KRas- Raf with 10 randomly isolated library variants (grey) and the 9 most enriched variants in passage 8 (with spike-in; purple) and two lesser enriched variants (the 139^th^ and the 398^th^). **e**, Mammalian PPI inhibitor split-Nanoluciferase complementation assay: two plasmid transfection into HEK293T (reporter plasmid with PPI (top left) and inhibitor expression plasmid (top right)). **f**, Transfection version of the mammalian PPI inhibitor split-Nanoluciferase complementation assay using Raf (not pen-Raf) variants (n =4, error bars indicate SD). **g**, Protein delivery of inhibitor (except where indicated as ‘transfected) version of the mammalian PPI inhibitor split-Nanoluciferase complementation assay using pen-Raf variants (n =4, error bars indicate SD). **h,** Depiction of pERK signaling assay for monitoring KRas-Raf inhibition phenotype with delivered pen-inhibitor protein. **i**, Quantification of pERK signaling via replicate WB of pERK signaling inhibition in HEK293T cells transfected with KRas^G12D^ expression plasmid to induce pERK signaling (representative example WB shown on right and replicates in **Fig. S13**); two-tailed *t*-test indicate that each inhibitor variant is significantly different than the non-inhibitor (p = 0.0015 and 0.0034, respectively). In **f**, **g**, and **i**, we have used an affibody that binds PDL1^57^ and does not bind KRas or Raf (**Fig. S12**) as our non-inhibiting control.

To measure the fitness of every variant, we sought to keep our library small enough so that we could fully sequence the relative copy number of every variant in our library for comparison to post-enrichment relative copy number. To ensure complete sequencing coverage, we limited ourselves to a 4-position NNK library (all amino acids possible; 1.05 million DNA variants, 160,000 protein variants). When considering this helix (L78 to G90), we decided to maintain intra-Raf contacts (hydrophobic core: L78, L82, M83, L86). The high affinity cRAFv1 has two mutations that alter the cap of this helix, rewriting several intra- and intermolecular hydrogen bonds to create an alternative network of interaction via V88R and R89H (**Fig. S9**). Therefore, we opted not to alter these residues in order to focus on epistatic interactions surrounding the A85 water pocket, rather than drastically altering the orientation of this helix via modifications to R89. Finally, instead of positions that were primarily solvent exposed (H79, D80, K87), we randomized C81, K84, A85, V88 (**Fig. 3a**), which are each oriented towards Ras. Finally, Nomura *et al.*^49^ recently reported that the cell-penetrating peptide “Pen” can be fused to the high affinity cRAFv1 (3.2 nM K_d_^30^), enabling direct delivery of this protein delivery into mammalian cells to inhibit the KRas-Raf interaction. However, they report that fusing Pen to the lower affinity wild-type Raf (130 nM K_d_)^50^ produces little to no inhibition. Thus, we included Pen in our library design such that we could optimize Raf-based inhibition in the context of the cell-penetrating Pen-Raf fusion. This DMS library provided us with a compelling proof-of-concept experiment: KRas-Raf inhibition is of high clinical importance, pen-Raf variants can be delivered to cells for inhibition, known mutations improve the affinity of the KRas-Raf interaction, and there is a clear gap in understanding the epistatic relationships within this helix.

We then ran PANCS-Inhibitors selections with this pen-Raf DMS library (**Fig. S10**) on our optimized KRas-Raf AP set for 8 passages using a 1/1000 dilution with and with-out a spike-in of empty phage (1:1 ratio of library to empty). As before, we monitored de-enrichment of empty phage in the spike-in selection by PCR, which suggested complete de-enrichment of empty phage by Passage 5 (**Fig. 3b**). We collected NGS (see methods) for the library and each even Passage (2, 4, 8) from the selection with and without the spike-in (**Fig. 3b**, **Table S2**). In the NGS of the library, we obtained 92% (964,398/1,048,576) coverage of unique DNA variants (including stop codon variants) and 99% (158,448/160,000) coverage of full-length protein variants (excluding variants where one or more of the randomized codons included a stop codon). We compared Passage 8 (P8) results from the two separate selections (with and without spike-in) and saw strong reproducibility between the selections (Pearson Correlation r = 0.82, p = <0.001; **Fig. S11**). Given this high correlation, we restricted our later analysis to the selection with the spike-in of empty phage. We detected 8,378 unique protein variants in P8; 3079 had enriched more than NNK-coded pen-Raf WT variants (**Fig. 3c**). We subcloned the top nine variants (**Table S1**) from P8 and ten random variants (**Fig. S10**) from the library and tested these for inhibition in our *E. coli* inhibitor luciferase assay (**Fig. 3d**). WT pen-Raf showed 63% inhibition while the high affinity pen-cRAFv1 and WT Raf (no pen) each showed 98% inhibition; random library variants showed 0-67% inhibition (22% mean inhibition); and eight of nine enriched variants showed at least 95% inhibition, five of which showed >98% inhibition, and one variant showed >99% inhibition (equivalent to cRAFv1 without pen). We tested a subset of the top point mutants and top variants as pen-Raf and Raf only in our *E. coli* LuxAB soluble expression (RNAP_N,wt_ recombines independent of binding, **Fig. S12**), binder, and inhibitor assays. These results suggest that: i) these mutations improve binding in pen-Raf and Raf; ii) when corrected for expression, binding and inhibition are highly correlated (Pearson Correlation, r = 0.98); and iii) mutations that improved affinity often reduced soluble expression. We tested a subset of our top variants as transfected Raf variants or as purified His-tagged pen-Raf variants in HEK293T cells using a split nano luciferase assay (**Fig. 3e**). When transfected, these Raf variants showed improved inhibition of the KRas-Raf interaction compared to WT Raf (**Fig. 3f**). Similarly, when delivered as purified protein, these variants performed much better than WT pen-Raf (**Fig. 3g**). Finally, we stimulated MAPK signaling in HEK293T cells by transfecting KRas^G12D^, which constitutively activates MAPK signaling, and monitored functional disruption of the endogenous KRas^G12D^/Raf PPI by monitoring ERK phosphorylation. When delivered as proteins, our evolved pen-Raf variants showed robust inhibition, unlike pen-Raf WT (**Fig. 3h**, **Fig S13**). Additionally, we demonstrate that the Raf binding affibodies (<10 nM K_d_) that inhibit the KRas-Raf interaction in our *E. coli* inhibitor luciferase assay (**Fig. 1g**) also function in this set of mammalian cell experiments (**Fig. 3f-h**) as a Raf targeting means to inhibit the KRas-Raf interaction. In sum, these results suggest that PANCS-Inhibitors performed as expected at the technical level and enriched highly inhibitory pen-Raf variants.

### Ruggedness of the pen-Raf fitness landscape

We sought to characterize the sequence-function, or fitness, landscape of the pen-Raf interaction to understand how the landscape could be navigated rationally (i.e., a Greedy algorithm) or evolutionarily (step-wise mutations from WT). Rather than raw percentage of the population at the end of the selection as the proxy for fitness (**Fig. 3c**), we sought to use the slope of each variant from its population in the library to a later point in the selection as this is highly predictive (Pearson’s Correlation, r = 0.86, p = <0.0001) and corrects for initial variation in population, unlike enrichment **(Fig S14)**. We compared possible endpoints for the slope calculation and decided to use P4 which balances the number of sequences quantified in the passage vs. alignment with later passages (**Fig S15**). Slopes ranged from 389 to 0.02 (**Fig. 4a**), with 2229 variants with slopes better than WT (slope of 3.6). We then normalized slopes to WT=1. We visualized this fitness landscape (**Fig. S16** for organization schematic) and observed a landscape with many local maxima (**Fig. 4b**). We performed hierarchical clustering of the top 1000 variants resulting in 118 distinct clusters (**Fig. 4c**, **Table S3**).^58^. We sorted cluster by which positions were constant (**Table S4**) allowed us to determine: i) how frequently each position was converged (**Fig S17**), and ii) how frequently two positions converge together (**Fig. 4d**). Position 85 and 88 appear tightly connected, while position 81 is both infrequently constant in a cluster and only loosely connected to these other positions.

**Figure 4:**
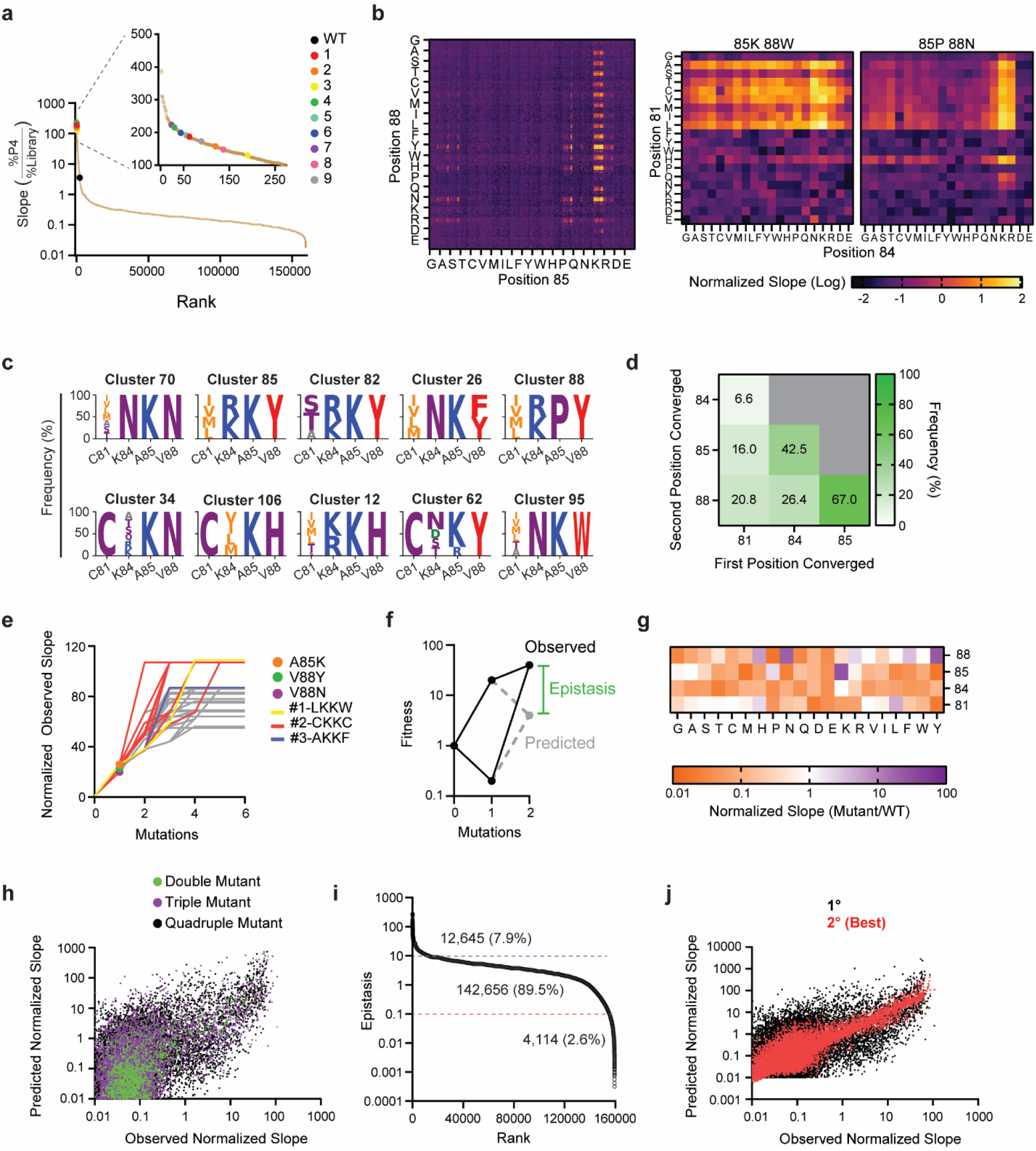
Navigating the pen-Raf fitness landscape. **a,** Slopes for every protein variant observed in the library and/or passage 4 NGS; slopes are calculated as the relative abundance of the variant in Passage 4 divided by the relative abundance of the variant in the initial library (any variant not observed in either NGS data is set to a relative abundance of 0.001%). The library coded WT as well as the top nine variants by enrichment (Fig. 3c) are indicated in the legend/inset. **b**, The fitness of every variant in the data set is shown on this 400x400 heatmap organized as smaller squares of Position 81 by Position 84 (inset examples to the right) within a larger set of squares for Position 88 by Position 85 (**Fig. S16** has a depiction of this organization). **c**, The sequence logos for the top ten clusters by average variant fitness (**Table S3**). **d**, Frequency of co-convergence between two positions within the clusters (**Table S4**). **e**, Changes in variant normalized slope (fitness) as the WT variant evolves with the constraint that the fitness must increase substantially (at least +6 in slope). Each line depicts a different evolutionary path. The three possible single mutations are indicated as well as the three most fit stable evolutionary endpoints. **f**, Depiction of epistasis when single mutant fitness data is used to predict double mutant fitness. **g**, The normalized slope for all single mutations from WT. **h**, The epistasis score for all variants using the 0.01 normalized slope floor for predicted fitness using single mutation fitness. Lines are indicated at 10x below and above WT (=1). **i**, The observed vs. predicted fitness for all double, triple, and quadruple mutants. **j**, The epistasis score for all triple and quadruple mutants using predicted fitness values based only on the single mutant data is shown in black (**Fig. 4f**) while the “best” of the predicted fitness values based on both single and double mutant data is shown in red (**Fig. S21**).

Given the observed ruggedness, we were interested in how well evolution or rational optimization algorithms might navigate this fitness landscape. To allow for easy navigation of this fitness landscape, we also generated an interactive webpage (https://bryandickinson-create.github.io/dms-fitness-explorer/). We followed three step-wise algorithms commonly used in affinity maturation (**Table S5**). First, we used a Greedy algorithm for each possible pathway to specify each position with the best available amino acid (based on the average value of all variants possible). Second, we start with WT and make a single mutation for each possible pathway to specify each position with the best available point mutant. Third, we start with WT and identify every evolutionary pathway that substantially (at least >+6 in slope) improves fitness (**Fig. 4e**) for 35 total pathways. These algorithms result in average fitness outcomes of 78 (+/-21 STD), 81 (+/-24 STD), and 87 (+/-17 STD) respectively, with each having limited access to either the best variant (mutations from WT) or the second best variant (Greedy algorithm) with all pathways substantially improving over WT, but with each approach yielding suboptimal (> +20 in slope is possible) in 75, 70, and 67% of pathways, respectively. These results demonstrate the difficulties presented in navigating this sequence landscape to reach the maximal fitness using step-wise evolutionary or algorithmic processes.

### Predictability of multi-mutant fitness from single mutations

Several recent efforts have used single-^59,60^ or double-mutant^43,61,62^ fitness data to improve protein engineering and directed evolution. While much more difficult to obtain, double mutant fitness data includes information on epistatic connections between two single point mutants and their non-additive effects on fitness (**Fig. 4f**). Such effects are potentially quite important in protein engineering,^43,44^ but often sparse data identifies only a fraction of potential epistatic connections. Given our nearly complete mapping of fitness within this 4-position evolutionary landscape, we sought to quantify how epistasis impacts the fitness landscape and its navigability. We have quantification of every point mutant variant in both the library and P4 NGS with normalized slopes ranging from 26.5 (A85K) to 0.02 (A85M) (**Fig. 4g**). Based on how each individual mutation alters fitness, we predicted the fitness and epistasis of all double, triple, and quadruple mutations (**Fig. S19**). However, many individual mutations confer loss of function (slopes <1) resulting in predictions with very low slopes (<0.001), causing a distortion in the relationship between predicted and observed slopes (Pearson Correlation, r = 0.282) and several wildly high (>10^6^) epistasis scores. As the lowest observed slope was 0.02 (this lower limit was set primarily by our NGS coverage depth), we re-calculated the predictions setting a lower limit on predicted slope of 0.01 (**Fig. 4h**; Pearson’s Correlation, r= 0.540, p = <0.0001). The predictions were much better for double mutants (r = 0.702) than for triple (r = 0.553) or quadruple (r = 0.525) mutants, reflecting the compounding effects of multiple epistatic interactions. We observed epistasis values (**Fig. 4i**) ranging from 272 to 0.003 with a median epistasis score of 3.7. Only 880 variants were predicted to have a slope >10 and only 106 with a slope >100. Active variants had epistasis scores centered between 1 and 10 (**Fig. S20a**), and the range of epistasis scores showed minimal differences by number of mutations (**Fig. S20b**). Next, we used both single and double mutation observed fitness in order to predict fitness of the triple and quadruple mutants; however, this did not produce a single prediction as with the point mutants, but rather a set of predictions (**Fig. S21**; see Methods) that can be characterized as a single value using the median value or the best value (closest to Observed). With single mutant based predictions (1°), the average epistasis score is 7.26; using the median epistasis score with doubles and singles (2° Median), the average is 3.96, but if only the best 2° value (2° Best) is considered, the average is 1.14; **Fig. S21**) and the correlation between observed and predicted slope improves dramatically (**Fig. 4j; Fig. S21**, 1° r = 0.543, 2° Median r = 0.610, and 2° Best r = 0.902). We can see that epistasis inserts significant error into the 1° predictions of fitness and that if we know which 2° prediction was “best” we could eliminate nearly all of this error; however, if we only had the single and double mutant data, we would only know the set (and median) of 2° predictions and this significantly impacts our ability to identify optimal variants. If we just consider the top 10 predicted variants for each set (1° or 2°), we have an average observed slope of 42 (1°) and 54 (2°) and highest fitness variant with slopes of 65 (1°) and 86 (2°); therefore, the predictivity is high enough to be useful, but insufficient for maximizing variant fitness (which can reach slopes of 109). The “best” prediction from the double and single mutants nearly captures all the remaining variance between observed and predicted fitness suggesting that second-order epistasis (which arises based on interactions between three positions) could only have minor impacts in the higher order fitness landscape. Therefore, further refinements to isolate the “dominant” epistatic interaction(s) (rather than using the median predictions) are needed to improve the accuracy of prediction of higher order mutant fitness.

### PANCS-Inhibitors selections for *de novo* mini-protein PPI inhibitors

Lastly, we sought to use PANCS-Inhibitors to select for *de novo* PPI inhibitors. We constructed a randomized affibody library (∼10^8^ variants, **Fig. S22**) in our inhibitor phage design (**Fig. 5a**, analogous to our previously reported randomized affibody library in our binder phage). We then performed an 8-passsage PANCS-Inhibitor selection with this library and our optimized selection strains for Myc-Max and Mdm2-p53. We used spiked-in empty phage to monitor the selection progress. Both selections resulted in rapid de-enrichment of the empty spike-in phage and increased intensity of the affibody library band (**Fig. 5b**), consistent with the strong performance of these strains in our mock selections (**Fig. 2f**) and suggesting that the remaining affibody variants were likely inhibitors. We performed NGS to observe the distribution of enriched variants. As expected based on our previous PANCS-Binders results,^41^ one variant dominated the Myc-Max selections (**Fig. 5c**, SGAD) while much less convergence was observed in the Mdm2-p53 selection (**Fig. 5c**), consistent with the minimal Mdm2 binding motif (which competes with p53 binding) having roughly a 1% frequency in the library design. We subcloned the top variant from each of these selections and confirmed that they inhibit their respective PPIs in our *E. coli* inhibition luciferase assay (**Fig. 5d**). Finally, we used our *E. coli* binding luciferase assay with each target protein to determine which protein the inhibitor bound to (**Fig. 5e**): TWAA bound to Mdm2 (as expected) and SGAD bound to Myc. AlphaFold3 fails to predict the correct binding pattern for SGAD and TWAA, but in the predicted structures for Myc-SGAD and Mdm2-TWAA, the predicted binding orientation should inhibit Myc-Max and Mdm2-p53 respectively (**Fig. S23**). We determined the affinity of these binding interactions by SPR (**Fig. 5d, Fig. S24**) suggesting moderate affinity interactions for both (consistent with the binding affinities we had previously identified from 10^8^ affibody library selections). Finally, we confirmed that these variants inhibit their respective PPI in mammalian cells using a split Nano luciferase reporter (**Fig. 5f**) and via inhibition of the endogenous interactions: for Mdm2-p53, transient transfection of TWAA leads to activation of p53 transcription including overexpression of Mdm2 and p21 (**Fig. 5g**), and for Myc-Max, we transient transfection of SGAD (fused with a nuclear localization signal (NLS)) leads to loss of Max pull down in co-immunoprecipitation when endogenous Myc is pulled down using α-cMyc (**Fig. 5h**). These results demonstrate that PANCS-Inhibitor can be used to rapidly isolate *de novo* PPI inhibitors that function in the desired context.

**Fig. 5:**
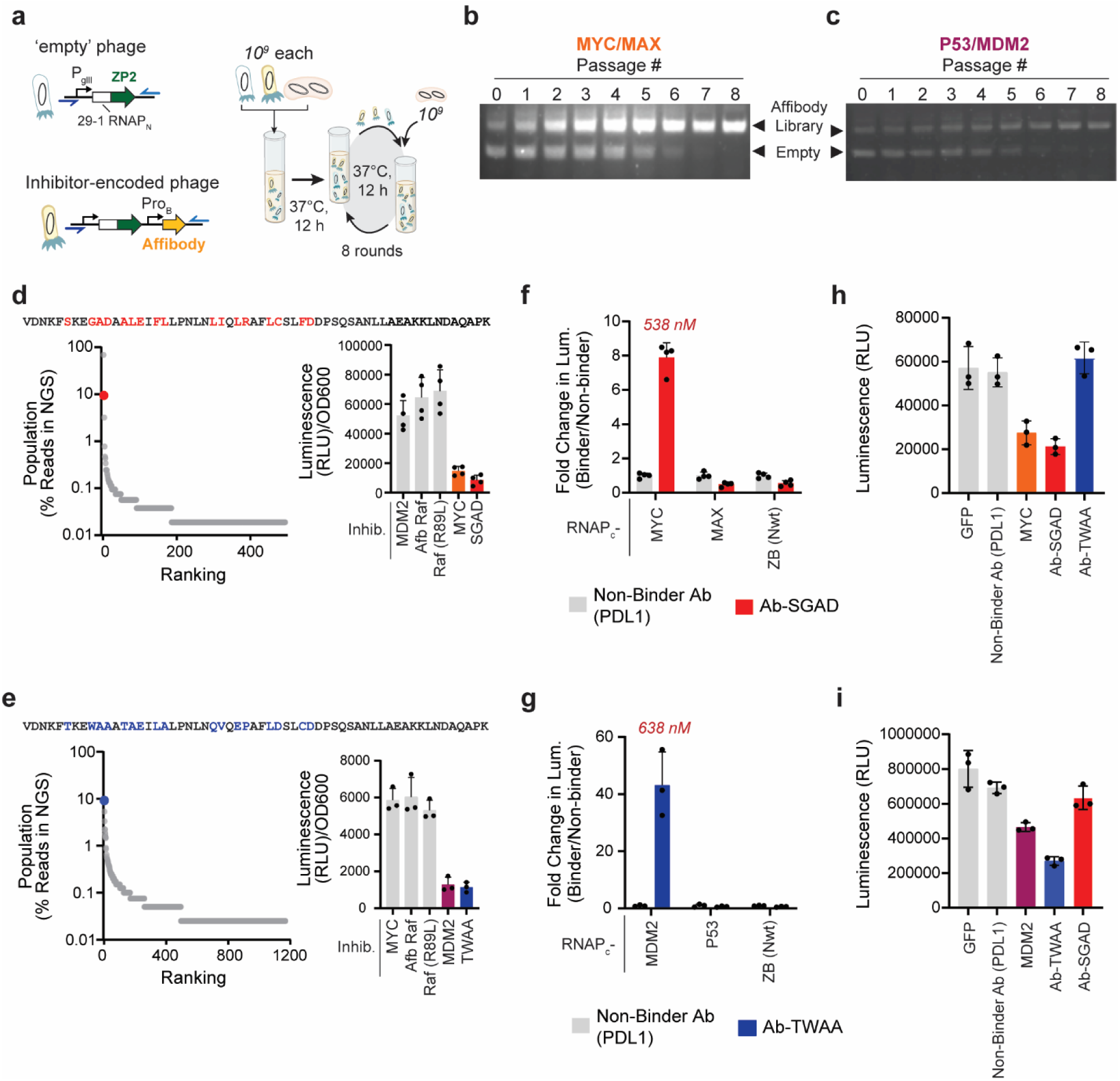
D*e novo* selection of PPI inhibitors. **a,** Using an affibody library (**Fig. S22**) for *de novo* inhibitor discovery. **b-c**, PANCS-Inhibitors affibody selection for Myc-Max (**b**) and Mdm2-p53 (**c**) inhibition monitored by PCR. **d**, Sequence prevalence plot and *E. coli* lux inhibition for the Myc-Max selection (Lux: n=4, error bars indicate SD). **e**, Sequence prevalence plot and *E. coli* lux inhibition for the Mdm2-p53 selection (Lux: n=3, error bars indicate SD). **f**, *E. coli* binding luciferase data for Affibody SGAD with Myc and Max and for expression luciferase with ZB (RNAP_Nwt_) (n=4, error bars indicate SD). The K_d_ of the Myc-SGAD interaction, as measured by SPR (**Fig. S24**) is listed above the luciferase signal. **g**, *E. coli* binding luciferase data for Affibody TWAA with Mdm2 and p53 and for expression luciferase with ZB (RNAP_Nwt_) (n=4, error bars indicate SD). The K_d_ of the Mdm2-TWAA interaction, as measured by SPR (**Fig. S24**) is listed above the luciferase signal. **h**, Inhibition of Myc-Max PPI in the mammalian split Nano-Luciferase assay when inhibitors are delivered by transient transfection(Lux: n=3, error bars indicate SD). **i**, Inhibition of Mdm2-p53 PPI in the mammalian split Nano-Luciferase assay when inhibitors are delivered by transient transfection (Lux: n=3, error bars indicate SD).

## Discussion

Typically, PPI inhibitors are discovered by first performing selections to identify binders to one of the proteins of the desired PPI target and then testing individual binders for the ability to inhibit the PPI. This process, even when successful, is costly and inefficient. Identified binders interact with hot-spots (**Fig. 1a**) on the protein target, and binding a hot-spot does not necessarily entail inhibition. Therefore, a selection for binding is likely to produce a mixture of binding variants that one must just sort through to find the ones that inhibit. Additionally, binders that inhibit may not be great binders; for example, binders that induce conformational changes in one of the target proteins, likely to be allosteric inhibitors, are unlikely to be the highest affinity binders in binder selections as these binding interactions must pay the cost of altering the target’s conformation. Therefore, we propose that direct selection of variants for their function as inhibitors of a PPI, rather than as binders to one of the PPI targets, is likely to isolate only binders that inhibit, abrogating the need to screen for variants, and allowing enrichment of lower affinity binders that actually have the desired inhibitory function. To this end, we developed PANCS-Inhibitors as a rapid, high fidelity means to directly select for PPI-inhibiting variants.

In addition to the direct selection of PPI inhibition, PANCS-Inhibitors has several clear strengths. First, as with most selection platforms, we recognize that the antigen used (purified protein in display methods or expressed proteins in the *E. coli* cytoplasm used here) may not accurately recapitulate the target protein in the desired context; however, because PANCS-Inhibitors relies on the detection of a known PPI between the two target antigens, we can be confident that the PPI in *E. coli* compares reasonably to PPI in mammalian cells as the observation of the interaction entails at minimum proper folding of the two target proteins. We would expect, however, that some PPIs simply will not be detectable in our system (as currently designed) such as PPIs involving multi-pass transmembrane proteins or PPIs dependent on a post-translational modification. Second, like PANCS-Binders, these selections utilize gradient selection pressure and activity dependent replication, and therefore, we were unsurprised that we did not identify any false-positive enriched variants. Third, the selection biosensor is tunable, enabling detection of a wide range of PPI affinities with a clear strategy for optimization: vary the expression on AP2, and if necessary, lower the expression of AP1. Fourth, while we did not utilize the off-target PPI that must not be disrupted in our biosensor (ZA and ZB) for counterselection, this off-target interaction could be used to select for inhibitors that are specific for one PPI over a closely related PPI (i.e., KRas-Raf inhibition but not KRas-Sos1 inhibition) as we have shown in PACE^63^ and PANCS-Binders.^45^ This aspect of the PANCS-Inhibitors could allow for the development of tools to precisely dissect the importance of a PPI where simple genetic knock-out experiments cannot.

In summary, this work presents a powerful new platform for the discovery of PPI inhibitors. PPIs are critical therapeutic targets. Additionally, basic research surrounding PPIs suffer confounding effects when explored via genetic knock-out as most proteins participate in 3+ PPIs. Current approaches to identifying PPI inhibitors generally takes months to years, if successful, generally via a binder discovery campaign followed by screening of binding variants for whether they act as inhibitors. This process is severely limited by the nature of the binder discovery process which favors only binding affinity – thus failing to identify binders to lower affinity sites or lower affinity binders that induce alternative target conformations (allosteric inhibitors). PANCS-Inhibitors offers direct selection for the desired function – disruption of a target PPI – in a manner agnostic to mechanism. We have demonstrated that selection outcomes differ when a highly similar library is used to identify binders vs. identify inhibitors, and this specifically allowed us to identify a Myc-Max inhibitor where we previously had only identified non-inhibitory binders to these proteins. Finally, we demonstrate that PANCS-Inhibitors can be used for deep mutational scanning and affinity maturation of known inhibitors. These proof-of-concept studies have yielded a suite of new molecules that inhibit the Myc-Max, Mdm2-p53, and KRas-Raf oncogenic interactions that function in the mammalian cell context.

## SUPPORTING INFORMATION

Details of bacterial strains, plasmids, primers, additional data figures and tables.

## Data Availability

Links to electronic vector maps are included in Supplementary Information. All physical vectors will be made available upon reasonable request. Source data are provided with paper.

## Acknowledgements

This work was supported by the National Institute of General Medical Sciences (GM119840 to B.C.D and F32GM147968 to M.J.S) and then National Cancer Institute (P30CA014599) of the National Institutes of Health. Critical early support for this project was provided by the National Cancer Institute IMAT program (R21 CA217754). B.C.D. is a Biohub Investigator. V.C.X. was supported by a National Science Foundation Graduate Research Fellowship (DGE-1746045). We thank S. Ahmadiantehrani for assistance with preparing this paper.

## Contributions

Conceptualization: V.C.X., M.J.S. and B.C.D. Methodology: M.J.S. and V.C.X. Investigation: S.L, M.J.S., V.C.X., and J.A.P. Writing – original draft: M.J.S., V.C.X., and B.C.D. Writing – reviewing and editing: J.A.P. and S.L. Supervision: B.C.D.

