## Supplementary Information for "PANCS-Inhibitors: A rapid method to directly select for protein-protein interaction inhibitors"

**TABLE OF CONTENTS**

**Methods**

**Supplementary Fig. 1: KRas PPIs and *de novo* binders**

**Supplementary Table 1: Sequences of PPI partners and previously reported binders and inhibitors**

**Supplementary Fig. 2: Additional controls in optimized PPI inhibition E. coli luxAB assay**

**Supplementary Fig. 3: AF3 predicted binder-target complexes aligned with native target PPIs**

**Supplementary Fig. 4: *In vivo* phage-assisted directed evolution/selection platforms (in general)**

**Supplementary Fig. 5: PANCS-Binder selection platform**

**Supplementary Note 1: Unsuccessful PANCS-Inhibitor biosensor designs**

**Supplementary Fig. 6: Optimization on- and off-target inhibitor phage replication rates**

**Supplementary Fig. 7: Additional Mock Selection results**

**Supplementary Fig. 8: Re-wired interaction network for cRAFv1 – RAS PPI**

**Supplementary Fig. 9: RAS-Raf PPI water pocket near A85 of Raf**

**Supplementary Fig. 10: pen-Raf Library**

**Supplementary Table 2: DMS NGS Results**

**Supplementary Fig. 11: Comparison of selections with and without spike-in**

**Supplementary Fig. 12: Soluble Expression, Binder, and Inhibitor Lux Comparison**

**Supplementary Fig. 13: Replicate pERK Western Blots**

**Supplementary Fig. 14: Comparing enrichment and slope to measured inhibition**

**Supplementary Fig. 15: Comparing slopes using different endpoint measures**

**Supplementary Fig. 16: Diagram of how the fitness landscape is organized**

**Supplementary Table 3: Clustering of top 1000 DMS variants**

**Supplementary Table 4: Analysis of DMS Clustering**

**Supplementary Fig. 17: Breakdown of convergence pattern within clusters**

**Supplementary Table 5: Step-wise Algorithmic exploration of the pen-Raf fitness landscape**

**Supplementary Fig. 18: Step-wise Algorithmic exploration of the pen-Raf fitness landscape**

**Supplementary Fig. 19: Observed vs Predicted fitness (from single mutants) without lower bound**

**Supplementary Fig. 20: Epistasis using single mutants with a 0.01 Predicted Slope lower bound**

**Supplementary Fig. 21: Predicting Fitness with single and double mutant data**

**Supplementary Fig. 22: PANCS-Inhibitor Affibody Library Construction**

**Supplementary Fig. 23: AlphaFold Prediction of de novo inhibitors**

**Supplementary Fig. 24: SPR of *de novo* inhibitors**

**Supplementary Table 6: Plasmids utilized in this study**

**Supplementary Table 7: Primers used in this study**

**Supplementary Fig. 25: Full WB Images**

**Supplementary Information References**

### Methods

#### Cloning and Bacterial Strain Handling

All plasmids and phage (**Supplementary Table 6)** were cloned by Gibson Assembly (GA) of PCR fragments generated using Q5 DNA polymerase (NEB). All primers (**Supplementary Table 7**) were ordered from IDT. Three *E. coli* strains were utilized in this work: DH10β (Thermo Fisher EC0113), BL21(DE3) (Thermo Fisher EC0114), and S1030 (Addgene 105063). For plasmids, GA mixtures were transformed into chemically competent DH10β *E. coli* and after a 1 h outgrowth in 2xYT media, were plated on antibiotic selective agar plates to isolate individual clones. For phage, GA mixtures were transformed into chemically competent S1030-1059 *E. coli*, and after a 2 h outgrowth, a plaque assay was performed to isolate individual phage clones. All plasmids and phage were confirmed by Sanger sequencing. Plasmid maps with annotations of key features are available in **Supplementary Table 6**. For constructing selection (AP1/AP2) and *E. coli* luciferase (inhibitor/N-PPI_1/PPI_2-C-lux) strains, S1030 *E. coli* was made chemically competent and then single or double transformations were used (and then repeated as needed until all plasmids were incorporated). *E. coli* strains were grown on agar plates static at 37 °C or in solution at 37 °C with 200 rpm shaking with Luria Broth (LB) supplemented with the appropriate antibiotic unless otherwise indicated. Antibiotics were used at standard concentrations: kanamycin (40 ug/mL), chloramphenicol (33 ug/mL), and carbenicillin (100 ug/mL).

#### Plaque Assays

Activity independent plaque assays were used to determine the phage titer via plaque counting. For activity independent plaque assays, an S1030-1059 *E. coli* culture (1059 plasmid encodes *gIII* expressed from the phage shock promoter to produce *gIII* after phage infection), was grown to stationary phase in LB with carbenicillin, subcultured 1:10 in fresh LB with antibiotic to an OD600 of 0.4-0.6, and then used as the *E. coli* strain in the plaque assay. For the plaque assay, an initial dilution of the stock can be added based on the expected titer, but generally, 2 µL of a phage stock (or diluted stock) was added to 100 µL of subculture, mixed, and then serially diluted (2 µL into 100 µL) to create four dilutions. 750 µL of 50 °C top agar (7 g/L agar, 25 g/L LB) was added to each dilution, which was then transferred in its entirety to one quadrant of a bottom agar plate (15 g/L agar, 15 g/L LB). After 10-16 h of incubation at 37 °C, plaques became visible and were counted in the quadrant with 10-200 plaque forming units (PFU).

#### Phage Amplification Rates

To determine phage amplification rate, the titer of a phage stock was determined using an activity independent plaque assay. Based on this titer, a 1000x stock was prepared (i.e. for $10^3$ PFU, a $10^6$ PFU/µL stock). The activity dependent strain (AP1/AP2) was grown to stationary phase in LB with carbenicillin and kanamycin, subcultured 1:10 in fresh LB with antibiotic to an OD600 of 0.4-0.6. The desired PFU (1,000 unless otherwise stated) was added to 1 mL of this subculture and then incubated at 37 °C with shaking for 12 h. The cells were pelleted and the cell-free supernatant collected for activity independent plaque assay to determine the titer at the end of the amplification assay. The endpoint titer was divided by the starting titer to determine the amplification rate.

### PANCS-Inhibitor

Protocol used in mock library selections. For each passage, a selection strain (AP1/AP2) was grown to stationary phase in LB with carbenicillin and kanamycin, 10 mL overnight culture was added to fresh LB with carbenicillin and kanamycin prior to adding phage. All PANCS performed in this work (unless otherwise noted) were performed in 1 mL cultures in a deep 96-well plate. For passage 1, stock phage were added to the subculture and incubated at 37 °C with shaking (200 rpm) for 10-14 h, then centrifuged to pellet the cells and collect the cell-free supernatant (referred to as passage 1 phage). For subsequent passages, some fraction of the cell-free supernatant from the prior passage was added to the subculture and incubated at 37 °C with shaking (200 rpm) for 12 h, then centrifuged to pellet the cells and collect the cell-free supernatant (referred to as passage # phage). Titers of each passage or just the final passage were then determined using activity independent plaque assays.

Protocol used in library selections. For each passage, a selection strain (AP1/AP2) was grown to stationary phase in LB with carbenicillin and kanamycin, subcultured 1:10 in fresh LB with carbenicillin and kanamycin to an OD600 of 0.4-0.6 prior to adding phage. All PANCS performed in this work (unless otherwise noted) were performed in 1 mL cultures in a deep 96-well plate. For passage 1, stock phage were added to the subculture and incubated at 37 °C with shaking (200 rpm) for 10-14 h, then centrifuged to pellet the cells and collect the cell-free supernatant (referred to as passage 1 phage). For subsequent passages, some fraction of the cell-free supernatant from the prior passage was added to the subculture and incubated at 37 °C with shaking (200 rpm) for 12 h, then centrifuged to pellet the cells and collect the cell-free supernatant (referred to as passage # phage). Titers of each passage or just the final passage were then determined using activity independent plaque assays.

### PCR to monitor empty phage de-enrichment

Selections were monitored by Phusion PCR (NEB, 50 uL reactions) using 1 µL of phage from selections that contained an empty phage spike-in control using the following primers: BR-76 and JD-1060, which result in a 735 bp fragment when used on just the empty phage and an at least 931 kb fragment when used on inhibitor-encoding phage (exact length depends on the inhibitor). The limit of detection after 25 cycles was ~$10^6$ PFU/mL.

### Split-RNAP *E. coli* Luciferase Assays

Binding. For the *E. coli* luciferase binding assays, one vector contained the previously-evolved PPI-dependent N-terminal half of RNAP[1] fused to one protein, and another vector encoded both the C-terminal half of CGG RNAP fused to one protein and the CGG promoter-driven luciferase reporter.[2] After transformation to assemble the correct set of plasmids in S1030 cells, colonies (3-4 biological replicates per condition) were picked for each binder and non-binder and each strain was inoculated (1 mL of LB with kanamycin and carbenicillin) and grown at 37 °C with shaking (200 rpm) for 12-16 h. Strains were then subcultured 1:10 in LB with kanamycin and carbenicillin in white sided, clear bottom 96-well assay plates (Corning 3610). Plates were incubated at 37 °C with shaking (200 rpm) for 3.5 h prior to reading the OD600 and luminescence signal on a BioTek Synergy Neo2 plate reader. Luminescence signal is first divided by the OD600 to normalize luminescence to cell growth. Then the luminescence/OD600 is normalized for all strains with the same target-RNAP$_C$ luciferase plasmid: all luminescence/OD600 are divided by the non-binder signal (non-binders set equal to 1). Data was analyzed and plotted in GraphPad Prism10. Due to differences in expression levels of each target and differences in how binding impacts expression level of a target, we do not believe direct comparisons in fold change over non-binder can be made across different targets.

_Inhibition_. The *E. coli* luciferase inhibitor assays contained both vectors described for binding assays, as well as a vector containing an isopropyl -D-1thiogalactopyranoside (IPTG)-inducible inhibitor. Similarly, the trimolecular complex, i.e., inducer, assay (**Fig. 2b**) likewise contained both vectors described above as well as a vector containing an IPTG-inducible expression cassette with various protein fusions.[3] Assays were preformed identically, but with chloramphenicol added to the overnight and assay culture media and with IPTG (1 mM inhibitor or 0.1 mM inducer, both from 1 M stocks) added to assay media.

_Expression._ Identical to the binding *E. coli* luciferase assay, but instead of using the PPI-dependent, evolved RNAP N-terminal fused to the testing protein, we used the WT RNAP N-terminal which assembles independently of a PPI. As a result, an off-target-RNAP$_C$ fusion protein (the short zipper peptide, ZB) was used as well.[2]

### Library Construction

Libraries (four NNK pen-Raf and Affibody[2]) were constructed as we previously reported[2], but with the PPI inhibitor SP design as template phage (pen-Raf and Affibody (PDL1) respectively). Randomization was installed into a template phage using primers with degenerate codons (IDT; see **Supplementary Table 7**). Briefly, PCR was then scaled up to produce 20-100 μg of PCR product which was concentrated (Wizard Kit Promega) and then digested (~20-50 μg of PCR product in 300-400 μL) using DpnI and restriction enzyme (pen-Raf – PstI-HF; affibody - NheI-HF; NEB), agarose gel purified (Zymo Gel Extraction kit), and then ligated with T4 DNA Ligase (NEB). Ligated products were then electroporated into 1059 *E. coli* cells. The cells were then recovered in 50 mL of 37 °C SOC media and incubated for 2 h at 37 °C with shaking – samples were collected throughout this time for determining the titer by plaque assay. At 2 h, the cells were pelleted and the cell-free supernatant was collected. 1030-1059 (activity independent replication strain) was grown overnight and then subcultured 1:10 to and OD600 of 0.6 at 37 °C with shaking (200 rpm). The phage (cell-free supernatant) was then amplified by adding the phage to this subculture for 8-10 h. At the conclusion of this outgrowth, cells were pelleted and the cell-free supernatant was sterile filtered to create the final library stock (titer determined by activity independent plaque assay).

### Next-Seq of Pen-Raf Library and Selection

To prepare Illumina sequencing libraries, we followed our previously published protocol.[4] Briefly, phage stocks were boiled at 95°C for 5 minutes, and 2 μL of each sample were used as template for the first round of PCR amplifications using 16 cycles. We used a combination of four different forward and four different reverse primers (**Supplementary Table 7**) which add a random barcode with varied length (6-9 N's) to each sample for library quality assessment. All 16 combinations of forward and reverse primers were used for each phage sample. The 16 PCR products were pooled and 1 μL was used for the second-round PCR to add a unique index for each sample and the Illumina adaptor as shown in the table here (primers in **Supplementary Table 7**). These PCRs were purified using the Zymo DNA clean and concentrator kit (catalog number D4013). Concentration was measured using a Qubit 4 Fluorometer and length assessed using an Agilent TapeStation 4200. DNA samples were mixed to maximize diversity in earlier passages/library, diluted and spiked with PhiX following the Illumina NextSeq System Denature and Dilute Libraries Guide (for use with a P1 300 cycle kit) prior to running on a NextSeq 2000.

| *Round 2 PCR and Relative Amount* | | |
|---|---|---|
| **Samples** | **Primers** | **%** |
| Library | D701/D501 | 40 |
| P2 | D702/D501 | 16 |
| P4 | D703/D501 | 8 |
| P6 | D704/D501 | 4 |
| P8 | D705/D501 | 2 |
| Spiked-P2 | D706/D501 | 16 |
| Spiked-P4 | D707/D501 | 8 |
| Spiked-P6 | D708/D501 | 4 |
| Spiked-P8 | D709/D501 | 2 |

### Calculations: Enrichment, Slope, Epistasis

*Enrichment ($E_i$):* $E_i = 100^* R_i / R_T$

Enrichment for variant i ($E_i$) as a percentage of the population in a sample: NGS reads for a variant ($R_i$) divided by the total NGS reads ($R_T$) for that sample (e.g. Library or Passage 2 with spike-in).

*Slope ($S_{i,\#}$):* $S_{i,\#} = E_{i,\#} / E_{i,L}$

The slope of a variant in a passage ($S_{i,\#}$) is the $E_i$ in a passage ($E_{i,\#}$) divided by the $E_i$ in the library ($E_{i,L}$).

*Normalized Slope ($N_{i,\#}$):* $N_{i,\#} = S_{i,\#} / S_{WT,\#}$

The normalized slope is the slope of a variant, $S_{i,\#}$, divided by the wild-type variant slope, $S_{WT,\#}$.

*1° Predicted Slope ($P_{1,ABCD}$):* $P_{1,ABCD} = N_A{}^*N_B{}^*N_C{}^*N_D$

The first-degree predicted slope, based only on single mutation data, is the product of the $N_{i,\#}$ for each single mutation that composes the variant. For example, for variant ABCD, the predicted slope would be $N_A{}^*N_B{}^*N_C{}^*N_D$. If A, B, C, or D are WT, then that normalized slope = 1 and does not alter the predicted slope. We argue that these values should be multiplied based on predicting the Slope based on the additivity of the change in free energy of the individual mutations:

$$\Delta\Delta G_{i,o} = \Delta G_{i,o} - \Delta G_{WT} \quad \text{and} \quad \Delta G_{i,o} = RT^*\ln(K_d) \quad \text{and} \quad K_d \propto 1/S_{i,o}$$

Therefore $-\Delta\Delta G_{i,o} = RT^*\ln(S_{i,o}) - RT^*\ln(S_{WT,o}) = RT^*\ln(S_{i,o}/S_{WT,o}) = RT^*\ln(N_{i,o})$

Therefore $-\Delta\Delta G_{ABCD,o} = RT^*\ln(N_{A,o}) + RT^*\ln(N_{B,o}) + RT^*\ln(N_{C,o}) + RT^*\ln(N_{D,o}) = RT^*\ln(N_A{}^*N_B{}^*N_C{}^*N_D)$

Therefore, the change in free energy is inversely proportional to the natural log of the product of the normalized slope (we want inverse proportionality such that increases in slope are equal to lowering of free energy).

*1° Epistasis ($E_{1, ABCD}$):* $E_{1,ABCD} = P_{1, ABCD}/N_{1, ABCD}$

The first-degree epistasis, based only on single mutation data, is the observed normalized slope divided by the first degree predicted normalized slope.

*2° Predicted Slope ($P_{2,ABCD}$):* $P_{2, ABCD} = N_{AB}*N_C*N_D$

The second-degree predicted slope, based only on single and double mutation data, has nine possible values based on which double or double and single mutation normalized slopes are used:

1) $N_{AB}*N_{CD}$
2) $N_{AB}*N_C*N_D$
3) $N_{AC}*N_{BD}$
4) $N_{AC}*N_B*N_D$
5) $N_{AD}*N_{BC}$
6) $N_{AD}*N_B*N_C$
7) $N_{BC}*N_A*N_D$
8) $N_{BD}*N_A*N_C$
9) $N_{CD}*N_A*N_B$

*2° Epistasis ($E_{2, ABCD}$):* $E_{2,ABCD} = P_{2, ABCD}/N_{2, ABCD}$

The second-degree epistasis, based only on single and double mutation data, has nine possible values based on which double or double and single mutation normalized slopes are used to predict the normalized slope.

For both 2° Predicted Slope and 2° Epistasis, we can work with the Maximum, Minimum, Average, and Median values independent of the observed Normalized Slope. If we know the observed Normalized Slope, we can then identify the "Best" 2° Predicted Slope based on the closest 2° Predicted Slope to the observed Normalized Slope, and "Best" 2° Epistasis score as the one closest to 1.

### Hierarchical Clustering, Motif Analysis, and Algorithmic Searching of the Landscape

We performed hierarchical clustering using a MatLab script for clustering sequences based on similarity from a prior publication.[5] We modified the script to ensure clusters had at least four unique sequences (rather than three, as more unique sequences in a cluster allows for more robust convergences) and used the most enriched 1000 unique sequences from the passage 8 (with spike-in) HT NGS. We then analyzed each cluster to identify the motif by asking whether each position (81, 84, 85, 88) converged (100%) on a single amino acid for that position for the unique sequences within that cluster and assigned the cluster a binary code based on convergence (for example, if positions 85 and 88 converge, but 81 and 84 do not, then the binary code was 0011). For the algorithmic searching, we relied on a custom web portal we developed (https://bryandickinson-create.github.io/dms-fitness-explorer/) of the fitness landscape data (**Fig. 4b**; image of portal below). For the Greedy Algorithm, we went through each possible pathway to specify the pen-Raf variant (with no prior mutations at 81, 84, 85, or 88 specified). The highest value (average of all remaining variants fitness; i.e. if 88Y has been specified and 81 needs to be specified, then each of the 20 possible AAs at position 81 have an average fitness value based on the 400 possible variants with each position 81 AA (20x20 for positions 84 and 85 which haven't been specified) is selected along each possible pathway to specify positions 81, 84, 85, and 88. Similarly, from WT, each position (81, 84, 85, 88) is specified by choosing the highest fitness point mutation available for all possible pathways to mutate at each position (e.g. for specifying in the order 88, 81, 84, 85; V88Y is the highest fitness point mutation, then from V88Y, C81H is the next highest fitness point mutation; then from C81H/V88Y, K84K (WT AA) is the highest fitness AA; finally, from C81H/K84K/V88Y, A85P is the highest fitness point mutation). Finally, rather than specifying from every possible pathway, we simply start with WT and follow every possible evolutionary pathway that includes a > +6 increase in fitness (slope). Only three single mutations from WT meet this criterion: A85K, V88Y, and V88N; thus, these three mutations constrain the initial

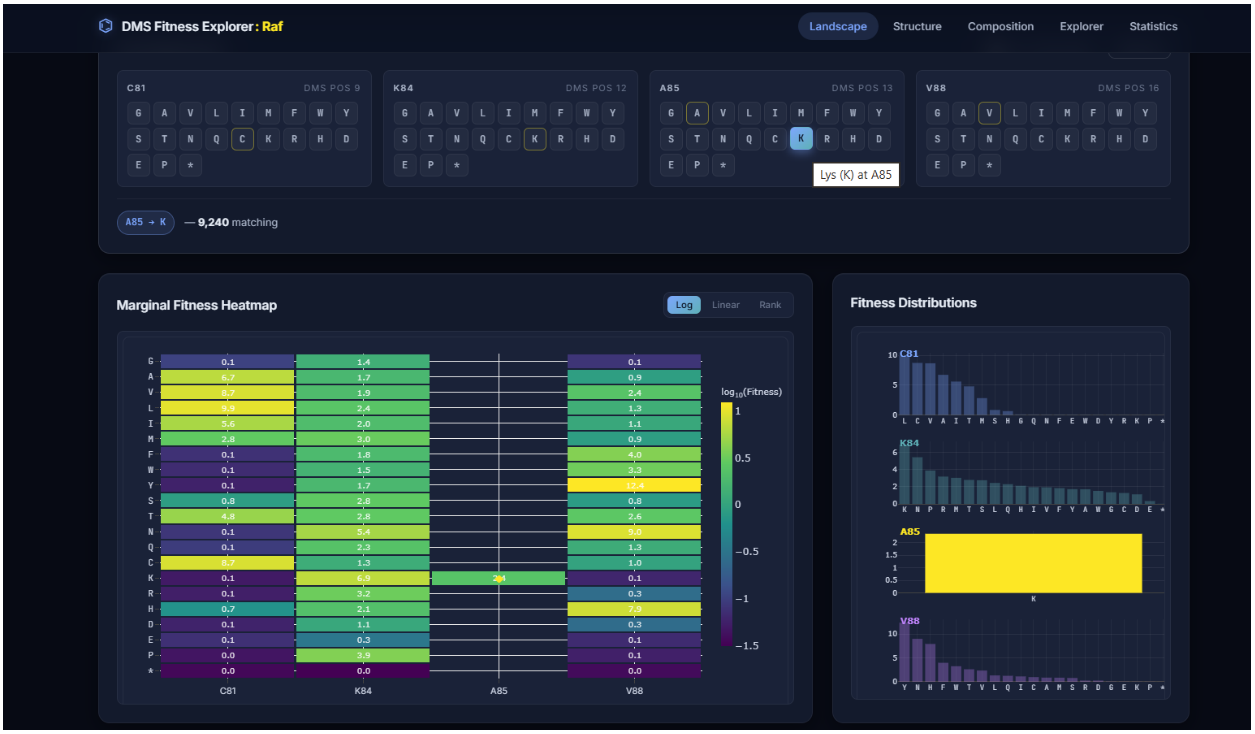

evolutionary pathways.

### NGS of *de novo* Affibody PANCS-Inhibitor Selections

We used the Amplicon EZ service provided by Genewiz (Azenta) for Illumina sequencing of each of our hits which provides 50,000+ paired end reads per sample. We used primers to install Illumina partial adaptors (red/purple) and barcodes (blue) to PCR products extending from the linker to after the stop codon of our scaffold in phage (primed with green regions) – see **Supplementary Table 7**. PCR was performed with Q5 DNAP polymerase (NEB) directly from phage (1 µL) in a 25 µL PCR reaction. The initial denaturation step was 10 min at 98 °C to release the ssDNA from the phage particle, followed by a 63 °C Ta, and a 40-second extension time over 30 cycles. PCR products are confirmed by gel (5 µL), and the remaining 20 µL of PCR product is pooled with other barcoded PCR products and column purified (Zymo DCC5). Qubit dsDNA High Sensitivity kit is used to determine an accurate concentration of the sample prior to dilution and submission for sequencing. BB Merge was used to merge each paired end reads (https://jgi.doe.gov/data-and-tools/software-tools/bbtools/bb-tools-user-guide/bbmerge-guide/) and then MATLAB was used to separate reads by barcode and to translate reads using modified scripts as described previously (MATLAB scripts provided as a supplement)[6].

### Alpha fold predictions

AlphaFold3 was used to predict binding interactions for our known binding partners (**Fig. S1** and **S3**) and for our de novo inhibitors (**Fig. S23**).[7] We implore readers to utilize these predictions only for hypothesis generation.

### Protein Purification

*General Protocol for Target Proteins for SPR*: Each target protein (Myc DBD and Mdm2) was cloned into a pET28 vector with a C-terminal 6xHis tag and transformed into BL21 *E. coli* (**Supplementary Table 6**). Cells were grown to an OD600 of 0.8 (37 °C with shaking), chilled on ice, induced with 1 mM IPTG, and then incubated with shaking at 16 °C overnight. Cells were pelleted and resuspended in a lysis buffer (25 mM Tris (pH 7.8), 10% glycerol, 200 mM NaCl). Prior to lysing by sonication, cells were treated with PMSF. The soluble fraction of the lysate was incubated with $Ni^{2+}$ resin, washed with lysis buffer containing 50 mM imidazole, then eluted in lysis buffer containing 250 mM imidazole, and finally buffer exchanged into lysis buffer and concentrated.

*General Protocol for Binder/Inhibitor Proteins for SPR*: Each binder/inhibitor variant was cloned into a pET30 vector with an N-terminal 3xFLAG and GST tag and transformed into BL21 *E. coli* (**Supplementary Table 6**). Cells were grown to an OD600 of 0.8 (37 °C with shaking), chilled on ice, induced with 1 mM IPTG, and then incubated with shaking at 16 °C overnight. Cells were pelleted and resuspended in a lysis buffer (25 mM Tris (pH 7.8), 10% glycerol, 100 mM NaCl). The soluble fraction of the lysate was incubated with GST resin, washed with lysis buffer, then eluted in lysis buffer containing 10 mM L-glutathione, and finally buffer exchanged into lysis buffer and concentrated.

*General Protocol for pen-Raf or pen-Affibody variants for protein delivery to mammalian cells*: Each binder/inhibitor variant was a pET28 vector with a N-terminal 6xHis tag and transformed into BL21 *E. coli* (**Supplementary Table 6**). Cells were grown to an OD600 of 0.8 (37 °C with shaking), chilled on ice, induced with 1 mM IPTG, and then incubated with shaking at 16 °C overnight. Cells were pelleted and resuspended in a lysis buffer (25 mM Tris (pH 7.8), 10% glycerol, 100 mM NaCl) supplemented by protease inhibitors (200 nM Aprotinin, 10 µM Bestatin, 20 µM E-64, 100 µM Leupeptin, 1 mM AEBSF, 20 µM Pepstatin A). As these "Pen" containing proteins are insoluble, we followed a previously published protocol for purifying them from inclusion bodies.[8] The cell pellet was further solubilized with 20 mL of 6 M GdHCl/50 mM Tri-HCl buffer (pH 8) by shaking at 200 rpm at room temperature for 24 hours, followed by centrifugation at 12,000 g for 60 min at 4°C. The supernatant was collected and kept, and the cell pellet was again further solubilized with 20 mL of 6 M GdHCl/50 mM Tri-HCl buffer (pH 8) by shaking at 200 rpm at room temperature for 3 hours, followed by centrifugation at 12,000 g for 60 min at 4°C, and the supernatant was collected and added to the first fraction, which was incubated with $Ni^{2+}$ resin, washed with lysis buffer containing 50 mM imidazole, then eluted in lysis buffer containing 250 mM imidazole, and finally buffer exchanged into lysis buffer and concentrated.

### Surface Plasmon Resonance

Surface Plasmon Resonance was performed on a Biacore 8000 using a NTA chip for immobilizing the His-tagged target proteins. Target concentrations were optimized to elicit a response of ~50-100 RU (180 s of 5 uL/s) and then a range of binder concentrations were tested to identify concentrations that produced robust binding (90 s of 30 uL/s). All SPR conducted at 10 °C to maintain slow dissociation of the His-tagged immobilized protein. All dose-responses were fit to a kinetic model for 1:1 binding using the Biacore evaluation software – all fits passed the quality checks in this software (**Fig. S24**).

### General Mammalian Tissue Culture

Human embryonic kidney (HEK) cell line 293T (female, ATCC) was maintained at 37◦C with 5% carbon dioxide in DMEM (L-glutamine, high glucose, sodium pyruvate, phenol red, Corning) with 10% fetal bovine serum (FBS, Gemini Benchmark) and 1x penicillin/streptomycin (GIBCO/Life Technologies). Cells were passaged at a ratio of 1:10 to 1:20 every 2-3 days when at approximately 90-100% confluency by washing with Dulbecco's phosphate-buffered saline (PBS) and treating with Trypsin-EDTA 0.25% (GIBCO) to lift.

### Mammalian Split Nano-Luciferase Assay

90 ng of the PPI reporter plasmid and 35 ng of the inhibitor expression (or GFP expression plasmid for protein delivery of inhibitor) plasmid were co-transfected into HEK293T (ATCC CRL-3216) cells using 0.375 μL of Lipofectamine 2000 in 96-well clear bottom sterile TC treated plates. Transfection was performed in triplicate or quadruplicate. After 36 hours, the Nano-luciferase activity was measured using Nano-Glo® Live Cell Assay System (Promega, N2011) on a Synergy Neo2 Microplate Reader (BioTek).

### pERK Signaling Assay

250 ng of V5-tagged KRas G12D expression plasmid was transfected into HEK293T (ATCC CRL-3216) cells using 1.5 μL of Lipofectamine 2000 in a 24-well plate. After 43 hours, media was exchanged for a low FBS media (DMEM without FBS) for 4 hours. After 47 hours (from transfection), pen-inhibitor variants were added to the cells as 2x final concentration in 20% FBS DMEM media (1:1 volume) for 1 hour, then the cells were lysed with RIPA (with phosphatase inhibitor (Santa Cruz, sc-45044 and sc-45045) and subjected to western blot analysis with the appropriate antibodies: ERK (Santa Cruz sc135900) and pERK (Santa Cruz sc136521) followed by anti-mouse HRP (Abcam, ab6728).

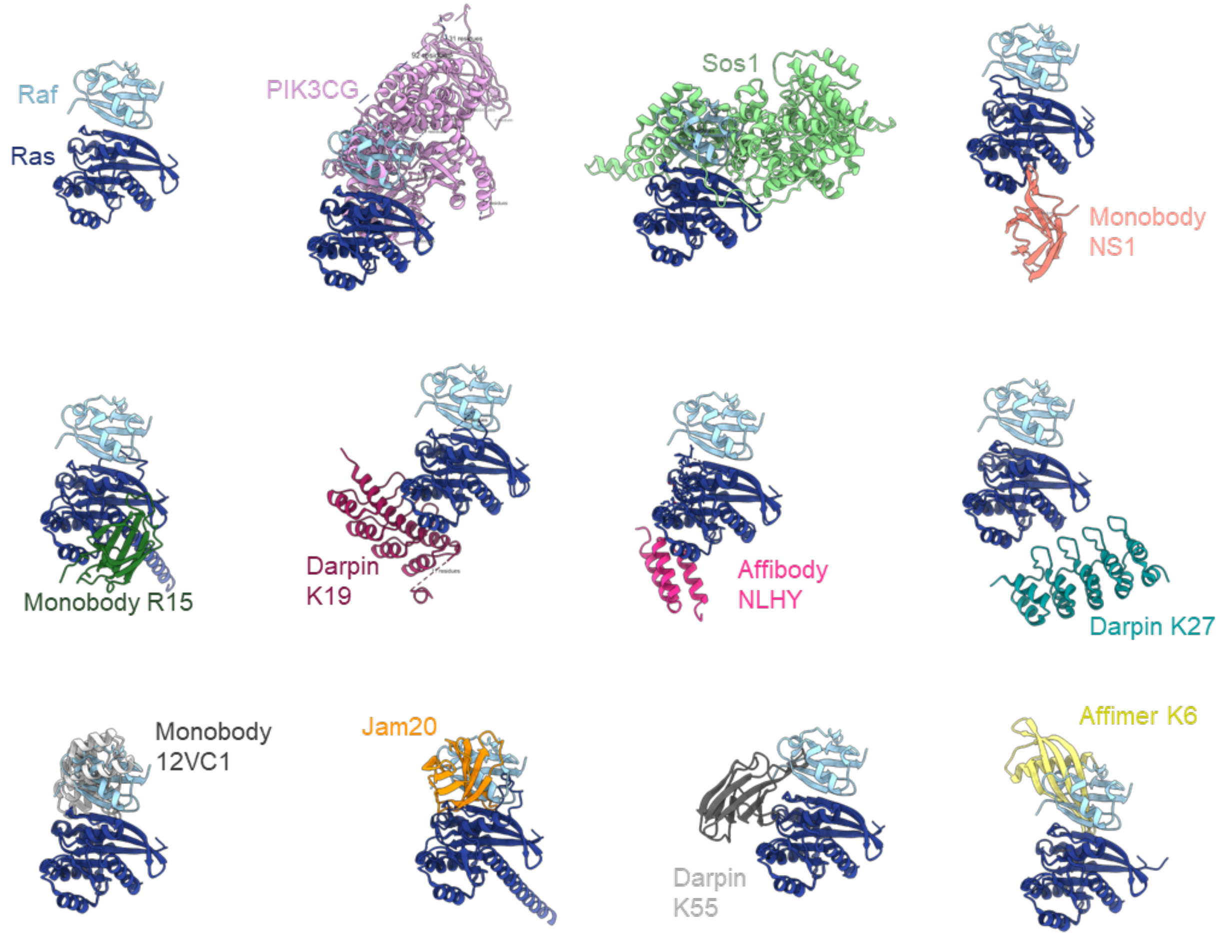

**Supplementary Figure 1. KRas-Raf interaction and *de novo* binders.** Several native PPI structures have been solved for RAS (dark blue) bound to: Raf (light blue, 6VJJ), PIK3CG (pink, 1HE8), and SOS1 (green, 1NVW). Many binders have been reported that do not inhibit the KRas-Raf interaction: Monobody NS1 (salmon, 5E95), Monobody R15 (dark green, AF3 predicted binding), DARPin K19 (maroon, 6H47), affibody NLHY (hot pink, 9Z5W), and DARPin K27 (dark teal, 5O2S); and binders that do inhibit the KRas-Raf interaction: Monobody 12VC1 (dark grey, 7L0G), JAM20 (orange, AF3 predicted binding), Affimer K6 (yellow, 6YR8), and Darpin K55 (white, 5MLA). Various Ras isoforms (HRas, KRas), mutants, and ligands were involved in these structures, but the Ras structures were aligned to KRas in the KRas-Raf structure (6VJJ).

**Supplementary Table 1: Sequences of PPI partners and previously reported binders and inhibitors.**

| ID | Library[ref] | AA Sequence |
|---|---|---|
| KRas (4b) | Uniprot: P01116 | MTEYKLVVVGAGGVGKSALTIQLIQNHFVDEYDPTIEDSYRKQVVIDGETCLLDILDTAGQEEYSAMRDQYMRTGEGFLCVFAINNTKSFEDIHHYREQIKRVKDSEDVPMVLVGNKCDLPSRTVDTKQAQDLARSYGIPFIETSAKTRQGVDDAFYTLVREIRKHKEKMSKDGKKKKKKSKTKCVIM |
| Raf (52-131; Ras binding domain) | Uniprot: P04049 | MSKTSNTIRVFLPNKQRTVVNVRNGMSLHDCLMKALKVRGLQPECCAVFRLLHEHKGKKARLDWNTDAASLIGEELQVDFL |
| Mdm2 (1-188) | Uniprot: Q00987 | MCNTNMSVPTDGAVTTSQIPASEQETLVRPKPLLLKLLKSVGAQKDTYTMKEVLFYLGQYIMTKRLYDEKQQHIVYCSNDLLGDLFGVPSFSVKEHRKIYTMIYRNLVVVNQQESSDSGTSVSENRCHLEGGSDQKDLVQELQEEKPSSSHLVSRPSTSSRRRAISETEENSDELSGERQRKRHKSDS |
| p53 (1-61; activation domain) | Uniprot: P04637 | MEEPQSDPSVEPPLSQETFSDLWKLLPENNVLSPLPSQAMDDLMLSPDDIEQWFTEDPGPD |
| Myc (368-451; DNA binding domain) | Uniprot: P01106 | MNVKRRTHNVLERQRRNELKRSFFALRDQIPELENNEKAPKVVILKKATAYILSVQAEEQKLISEEDLLRKRREQLKHKLEQL |
| Max | Uniprot: P61244 | MDKRAHHNALERKRRDHIKDSFHSLRDSVPSLQGEKASRAQILDKATEYIQYMRRKNHTHQQDIDDLKRQNALLEQQGEHP |
| Pen-Raf | [8] | RGSHHHHHHGMASLVPRGSMRQIKIWFQNRRMKWKKGSTMSGYPYDVPDYAGSMGPSKTSNTIRVFLPNKQRTVVNVRNGMSLHDCLMKALKVRGLQPECCAVFRLLHEHKGKKARLDWNTDAASLIGEELQVDFL |
| Pen-cRAFv1 | [8] | RGSHHHHHHGMASLVPRGSMRQIKIWFQNRRMKWKKGSTMSGYPYDVPDYAGSMGPSKTSNTIRVLLPNQEWTVVKVRNGMSLHDSLMKALKRHGLQPESSAVFRLLHEHKGKKARLDWNTDAASLIGEELQVDFL |
| cRAFv1 | [9] | SKTSNTIRVLLPNQEWTVVKVRNGMSLHDSLMKALKRHGLQPESSAVFRLLHEHKGKKARLDWNTDAASLIGEELQVDFL |
| Afb (Raf) | [10] | VDNKFNKEVNLAADEIWLLPNLNNQQAWAFITSLKDDPSQSANLLAEAKKLNDAQAPK |
| CI2 (Mdm2) | [11] | SSVEKKPEGVNTGAGDRHNLKTEWPELVGKSVEEAKKVILQDKPEAQIIVLPVGTMPRFMDYWEGLNRIDRVRLFVDKLDNIAEVPRVG |
| Aft-SYA (Raf) | 10^8 affitin[2] | VKVKFSYAGEEKEVDTSKIYHVSRLGKDVIFCYDDNGKVGSGAVFEKDAPKELLDMLARAEREKKL |
| Afb-TVPN (Raf) | 10^8 affibody[2] | VDNKFTKFVPNASDQFLLPNLNDYQVLAFLHSLRDPPSQSANLLAEAKKLNDAQAPK |
| Aft-CSF (Raf) | 10^10 affitin[2] | GSVKVKFCFSGEEKEVDTSKIVIVTRSGKDVVFFYDDNGKSGIGAVYEKDAPKELLDMLARAEREKKL |
| Afb-NWCN (Raf) | 10^10 affibody[2] | VDNKFNKEWCNADTEICNLPNLNDAQRCAFINSLYDDPSQSANLLAEAKKLNDAQAPK |
| Afb-YWID (Raf) | 10^10 affibody[2] | VDNKFYKEWIDADTEIANLPNLNDSQKAAFIDSLYNDPSQSANLLAEAKKLNDAQAPK |
| Aft-IFS (KRas) | 10^8 affitin[2] | GSVKVKFIFSGEEKEVDTSKIHAVFRNGKYVTFFYDDNGKHGFGSVLEKDAPKELLDMLARAEREKKL |
| Afb-NVCD (KRas) | 10^8 affibody[2] | VDNKFNKEVCDAYAQIGNLPNLNVYQIVAFIRSLDNDPSQSANLLAEAKKLNDAQAPK |

| | | |
|---|---|---|
| Aft-IYF (KRas) | 10^10 affitin[2] | GSVKVKFIYFGEEKEVDTSKISSVFRNGKFVTFFYDDNGKHGYGSVSEKDAPKELLDMLARAEREKKL |
| Afb-NLHY (KRas) | 10^10 affibody[2] | VDNKFNKELHYAIHEIGNLPNLNVYQIIAFVNSLDNDPSQSANLLAEAKKLNDAQAPK |
| Afb-TWDN | 10^8 affibody[2] | VDNKFTKEWDNASVQIVLPNLNDTQCAFVSLSDDPSQSANLLAEAKKLNDAQAPK |
| Aft-TIA | 10^8 affitin[2] | GSVKVKFTIAGEEKEVDTSKITPVARNGKSVCFLYDDNGKSGIGIVAEKDAPKELLDMLARAEREKKL |
| Afb-YWCT A46T | PACE evolved[2] | VDNKFYKEWCTASIQILSLPNLNPCQESAFLGSLVDPPSQSANLLTEAKKLNDAQAPK |
| Aft-LDA | 10^8 affitin[2] | VKVKFLDAGEEKEVDTSKIRLVDRFGKIVIFTYDDNGKVGVGDVDEKDAPKELLDMLARAEREKKL |
| Afb-SFVT | 10^8 affibody[2] | VDNKFSKEVFTAFDQIGGLPNLNLCQKDAFICSLVDDPSQSANLLAEAKKLNDAQAPK |
| Aft-SSA | 10^8 affitin[2] | VKVKFSSAGEEKEVDTSKIFVVRRLGKYVGFTYDDNGKAGLGLVCEKDAPKELLDMLARAEREKKL |
| Pen-Raf_E1 | pen- Raf (this work) | RGSHHHHHHGMASLVPRGSMRQIKIWFQNRRMKWKKGSTMSGYPYDVPDYAGSMGPSKTSNTIRVFLPNKQRTVVNVRNGMSLHD**L**M**NK**LK**N**RGLQPECCAVFRLLHEHKGKKARLDWNTDAASLIGEELQVDFL |
| Pen-Raf_E2 | pen- Raf (this work) | RGSHHHHHHGMASLVPRGSMRQIKIWFQNRRMKWKKGSTMSGYPYDVPDYAGSMGPSKTSNTIRVFLPNKQRTVVNVRNGMSLHD**V**LM**KP**LK**Y**RGLQPECCAVFRLLHEHKGKKARLDWNTDAASLIGEELQVDFL |
| Pen-Raf_E3 | pen- Raf (this work) | RGSHHHHHHGMASLVPRGSMRQIKIWFQNRRMKWKKGSTMSGYPYDVPDYAGSMGPSKTSNTIRVFLPNKQRTVVNVRNGMSLHD**V**LM**NK**LK**T**RGLQPECCAVFRLLHEHKGKKARLDWNTDAASLIGEELQVDFL |
| Pen-Raf_E4 | pen- Raf (this work) | RGSHHHHHHGMASLVPRGSMRQIKIWFQNRRMKWKKGSTMSGYPYDVPDYAGSMGPSKTSNTIRVFLPNKQRTVVNVRNGMSLHD**L**LM**NK**LK**Y**RGLQPECCAVFRLLHEHKGKKARLDWNTDAASLIGEELQVDFL |
| Pen-Raf_E5 | pen- Raf (this work) | RGSHHHHHHGMASLVPRGSMRQIKIWFQNRRMKWKKGSTMSGYPYDVPDYAGSMGPSKTSNTIRVFLPNKQRTVVNVRNGMSLHD**L**LM**KK**LK**Y**RGLQPECCAVFRLLHEHKGKKARLDWNTDAASLIGEELQVDFL |
| Pen-Raf_E6 | pen- Raf (this work) | RGSHHHHHHGMASLVPRGSMRQIKIWFQNRRMKWKKGSTMSGYPYDVPDYAGSMGPSKTSNTIRVFLPNKQRTVVNVRNGMSLHD**L**LM**KK**LK**H**RGLQPECCAVFRLLHEHKGKKARLDWNTDAASLIGEELQVDFL |
| Pen-Raf_E7 | pen- Raf (this work) | RGSHHHHHHGMASLVPRGSMRQIKIWFQNRRMKWKKGSTMSGYPYDVPDYAGSMGPSKTSNTIRVFLPNKQRTVVNVRNGMSLHD**A**LM**RK**LK**Y**RGLQPECCAVFRLLHEHKGKKARLDWNTDAASLIGEELQVDFL |
| Pen-Raf_E8 | pen- Raf (this work) | RGSHHHHHHGMASLVPRGSMRQIKIWFQNRRMKWKKGSTMSGYPYDVPDYAGSMGPSKTSNTIRVFLPNKQRTVVNVRNGMSLHD**M**LM**NK**LK**N**RGLQPECCAVFRLLHEHKGKKARLDWNTDAASLIGEELQVDFL |
| Pen-Raf_E9 | pen- Raf (this work) | RGSHHHHHHGMASLVPRGSMRQIKIWFQNRRMKWKKGSTMSGYPYDVPDYAGSMGPSKTSNTIRVFLPNKQRTVVNVRNGMSLHD**V**LM**NK**LK**Y**RGLQPECCAVFRLLHEHKGKKARLDWNTDAASLIGEELQVDFL |
| Pen-Raf_ E139 | pen- Raf (this work) | RGSHHHHHHGMASLVPRGSMRQIKIWFQNRRMKWKKGSTMSGYPYDVPDYAGSMGPSKTSNTIRVFLPNKQRTVVNVRNGMSLHD**L** |

| | | |
|---|---|---|
| | | LM**KP**LK**R**RGLQPECCAVFRLLHEHKGKKARLDWNTDAASLIGEE LQVDFL |
| Pen-Raf_ E398 | pen- Raf (this work) | RGSHHHHHHGMASLVPRGSMRQIKIWFQNRRMKWKKGSTMSG YPYDVPDYAGSMGPSKTSNTIRVFLPNKQRTVVNVRNGMSLHD**L** LM**KA**LK**R**RGLQPECCAVFRLLHEHKGKKARLDWNTDAASLIGEE LQVDFL |
| Afb-SGAD | Affibody Inhib (this work) | VDNKFSKEGADAALEIFLLPNLNLIQLRAFLCSLFDDPSQSANLLA EAKKLNDAQAPK |
| Afb-TWAA | Affibody Inhib (this work) | VDNKFTKEWAAATAEILALPNLNQVQEPAFLDSLCDDPSQSANLL AEAKKLNDAQAPK |
| Telobody (Myc) | [12] | LQLREPSFPDVQHVVLIHKVILGSPAHRAGLWPGDVILAIGEQMV QNAEDVYEAVRTQSQLAVQIRRGRETLTLYVTPEVTE |
| Monobody (KRAS) | [13] | GSVSSVPTKLEVVAATPTSLLISWDAPAVTVDYYVITYGETGGNS PVQKFEVPGSKSTATISGLKPGVDYTITVYAWGWHGQVYYYMGS PISINYRT |
| Affibody (PDL1) | [14] | VDNKFNKEPRAARLEITVLPNLNREQGGAFIVSLWDDPSQSANLL AEAKKLNDAQAPK |

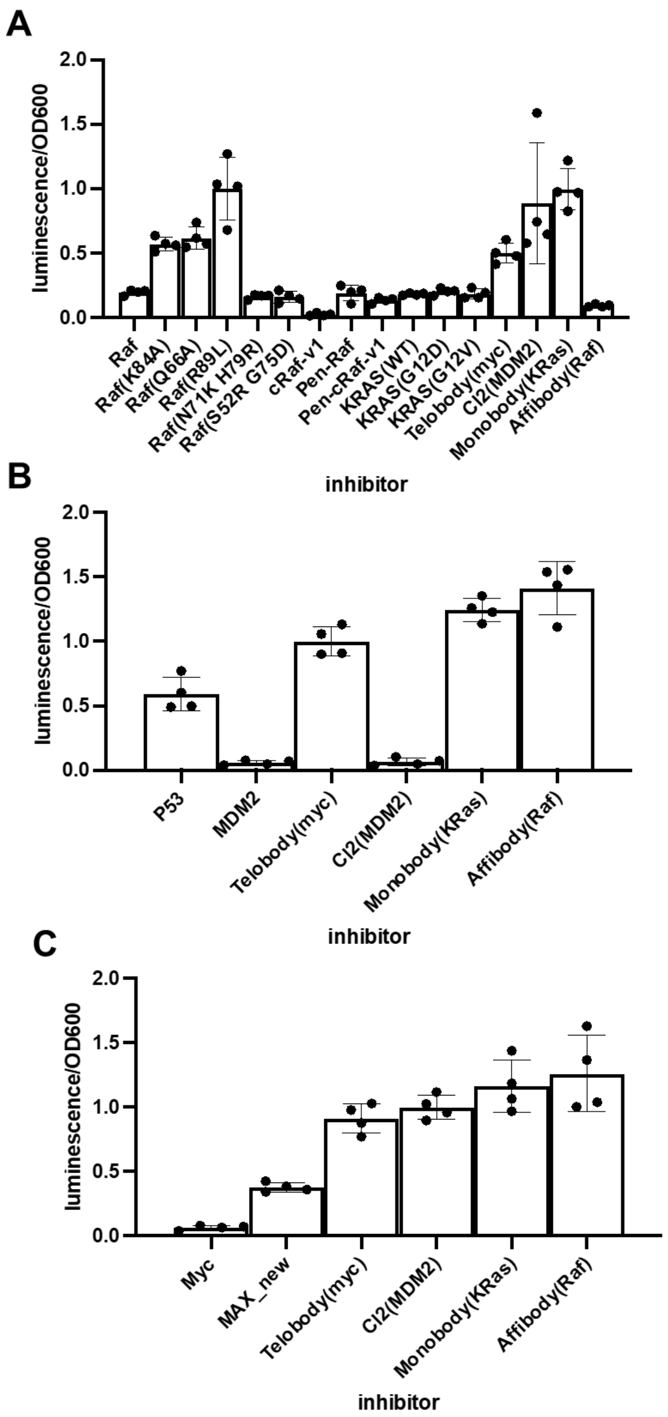

**Supplementary Figure 2.** Additional controls in *E. coli* PPI inhibition luxAB assay with A) KRas-Raf, B) Mdm2-p53, and C) Myc-Max (n=4, error bars indicate SD). See Supplementary Table S1 for details of proteins tested as inhibitors (listed below each plot). Telobody (myc)[12] and Monobody (KRas)[13] showed no inhibition and were not used in further assays.

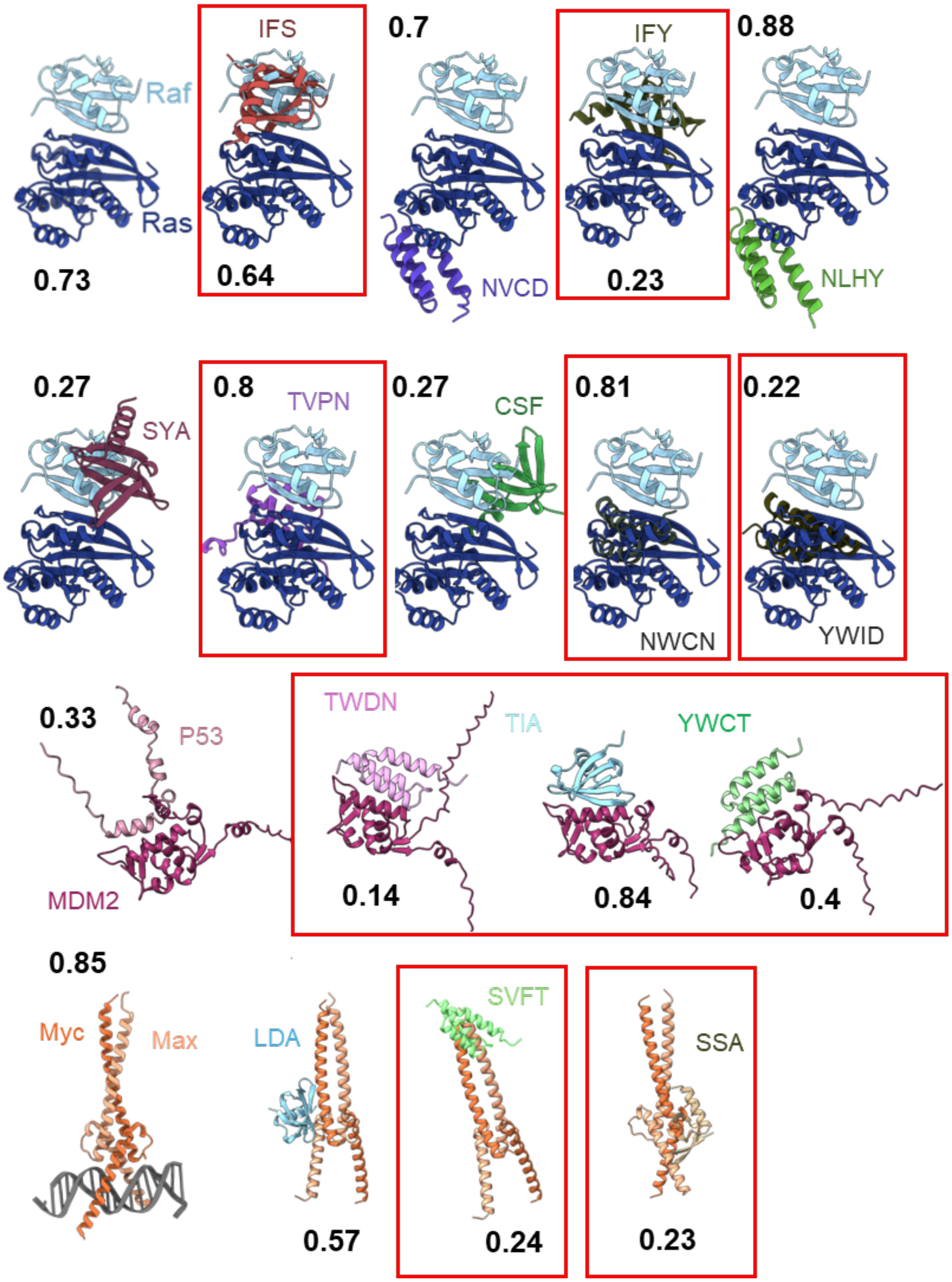

**Supplementary Figure 3.** Alphafold3 predicted interactions with our previously reported binder-target complexes (See **Supplementary Table 1** for sequences) aligned with native target PPI complexes. On the left, each PPI is shown (second row has five Raf binders rather than the native PPI which is shown in row 1), followed by each predicted binder-target structure (top: KRas binders; upper middle: Raf binders; lower middle: Mdm2 binders; and bottom: Myc and then Max binders). In black, iPTM values are listed below each structure. Each binder that is predicted to be an inhibitor is boxed in red.

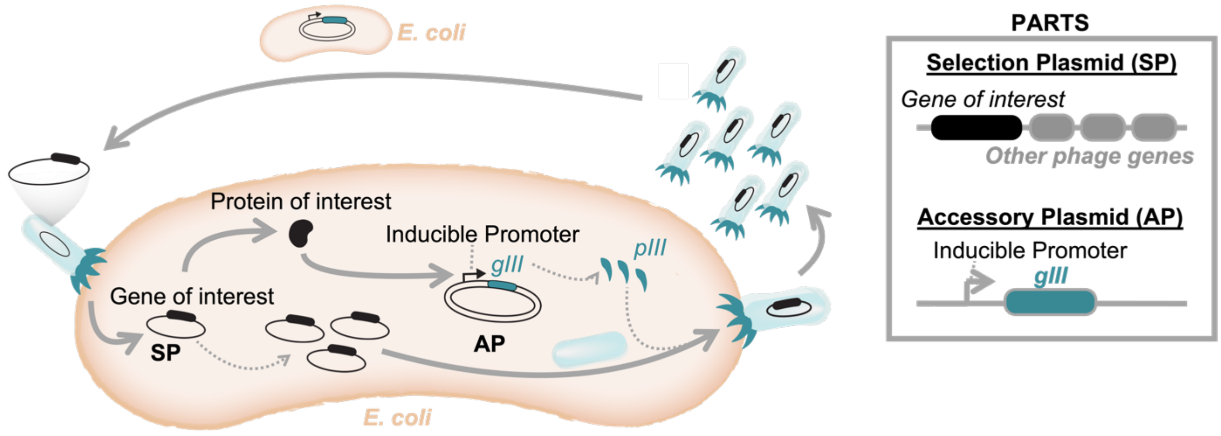

**Supplementary Figure 4. *In vivo* phage-assisted directed evolution platforms, in general.** *In vivo* phage-assisted directed evolution platforms function by encoding a gene of interest (black) into the phage genome (SP) in place of the essential *gIII*. *gIII* (teal) is instead encoded in an accessory plasmid (AP) in the *E. coli* and placed under control of an inducible promoter. The researcher must engineer a biosensor to link desired protein activity with the induction of p**III**. In this case, phage with proteins with the desired activity will be able to propagate on these engineered *E. coli* cells.

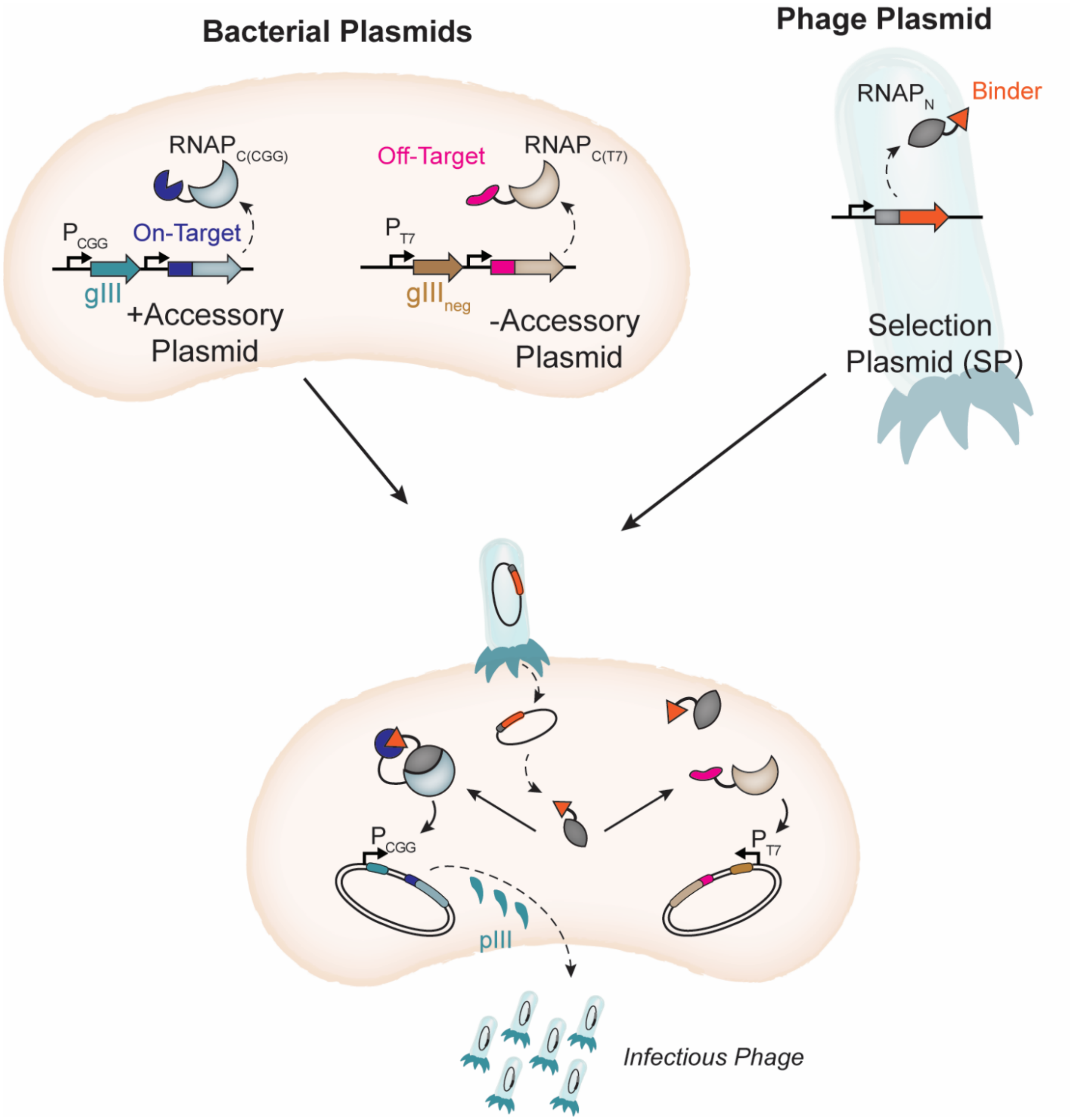

**Supplementary Figure 5.** PANCS-Binder selection platform. Our previously reported PANCS-Binder selection platform utilizes a two plasmid (+AP, -AP) bacterial biosensor (upper left) in which an on-target protein and an off-target protein are each fused to orthogonal C-terminal RNA polymerase (RNAPc) which control gIII and $gIII_{neg}$ expression, respectively. In the phage plasmid (upper right), SP, a binder (or potential binder library) is fused to the N-terminal RNAP (RNAPn). Phage infection (lower) results in expression of the RNAPn-Binder fusion and on-target binding dependent phage replication.

**Supplementary Note 1: Failed PANCS-Inhibitor Biosensor Designs.**

**Design 1**: We first attempted to create the PANCS-inhibitor platform by using the same SP and APs as the PACE platform we designed to evolve PPIs, but also adding the genetically encoded inhibitor into the SP (**Note Figure A**). However, we found that even out of a theoretically monoclonal phage population of $10^9$ phage, cheaters existed, purportedly through chance natural mutagenesis, that rendered the PPI Partner 1 in the phage non-functional, thus bypassing the negative selection. Even when we had two copies of PPI Partner 1 in the phage, there still existed cheater phage that removed the PPI Partner 1 genes, thus bypassing the selection.

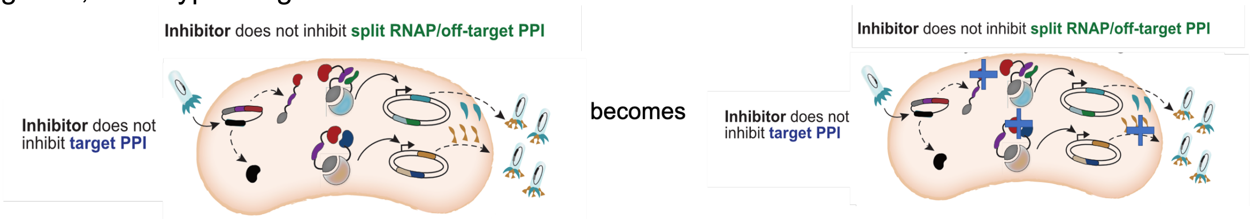

**Note Figure A.** Schematic showing supplementary note, design 1.

**Example 1**: We added $10^9$ of the following phage to S1030 *E. coli* cells with the following APs and performed 4 passages with 1:20000 dilution rate of phage (**Note Figure B**). The phage population over all passages was $10^{11}$ and phage variants at the end contained Raf variants in the phage that had the following inactivating mutations: Q66taa(stop) and R89C.

**Example 2**: We added $10^9$ of the following phage to S1030 *E. coli* cells with the following APs and performed 4 passages with 1:20000 dilution rate of phage (**Note Figure B**). The phage population over all passages was $10^{11}$ and phage variants at the end all contained the first Raf deleted and some also contained the Q66taa(stop) in the second Raf.

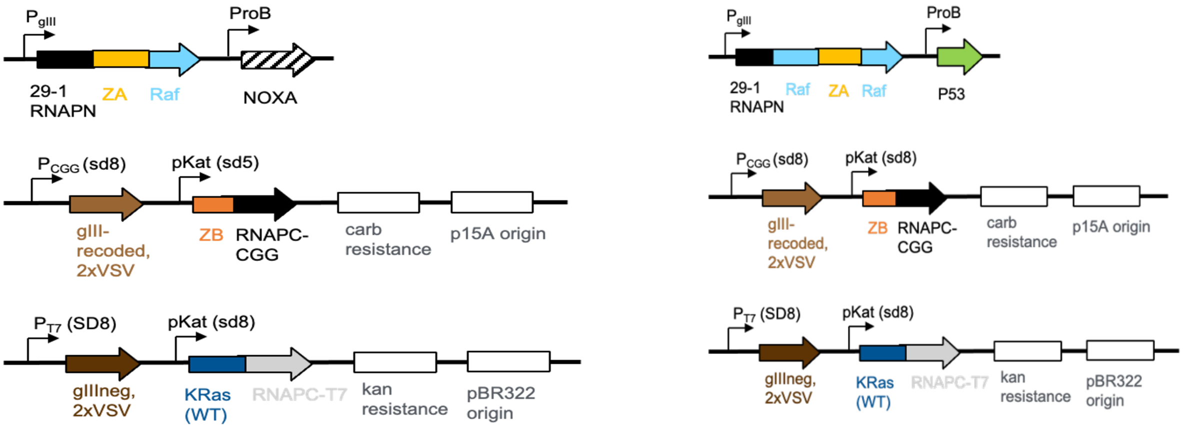

**Note Figure B. (Left)** Plasmid maps corresponding to supplementary note, design 1, example 1. **(Right)** Plasmid maps corresponding to supplementary note, design 1, example 2.

**Design 2**: We then attempted to modify this platform by removing the $RNAP_N$-ZA-PPI Partner 1 from the phage and encoding into an AP (**Note Figure C**). However, this enabled the *E. coli* to produce gIII prior to phage infection, which is known to prevent phage infection and thus prohibit our selection to take place. We experimentally observed this as well.

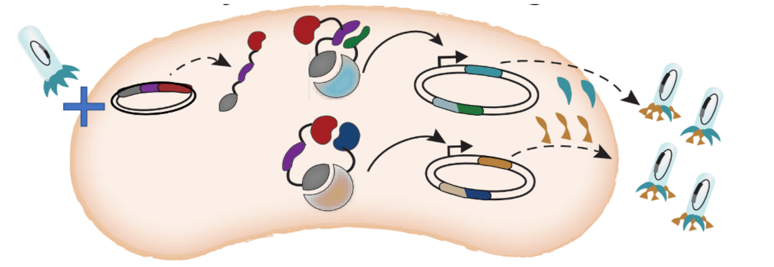

**Note Figure C.** Schematic showing supplementary note, design 2.

**Example 3**: Overnight phage growth assays starting with 1000 phage resulted in successful RNAPn-ZA phage replication when only the RNAPc was present in the *E. coli* strain (left), but no phage replication when the RNAPn-ZA-Raf (right) was added to the E. coli strain (**Note Figure D**); even cheater full RNAP containing phage were unable to replicate on this later strain indicating an inability to infect (pending expression level).

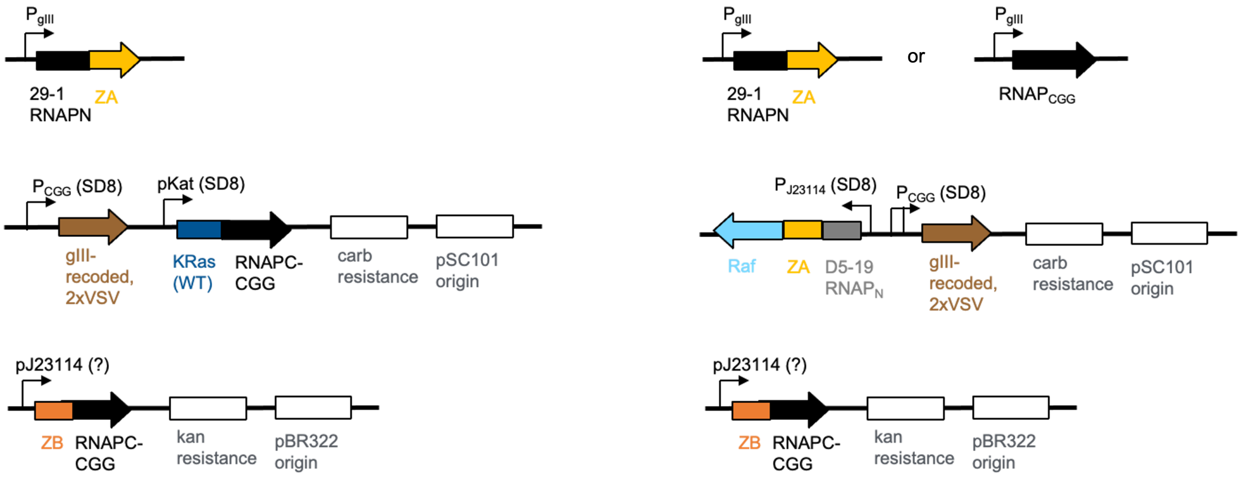

| RBS for pJ23114 | Titer (PFU/mL) |
|---|---|
| sd5 | 10^5 |
| sd8 | 10^11 |

| RBS for pJ23114 | Titer with full-length CGG phage (PFU/mL) | Titer with $RNAP_N$-ZA phage (PFU/mL) |
|---|---|---|
| sd5 | 10^10-11 | 10^6 |
| sd8 | <10^3 | <10^3 |

**Note Figure D**. Plasmid maps and phage growth assays corresponding to Supplementary Note, Design 2**,** Example 3. Number reported is PFU/mL of one replicate.

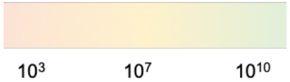

| | | phage | |
|---|---|---|---|
| | Name | 53-78, +control phage 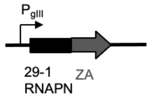 | 73-119, empty phage 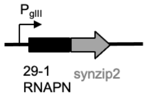 |
| +AP | 71-123 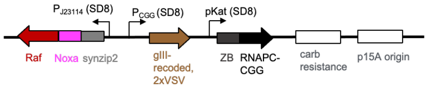 | 1.00E+11 | 2.00E+03 |
| | 71-125 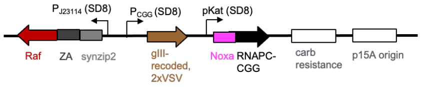 | 1.00E+03 | 3.00E+03 |

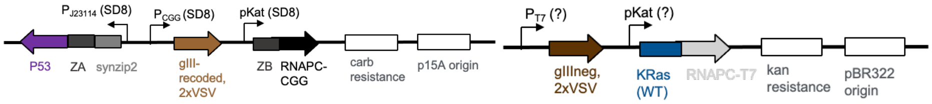

| +AP 71-111 | | | | | |
|---|---|---|---|---|---|
| | | phage (name, inhibitor) | | | |
| | | 53-78, +control phage | 73-119, empty phage | 71-114, MDM2 | 73-121, Pen-Raf |
| -AP (name, RBS of pT7, RBS of pKat) | 59-43, sd8, sd8 | 1.00E+11 | 1.00E+10 | 1.00E+10 | 1.00E+10 |
| | 59-42, SD8, sd8 | 1.00E+11 | 1.00E+10 | 1.00E+10 | 1.00E+10 |
| | 66-82 SD8, SD8 | 1.00E+11 | 1.00E+07 | 1.00E+09 | 1.00E+10 |

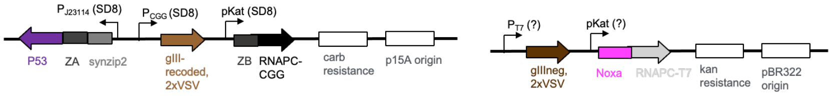

| +AP 73-120 | | | | | |
|---|---|---|---|---|---|
| | | phage (name, inhibitor) | | | |
| | | 53-78, +control phage | 73-119, empty phage | 71-114, MDM2 | 73-121, Pen-Raf |
| -AP (name, RBS of pT7, RBS of pKat) | 23-49, sd8, sd8 | 1.00E+11 | 1.00E+08 | 5.00E+09 | 1.00E+11 |
| | Jin 487, SD8, sd8 | 1.00E+11 | 1.00E+08 | 1.00E+08 | 1.00E+11 |
| | 28-67 SD8, SD8 | 1.00E+11 | 1.00E+07 | 1.00E+08 | 1.00E+11 |

**Supplementary Figure 6a. Optimization of AP1 (+AP) expression level.** In each case, an overnight replication (**Fig. 2C**) of 10,000 phage was performed with varying expression level APs. **(Top)** Validation of a set of control phage growth assays to test PANCS-inhibitor. Plasmid maps are shown above corresponding phage growth assay data. Number reported is PFU/mL of one replicate. **(Middle)** Non-binding PPI partners (p53 on adaptor and KRas on AP2 (-AP)). (Bottom) A second example of Non-binding partners (Noxa on AP2 (-AP)). RBS expression on AP2 (-AP) are listed to the left of the heatmap as plasmid ID, $P_{T7}$ (*gIII$_{neg}$* expression), $P_{kat}$ (PPI partner-$RNAP_{c,T7}$ expression). These results suggest the +AP 73-120 expression level produces high replication in the absence of productive PPI between the target proteins.

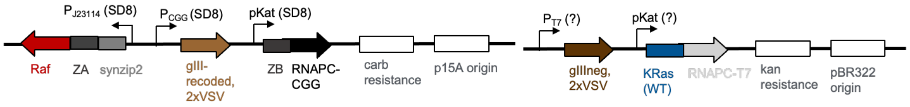

| +AP 73-120 | | | | | | | | | |
|---|---|---|---|---|---|---|---|---|---|
| | | phage (name, inhibitor) | | | | | | | |
| | | 53-78, +control phage | 73-119, empty phage | 73-121, Pen-Raf | 73-122, Pen-cRaf-v1 | 73-123, Pen-Raf (Q66A) | 73-118, Raf (R89L) | 71-157, Monobody (KRas) | 71-158, Affibody (Raf) |
| -AP (name, RBS of PT7, RBS of pKat) | 59-55, sd5, sd2 | 1.00E+11 | 1.00E+10 | 1.00E+10 | 1.00E+10 | 1.00E+10 | 1.00E+10 | 1.00E+11 | 1.00E+11 |
| | 66-95, sd5, sd5 | 1.00E+11 | 1.00E+08 | 1.00E+10 | 1.00E+10 | 1.00E+10 | 1.00E+08 | 1.00E+11 | 1.00E+11 |
| | 66-94, sd8, sd5 | 1.00E+11 | 5.00E+07 | 1.00E+10 | 5.00E+08 | 1.00E+10 | 1.00E+08 | 1.00E+11 | 1.00E+11 |
| | 59-43, sd8, sd8 | 1.00E+11 | 1.00E+05 | 1.00E+10 | 1.00E+09 | 1.00E+07 | 1.00E+05 | 5.00E+06 | 1.00E+09 |
| | 59-42, SD8, sd8 | 1.00E+11 | 5.00E+04 | 1.00E+10 | 1.00E+09 | 5.00E+05 | 5.00E+04 | 1.00E+06 | 1.00E+10 |
| | 66-82, SD8, SD8 | 1.00E+11 | 3.00E+03 | 2.50E+06 | 7.50E+06 | 6.00E+03 | <10^3 | 4.00E+03 | 9.50E+04 |

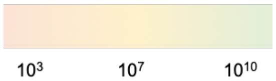

**Supplementary Figure 6b (continued). KRas-Raf AP1/2 optimization.** With optimized AP1 (+AP) expression level (SD8, SD8), we tested varied expression level AP2 (-AP) RBS levels (listed as plasmid ID, $P_{T7}$ (*gIII<sub></sub>*$_{neg}$ expression), $P_{kat}$ (PPI partner-$RNAP_{c,T7}$ expression) in overnight replication assays (**Fig. 2C**) with varied strengths of inhibitors encoded in phage. Number reported is PFU/mL of one replicate.

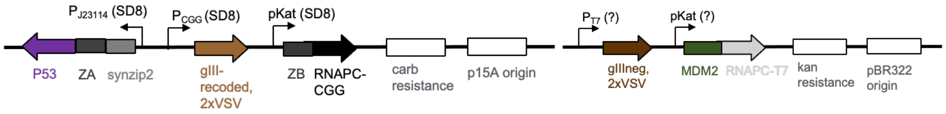

| +AP 71-111 | | | | | | |
|---|---|---|---|---|---|---|
| | | phage (name, inhibitor) | | | | |
| | | 53-78, +control phage | 73-119, empty phage | 71-114, MDM2 | 71-115, P53 | 71-156, CI2 (MDM2) |
| -AP (name, RBS of PT7, RBS of pKat) | 71-134, sd5, sd2 | 1.00E+11 | 1.00E+10 | 1.00E+11 | 1.00E+10 | 1.00E+10 |
| | 71-135, sd8, sd2 | 1.00E+11 | 1.00E+09 | 5.00E+09 | 1.00E+11 | 1.00E+11 |
| | 71-136, SD8, sd2 | 1.00E+11 | 5.00E+09 | 1.00E+11 | 1.00E+11 | 1.00E+11 |
| | 71-140, sd5, sd5 | 1.00E+11 | 1.00E+05 | 5.00E+06 | 1.00E+07 | 1.00E+10 |
| | 71-141, sd8, sd5 | 1.00E+11 | 1.00E+04 | 1.00E+05 | 1.00E+07 | 1.00E+10 |
| | 66-169, sd8, sd8 | 1.00E+11 | <10^3 | 7.00E+03 | <10^3 | <10^3 |
| | 66-168, SD8, sd8 | 1.00E+11 | <10^3 | 3.00E+03 | <10^3 | 5.00E+04 |
| | 66-167, SD8, SD8 | 5.00E+10 | <10^3 | <10^3 | 1.00E+03 | 1.00E+05 |

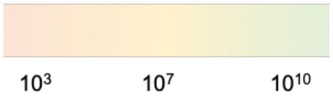

**Supplementary Figure 6c (continued). Mdm2/p53 AP1/2 optimization.** With optimized AP1 (+AP) expression level (SD8, SD8), we tested varied expression level AP2 (-AP) RBS levels (listed as plasmid ID, $P_{T7}$ (*gIII*$_{neg}$ expression), $P_{kat}$ (PPI partner-RNAP$_{c,T7}$ expression) in overnight replication assays (**Fig. 2C**) with varied strengths of inhibitors encoded in phage. Number reported is PFU/mL of one replicate.

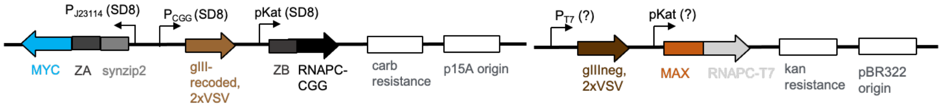

| +AP 71-143 | | | | | | | |
|---|---|---|---|---|---|---|---|
| | | phage (name, inhibitor) | | | | | |
| | | 53-78, +control phage | 73-119, empty phage | 71-149, MYC | 71-150, MYC_DBD | 71-151, MAX | 71-155, Telobody (MYC) |
| -AP (name, RBS of PT7, RBS of pKat) | 71-192, sd8, sd8 | 1.00E+11 | 8.00E+04 | 3.75E+06 | 1.00E+10 | 1.00E+11 | 1.00E+10 |
| | 71-189, SD8, sd8 | 1.00E+11 | 5.00E+04 | 1.75E+06 | 1.00E+10 | 1.00E+11 | 1.00E+08 |
| | 71-186, SD8, SD8 | 1.00E+11 | 3.00E+03 | 4.00E+04 | 1.00E+10 | 1.00E+10 | 1.80E+04 |

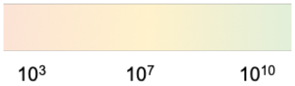

**Supplementary Figure 6d (continued). Myc/Max AP1/2 optimization.** With optimized AP1 (+AP) expression level (SD8, SD8), we tested varied expression level AP2 (-AP) RBS levels (listed as plasmid ID, $P_{T7}$ (*gIII*$_{neg}$ expression), $P_{kat}$ (PPI partner-RNAP$_{c,T7}$ expression) in overnight replication assays (**Fig. 2C**) with varied strengths of inhibitors encoded in phage. Number reported is PFU/mL of one replicate.

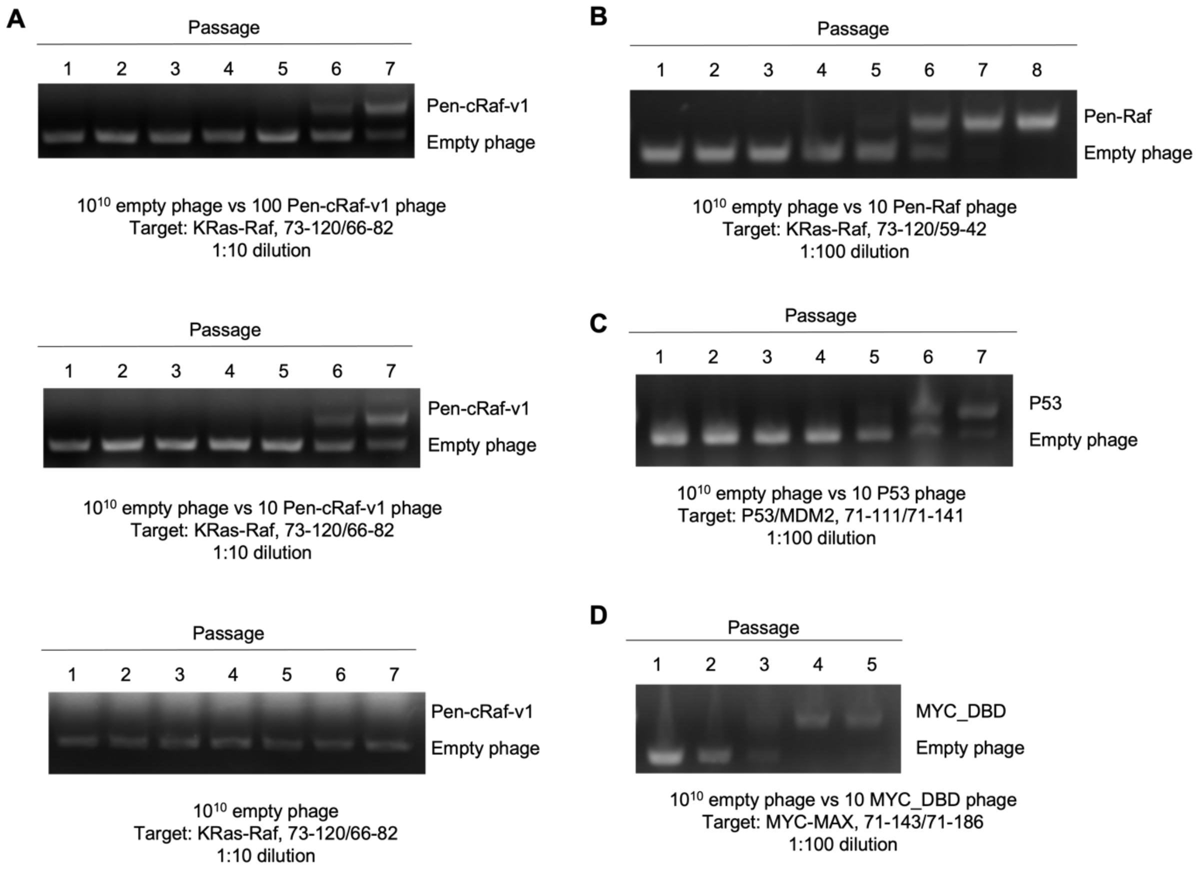

**Supplementary Figure 7. Additional mock PANCS-inhibitor experiments.** Gels with PCR products of passages from the mock PANCS of (**A**) Pen-cRAF-v1-encoding phage on one set of Raf-KRas APs starting with 100 (Top), 10 (Middle), or no Pen-cRAF-v1 phage (Bottom); (**B**) Pen-Raf-encoding phage on another set of Raf-KRas APs, (**C**) p53 phage on p53-Mdm2 APs, and (**D**) Myc phage on Myc-Max APs.

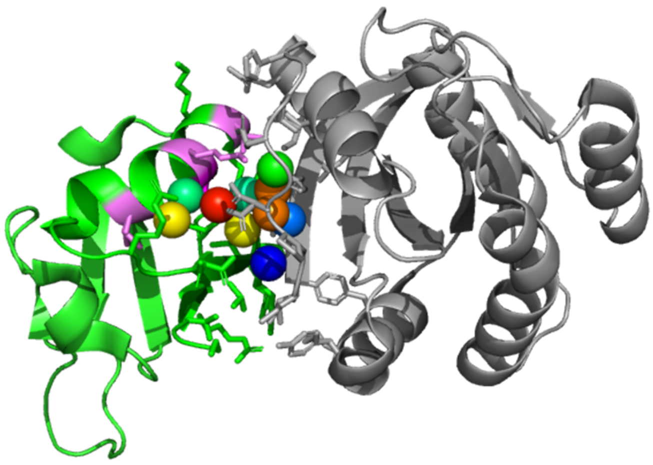

| PDB | RAS | RAF | # $H_2O$ |
|---|---|---|---|
| 6VJJ | KRAS4b GMppNp | RBD | 10 |
| 8T74 | KRAS4a GMppNp | RBD | 8 |
| 8EPW | KRAS G13D GMppNp | RBD | 5 |
| 7JHP | HRAS GMppNp | RBD+CRD | 4 |
| 6XI7 | KRAS GMppNp | RBD+CRD | 3 |
| 6XHB | KRAS GMppNp | RBD+CRD | 10 |
| 6XGV | KRAS G13D GMppNp | RBD+CRD | 7 |
| 4G0N | HRAS GMppNp | RBD | 7 |
| 1C1Y | RAP GMppNp | RBD | 12 |
| **3KUD** | **HRAS GDP** | **RBD A85K** | **0** |
| 8JNA | No RAS | RBD | 3 |

**Supplementary Figure 8. Pocket of waters at the KRas-Raf interface centered at Raf A85.** The crystal structure (left) of the Raf (green)- KRas (grey) interaction (6VJJ) shows 10 waters (spheres, colored by b-factor, red high, blue low) in between Raf and KRas centered around A85 (dark purple, the other randomized residues in our pen-Raf library are also shaded light purple). Investigation (right) of similar Ras-Raf structures shows that similar quantity of water are trapped reproducibly. Notably, an A85K mutation of Raf RBD showed zero waters in this pocket (purple line) and even in the absence of Ras, Raf RBD shows trapped water in this pocket.

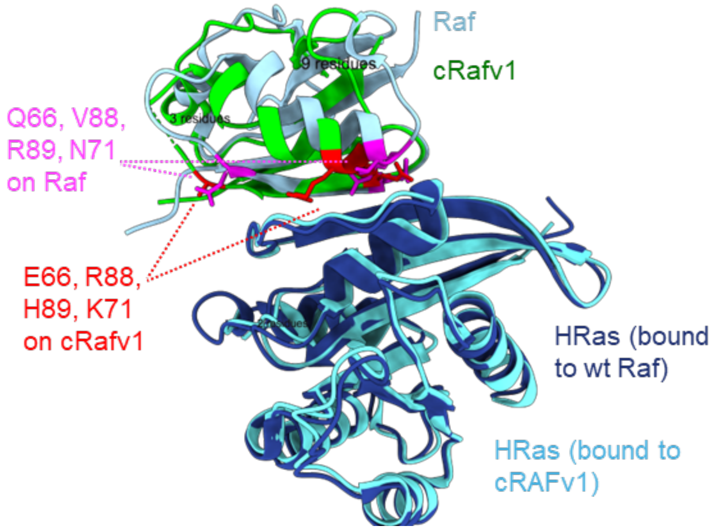

**Supplementary Figure 9. Altered KRas-Raf interaction network in the high affinity cRAFv1 variant of Raf.** HRas structure was aligned in the HRas-Raf (4G0N) and HRas-cRAFv1 (6NTC) structures. The mutations for cRAFv1 (red) alter which residues make contact with KRas.[9]

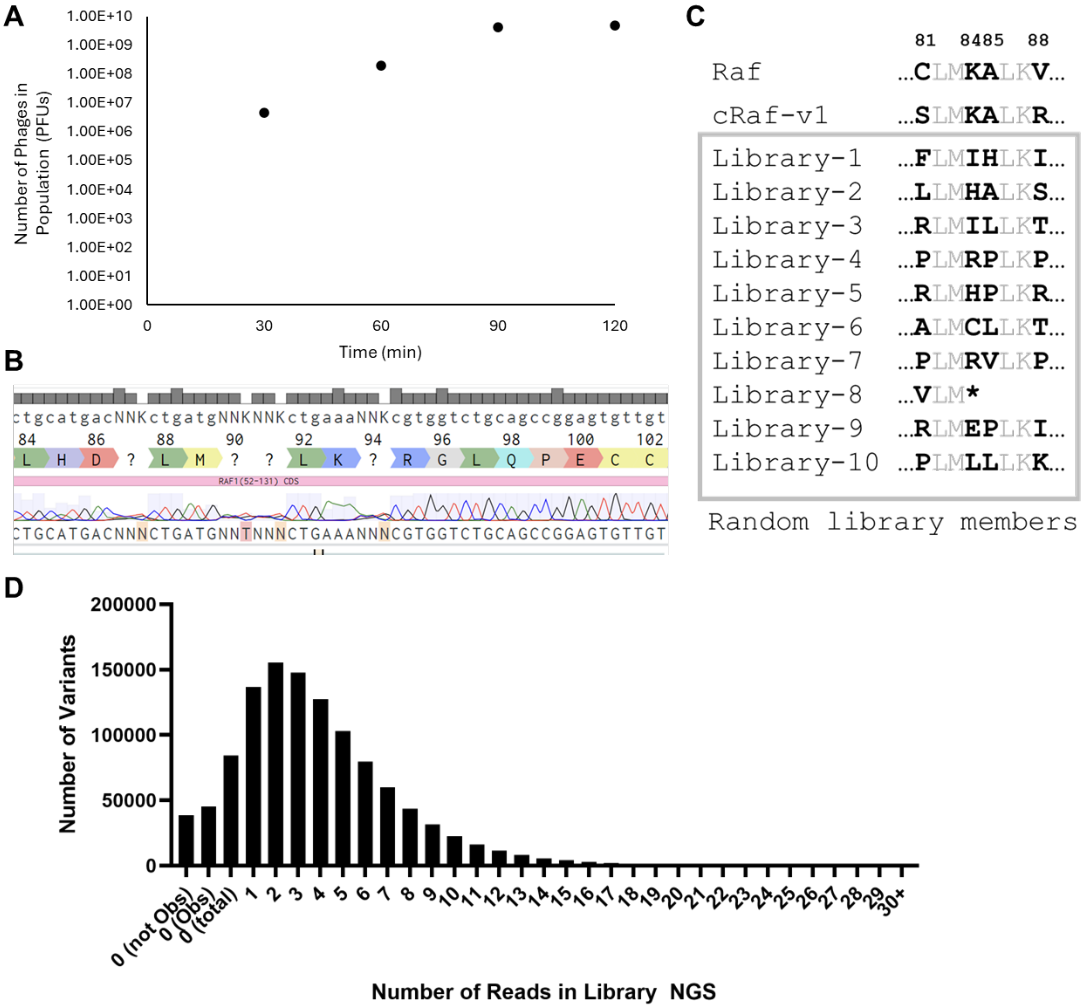

**Supplementary Figure 10. Pen-Raf Library Construction. A)** The pen-Raf 4 NNK (positions 81, 84, 85, and 88) was transformed by electroporation into *E. coli* and the phage titer of the population was monitored over time. At 30 minutes, the population was >4x the theoretical diversity, and by 60 minutes, the population was >190x the theoretical diversity. We estimate that the 30-minute timepoint is an undercount of the number of transformed cells while 60 minutes is likely a slight overcount (as many transformed cells have made multiple phage copies). Therefore, we think that this library size is likely sufficient for near universal coverage of the theoretical diversity. Each data point was collected as a single activity dependent phage titer assay. **B)** Sanger Sequencing of the bulk library showed correct diversification of the randomized positions, and **C)** isolated random library members have the correct diversification. **D)** In our NGS of the library, we did not observe a wide-spread in the distribution of variant frequency, 91% of variants had between 1 and 10 reads and <1% had more than 11 reads. 8% of variants had 0 reads in the library NGS ("0 (total)" in **D**), but most of those variants were observed in the NGS of one or more of the passages ("0 (Obs)"); thus, only 3.6% of variants had 0 reads in the library and were not observed in any of the sequencing ("0 (not Obs)").

**Supplementary Table 2. NGS Summary of pen-Raf selection.** Full length DNA sequencing reads, unique DNA library variants amongst those reads, and unique (full pen- Raf, no stop codon) protein library variants coded by those unique DNA variants. ND indicates not determined.

| | With Spike-in | | | Without Spike-in | | |
|---|---|---|---|---|---|---|
| Passage | Reads | Variants (DNA) | Variants (AA) | Reads | Variants (DNA) | Variants (AA) |
| 0 | N/A | | | 10,217,756 | 967,563 | 158,764 |
| 2 | 29,901,591 | ND | | 11,177,976 | 838,271 | 155,202 |
| 4 | 5,887,912 | 130,798 | 63,942 | 5,479,338 | 177,604 | 80,750 |
| 6 | 3,035,160 | 32,007 | 16,194 | 6,800,003 | 80,274 | 41,907 |
| 8 | 3,256,208 | 17,028 | 7,385 | 1,270,602 | 12,984 | 5,542 |

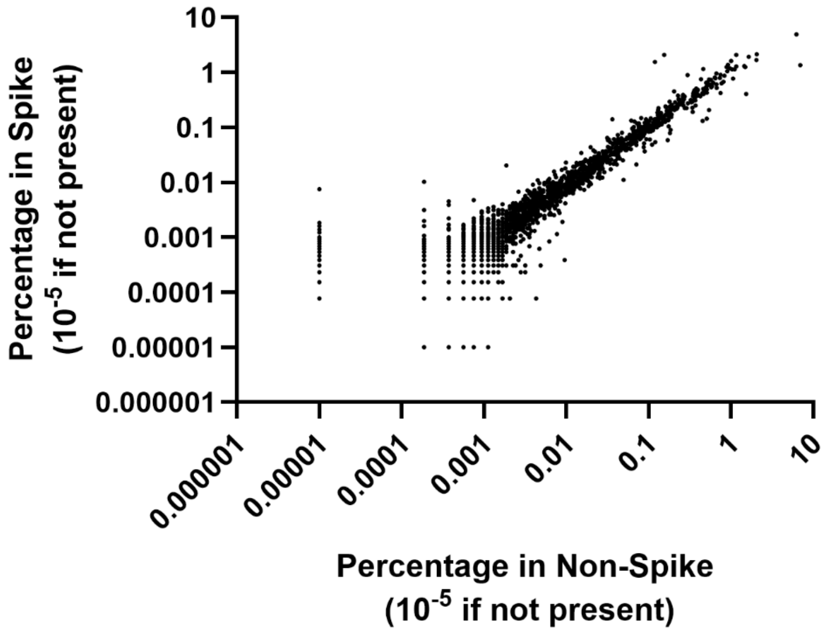

**Supplementary Figure 11. Comparison of selection with and without spike-in empty phage.** We compared the relative percent of each variant in the overall population of our selection of 4 NNK pen-Raf variants with and without a 1:1 empty phage spike-in (which was used to monitor de-enrichment by PCR) at the end of passage 8. Each variant that was observed in one of these samples, but was in the other, was changed from 0% to 0.00001% (1 read was ~0.0001% in each sample) so that the data could be plotted in log scale. We observed a very strong correlation between the two conditions suggesting that the spike-in did not substantially perturb the selection results (Pearson Correlation, r = 0.82, p = <0.001).

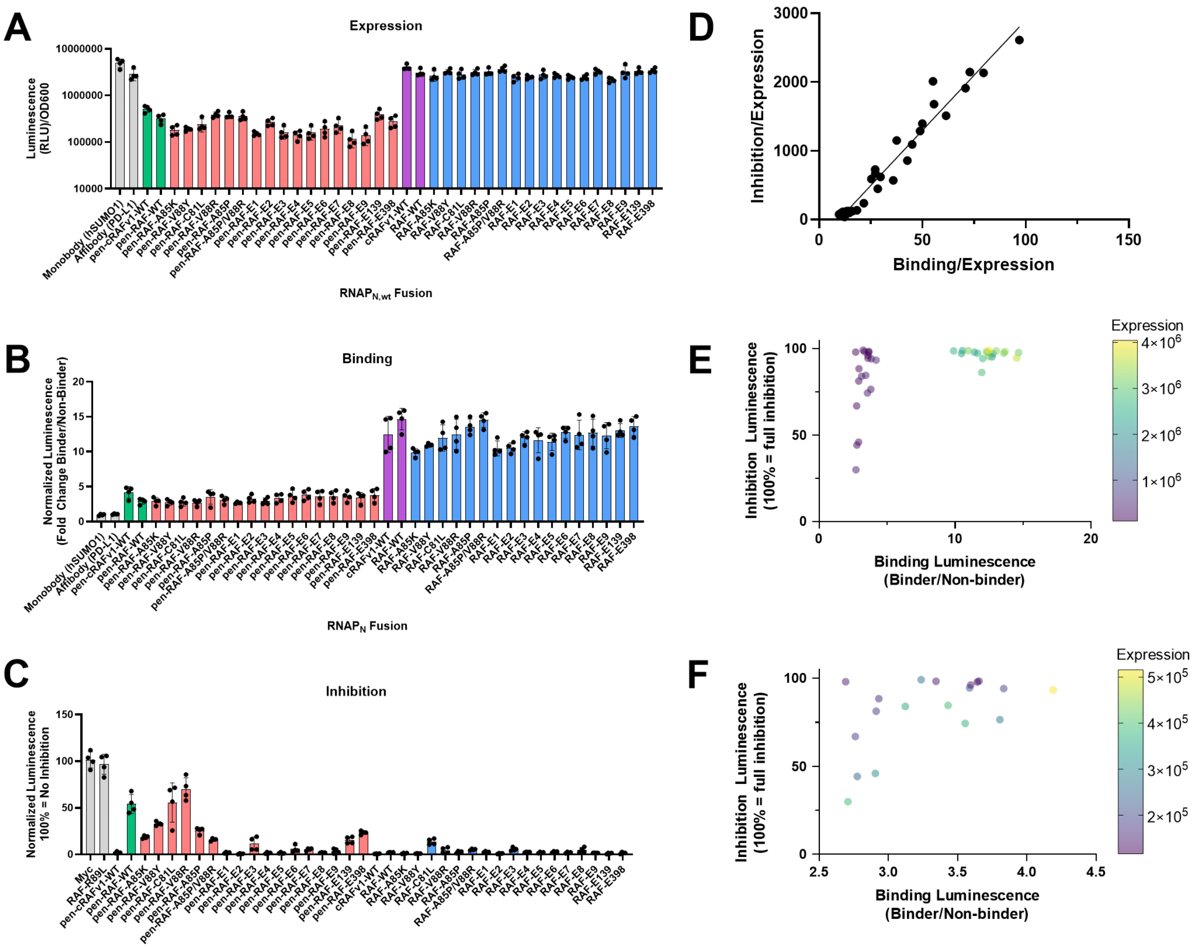

**Supplementary Figure 12. Expression, Binding, and Inhibition of pen-Raf and Raf variants. A)** The expression level of various pen-Raf and Raf variants were measured by fusing each variant to the WT split RNAP N-terminal fragment (which recombines with RNAP C-terminal independent of a binding interaction between fusion partners) with ZB-RNAP$_C$. **B)** The binding (**Fig. 1C**) of various pen-Raf and Raf variants to KRas (fused to RNAP$_N$ and RNAP$_C$ respectively). **C)** The inhibition (**Fig. 1E**) of various pen-Raf and Raf variants of the Raf - KRas PPI (RNAP$_N$- Raf and KRas-RNAP$_C$). For **A-C**, non-binding/interacting variants are shown in grey (Monobody (hSUMO1)[15] and Affibody (PD-L1)[14]), pen- Raf WT and pen-cRAFv1 in green and variants of pen- Raf in red, and the corresponding non-pen version in purple (WT) and blue (variants). For each measurement, n = 4 and error bars indicate SD. **D)** The binding value (binder/non binder) and inhibition value (normalized so 100% = full inhibition) are highly correlated when each value is divided by expression level (Pearson's Correlation, r = 0.98, p = <0.0001). **E)** Same data as in **D**, but rather than dividing by expression, expression is shown as a third variable (color of symbol). Pen variants (all purple, low expression) cluster at lower binding, but vary in the ability to inhibit. **F**) Same as **E**, but only shows Pen variants. The variant in the upper left corner (high inhibition, but low binding) is pen-RAF-E1 (most enriched library variant in passage 8).

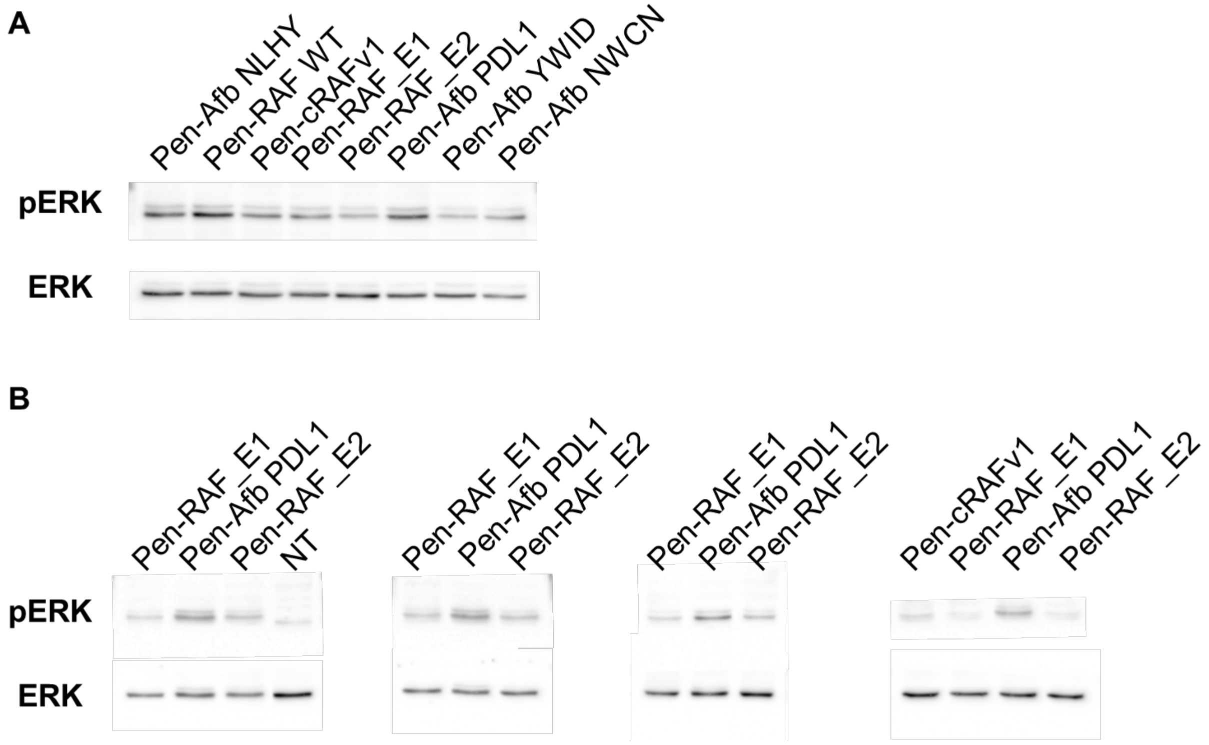

**Supplementary Figure 13. pERK Western Blots. A.** Single replicate of various inhibitors delivered as protein (15 μM). **B.** Replicate of select inhibitors delivered as protein (15 μM); used in quantification in **Fig. 3h**. These images came from three gels, see **Fig. S25** for full gel images. NT indicates that no CPP-protein was added and $KRas^{G12D}$ expression plasmid was not transfected.

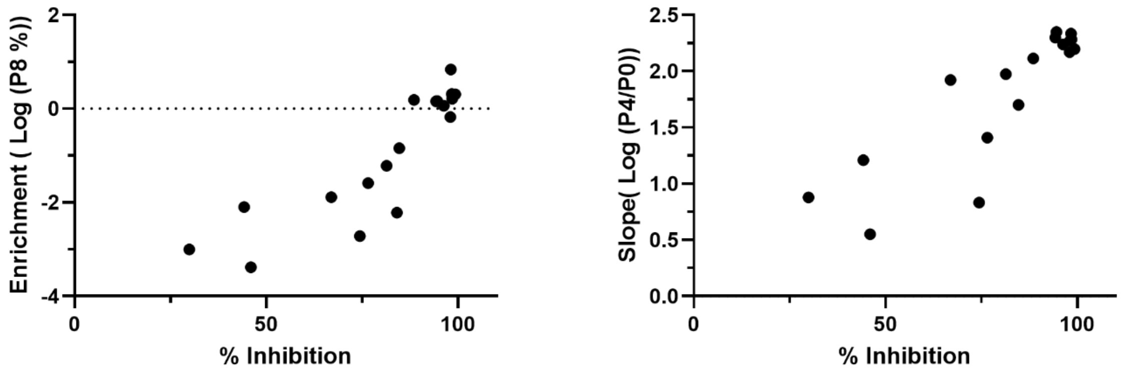

**Supplementary Figure 14. Comparing Inhibition to Enrichment (left) and Slope (right).** Enrichment in Passage 8, the means for picking variants to test in Fig. 3, is not corrected for differences in initial copy number in the library, but still is well correlated (Pearson's Correlation, $r = 0.79$, $p = <0.0001$) to measured Inhibition (100% = full inhibition). Slope from the library to Passage 4, the measure used throughout Fig. 4, is also well correlated (Pearson's Correlation, $r = 0.86$, $p = <0.0001$) to measured Inhibition (100% = full inhibition).

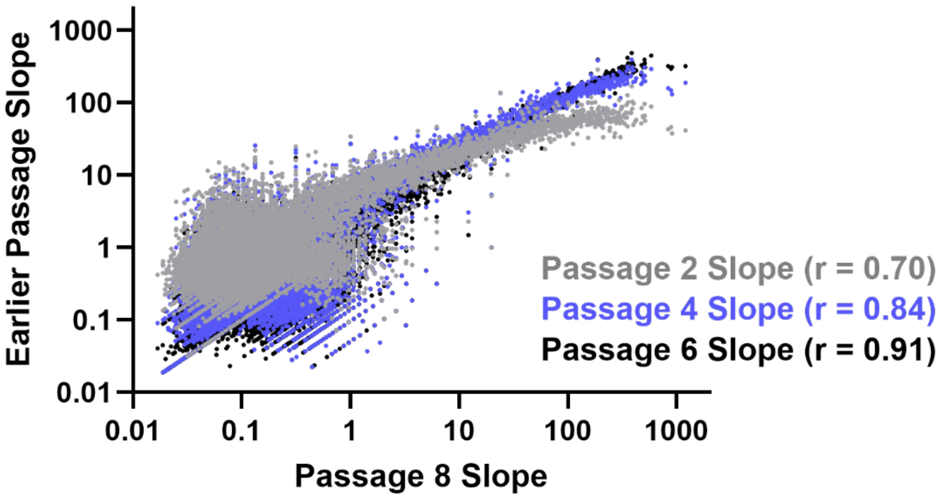

**Supplementary Figure 15. Choosing which passage to use for determining slope.** We see that as the selection proceeds from Library to Passage 2, 4, 6, and 8, we observe that the slope (Passage NGS% over Library NGS%) becomes more internally consistent. However, so do the total number of variants with reads dramatically declines between passage 4 and 6 (all variants with reads of 0 was set to 0.001%, which is at lower than 1 read as a %). Pearson's Correlation listed in legend.

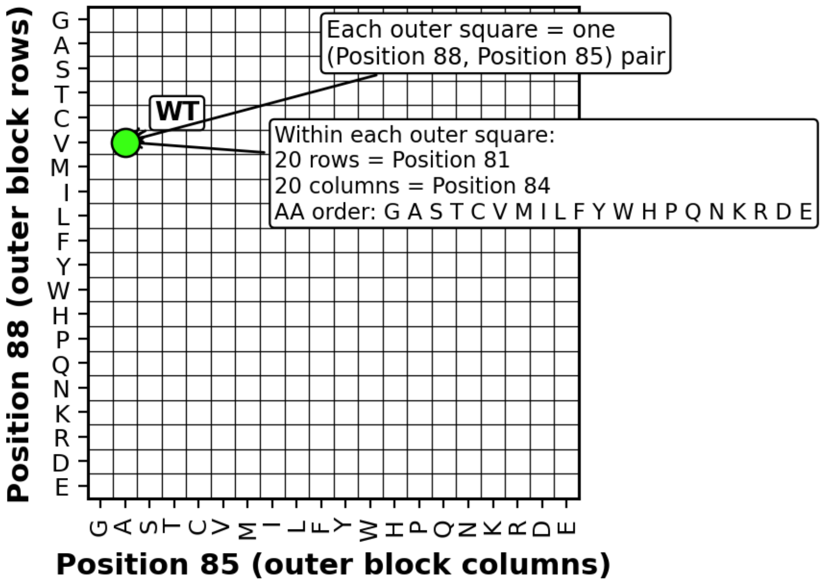

**Supplementary Figure 16. Schematic for layout of Fig. 4b fitness landscape.** The landscape is constructed as 400 squares of the 20x20 possible combination of mutations to positions 85 and 88 with each of these square being composed of an internal 400 squares of the 20x20 possible combination of mutations to positions 81 and 84. Each internal square is shaded based on the slope of that single protein variant.

### Supplementary Table 3. Hierarchical Clustering of Pen-Raf top 1000 enriched variants.

| Group1 (randomized positions indicated in blue in first sequence) | | Group70 | (* NGS reads in P8) |
|---|---|---|---|
| VRNGMSLHD**T**LM**KP**LK**Y**RGLQPECC | 259 | VRNGMSLHDTLMNKLKNRGLQPECC | 181* |
| VRNGMSLHDSLMKPLKYRGLQPECC | 78 | VRNGMSLHDSLMNKLKNRGLQPECC | 33 |
| VRNGMSLHDTLMRPLKYRGLQPECC | 15 | VRNGMSLHDALMNKLKNRGLQPECC | 279 |
| VRNGMSLHDALMKPLKYRGLQPECC | 156 | VRNGMSLHDMLMNKLKNRGLQPECC | 901 |
| VRNGMSLHDALMRPLKYRGLQPECC | 44 | VRNGMSLHDLLMNKLKNRGLQPECC | 809 |
| Group2 | | VRNGMSLHDVLMNKLKNRGLQPECC | 318 |
| VRNGMSLHDLLMQKLKYRGLQPECC | 236 | VRNGMSLHDILMNKLKNRGLQPECC | 191 |
| VRNGMSLHDMLMQKLKYRGLQPECC | 23 | Group71 | |
| VRNGMSLHDVLMQKLKYRGLQPECC | 141 | VRNGMSLHDVLMNKLKTRGLQPECC | 418 |
| VRNGMSLHDILMQKLKYRGLQPECC | 70 | VRNGMSLHDILMNKLKTRGLQPECC | 28 |
| Group3 | | VRNGMSLHDVLMNKLKSRGLQPECC | 10 |
| VRNGMSLHDALMWKLKNRGLQPECC | 33 | VRNGMSLHDLLMNKLKTRGLQPECC | 198 |
| VRNGMSLHDTLMWKLKNRGLQPECC | 13 | VRNGMSLHDMLMNKLKTRGLQPECC | 23 |
| VRNGMSLHDVLMWKLKNRGLQPECC | 23 | VRNGMSLHDLLMNKLKSRGLQPECC | 20 |
| VRNGMSLHDLLMWKLKNRGLQPECC | 13 | VRNGMSLHDLLMNKLKARGLQPECC | 20 |
| Group4 | | Group72 | |
| VRNGMSLHDLLMSKLKNRGLQPECC | 219 | VRNGMSLHDLLMMKLKYRGLQPECC | 360 |
| VRNGMSLHDMLMSKLKNRGLQPECC | 33 | VRNGMSLHDLLMLKLKYRGLQPECC | 304 |
| VRNGMSLHDLLMTKLKNRGLQPECC | 183 | VRNGMSLHDMLMLKLKYRGLQPECC | 74 |
| VRNGMSLHDMLMTKLKNRGLQPECC | 33 | VRNGMSLHDMLMMKLKYRGLQPECC | 54 |
| VRNGMSLHDILMSKLKNRGLQPECC | 281 | VRNGMSLHDVLMLKLKYRGLQPECC | 336 |
| VRNGMSLHDVLMSKLKNRGLQPECC | 261 | VRNGMSLHDILMLKLKYRGLQPECC | 128 |
| VRNGMSLHDVLMTKLKNRGLQPECC | 158 | VRNGMSLHDVLMMKLKYRGLQPECC | 286 |
| VRNGMSLHDILMTKLKNRGLQPECC | 64 | VRNGMSLHDILMMKLKYRGLQPECC | 174 |
| VRNGMSLHDLLMAKLKNRGLQPECC | 156 | VRNGMSLHDLLMVKLKYRGLQPECC | 196 |
| VRNGMSLHDMLMAKLKNRGLQPECC | 12 | VRNGMSLHDLLMIKLKYRGLQPECC | 160 |
| VRNGMSLHDVLMAKLKNRGLQPECC | 105 | VRNGMSLHDMLMVKLKYRGLQPECC | 14 |
| VRNGMSLHDILMAKLKNRGLQPECC | 15 | VRNGMSLHDMLMIKLKYRGLQPECC | 13 |
| Group5 | | VRNGMSLHDILMVKLKYRGLQPECC | 67 |
| VRNGMSLHDLLMKSLKNRGLQPECC | 53 | VRNGMSLHDILMIKLKYRGLQPECC | 34 |
| VRNGMSLHDMLMKSLKNRGLQPECC | 17 | VRNGMSLHDVLMVKLKYRGLQPECC | 342 |
| VRNGMSLHDVLMKSLKNRGLQPECC | 27 | VRNGMSLHDVLMIKLKYRGLQPECC | 188 |
| VRNGMSLHDILMKSLKNRGLQPECC | 9 | Group73 | |
| VRNGMSLHDLLMKALKNRGLQPECC | 86 | VRNGMSLHDALMLKLKYRGLQPECC | 322 |
| VRNGMSLHDMLMKALKNRGLQPECC | 17 | VRNGMSLHDALMMKLKYRGLQPECC | 228 |
| VRNGMSLHDVLMKALKNRGLQPECC | 24 | VRNGMSLHDALMVKLKYRGLQPECC | 123 |
| VRNGMSLHDILMKALKNRGLQPECC | 23 | VRNGMSLHDALMIKLKYRGLQPECC | 90 |
| Group6 | | VRNGMSLHDALMMKLKFRGLQPECC | 71 |
| VRNGMSLHDVLMIKLKMRGLQPECC | 9 | VRNGMSLHDALMLKLKFRGLQPECC | 55 |
| VRNGMSLHDVLMVKLKLRGLQPECC | 9 | VRNGMSLHDALMIKLKFRGLQPECC | 13 |
| VRNGMSLHDLLMIKLKLRGLQPECC | 9 | Group74 | |
| VRNGMSLHDLLMVKLKLRGLQPECC | 9 | VRNGMSLHDLLMMKLKNRGLQPECC | 147 |
| VRNGMSLHDLLMVKLKMRGLQPECC | 9 | VRNGMSLHDMLMMKLKNRGLQPECC | 10 |
| VRNGMSLHDLLMVKLKVRGLQPECC | 27 | VRNGMSLHDLLMLKLKNRGLQPECC | 68 |
| VRNGMSLHDVLMVKLKVRGLQPECC | 14 | VRNGMSLHDVLMLKLKNRGLQPECC | 130 |
| Group7 | | VRNGMSLHDILMLKLKNRGLQPECC | 20 |
| VRNGMSLHDVLMTKLKWRGLQPECC | 26 | VRNGMSLHDVLMMKLKNRGLQPECC | 88 |
| VRNGMSLHDVLMSKLKWRGLQPECC | 25 | VRNGMSLHDILMMKLKNRGLQPECC | 53 |
| VRNGMSLHDVLMAKLKWRGLQPECC | 12 | VRNGMSLHDLLMIKLKNRGLQPECC | 147 |
| VRNGMSLHDLLMSKLKWRGLQPECC | 29 | VRNGMSLHDLLMVKLKNRGLQPECC | 133 |
| VRNGMSLHDLLMTKLKWRGLQPECC | 29 | VRNGMSLHDMLMVKLKNRGLQPECC | 11 |
| Group8 | | VRNGMSLHDILMIKLKNRGLQPECC | 56 |
| VRNGMSLHDLLMPKLKTRGLQPECC | 125 | VRNGMSLHDILMVKLKNRGLQPECC | 48 |
| VRNGMSLHDMLMPKLKTRGLQPECC | 47 | VRNGMSLHDVLMVKLKNRGLQPECC | 332 |
| VRNGMSLHDLLMPKLKSRGLQPECC | 32 | VRNGMSLHDVLMIKLKNRGLQPECC | 79 |
| VRNGMSLHDVLMPKLKTRGLQPECC | 101 | Group75 | |
| VRNGMSLHDILMPKLKTRGLQPECC | 68 | VRNGMSLHDALMTKLKHRGLQPECC | 77 |
| VRNGMSLHDILMPKLKSRGLQPECC | 30 | VRNGMSLHDALMSKLKHRGLQPECC | 65 |
| VRNGMSLHDVLMPKLKSRGLQPECC | 15 | VRNGMSLHDALMAKLKHRGLQPECC | 25 |
| VRNGMSLHDLLMPKLKARGLQPECC | 51 | VRNGMSLHDALMLKLKHRGLQPECC | 121 |
| VRNGMSLHDMLMPKLKARGLQPECC | 11 | VRNGMSLHDALMMKLKHRGLQPECC | 70 |
| VRNGMSLHDILMPKLKARGLQPECC | 21 | VRNGMSLHDALMIKLKHRGLQPECC | 87 |
| VRNGMSLHDVLMPKLKARGLQPECC | 16 | VRNGMSLHDALMVKLKHRGLQPECC | 39 |
| Group9 | | Group76 | |

| | | | |
|---|---|---|---|
| VRNGMSLHDCLMSKLKVRGLQPECC | 11 | VRNGMSLHDTLMSKLKHRGLQPECC | 59 |
| VRNGMSLHDCLMSKLKIRGLQPECC | 9 | VRNGMSLHDTLMTKLKHRGLQPECC | 52 |
| VRNGMSLHDCLMTKLKVRGLQPECC | 21 | VRNGMSLHDTLMAKLKHRGLQPECC | 26 |
| VRNGMSLHDCLMTKLKIRGLQPECC | 15 | VRNGMSLHDTLMMKLKHRGLQPECC | 75 |
| VRNGMSLHDCLMAKLKVRGLQPECC | 18 | VRNGMSLHDTLMLKLKHRGLQPECC | 71 |
| VRNGMSLHDCLMAKLKIRGLQPECC | 11 | VRNGMSLHDTLMVKLKHRGLQPECC | 47 |
| VRNGMSLHDCLMTKLKMRGLQPECC | 17 | VRNGMSLHDTLMIKLKHRGLQPECC | 45 |
| VRNGMSLHDCLMTKLKLRGLQPECC | 12 | Group77 | |
| VRNGMSLHDCLMSKLKLRGLQPECC | 17 | VRNGMSLHDLLMLKLKHRGLQPECC | 135 |
| Group10 | | VRNGMSLHDLLMMKLKHRGLQPECC | 99 |
| VRNGMSLHDVLMCKLKYRGLQPECC | 198 | VRNGMSLHDMLMLKLKHRGLQPECC | 30 |
| VRNGMSLHDILMCKLKYRGLQPECC | 42 | VRNGMSLHDMLMMKLKHRGLQPECC | 16 |
| VRNGMSLHDLLMCKLKYRGLQPECC | 174 | VRNGMSLHDVLMLKLKHRGLQPECC | 184 |
| VRNGMSLHDTLMCKLKYRGLQPECC | 84 | VRNGMSLHDILMLKLKHRGLQPECC | 33 |
| VRNGMSLHDLLMCKLKFRGLQPECC | 75 | VRNGMSLHDVLMMKLKHRGLQPECC | 147 |
| VRNGMSLHDVLMCKLKFRGLQPECC | 10 | VRNGMSLHDILMMKLKHRGLQPECC | 86 |
| VRNGMSLHDALMCKLKYRGLQPECC | 148 | VRNGMSLHDLLMIKLKHRGLQPECC | 101 |
| VRNGMSLHDALMCKLKFRGLQPECC | 13 | VRNGMSLHDLLMVKLKHRGLQPECC | 97 |
| Group11 | | VRNGMSLHDMLMIKLKHRGLQPECC | 22 |
| VRNGMSLHDVLMKKLKARGLQPECC | 46 | VRNGMSLHDMLMVKLKHRGLQPECC | 14 |
| VRNGMSLHDILMKKLKARGLQPECC | 15 | VRNGMSLHDILMVKLKHRGLQPECC | 49 |
| VRNGMSLHDVLMRKLKARGLQPECC | 19 | VRNGMSLHDILMIKLKHRGLQPECC | 36 |
| VRNGMSLHDLLMKKLKARGLQPECC | 34 | VRNGMSLHDVLMVKLKHRGLQPECC | 157 |
| VRNGMSLHDLLMRKLKARGLQPECC | 18 | VRNGMSLHDVLMIKLKHRGLQPECC | 101 |
| Group12 | | Group78 | |
| VRNGMSLHDTLMKKLKHRGLQPECC | 194 | VRNGMSLHDTLMPKLKNRGLQPECC | 134 |
| VRNGMSLHDSLMKKLKHRGLQPECC | 27 | VRNGMSLHDSLMPKLKNRGLQPECC | 15 |
| VRNGMSLHDTLMRKLKHRGLQPECC | 32 | VRNGMSLHDALMPKLKNRGLQPECC | 50 |
| VRNGMSLHDLLMKKLKHRGLQPECC | 523 | VRNGMSLHDLLMPKLKNRGLQPECC | 254 |
| VRNGMSLHDMLMKKLKHRGLQPECC | 130 | VRNGMSLHDMLMPKLKNRGLQPECC | 108 |
| VRNGMSLHDVLMKKLKHRGLQPECC | 594 | VRNGMSLHDVLMPKLKNRGLQPECC | 241 |
| VRNGMSLHDILMKKLKHRGLQPECC | 203 | VRNGMSLHDILMPKLKNRGLQPECC | 181 |
| VRNGMSLHDLLMRKLKHRGLQPECC | 199 | Group79 | |
| VRNGMSLHDMLMRKLKHRGLQPECC | 17 | VRNGMSLHDTLMNKLKHRGLQPECC | 132 |
| VRNGMSLHDVLMRKLKHRGLQPECC | 168 | VRNGMSLHDSLMNKLKHRGLQPECC | 14 |
| VRNGMSLHDILMRKLKHRGLQPECC | 60 | VRNGMSLHDALMNKLKHRGLQPECC | 235 |
| Group13 | | VRNGMSLHDLLMNKLKHRGLQPECC | 382 |
| VRNGMSLHDLLMNKLKLRGLQPECC | 54 | VRNGMSLHDMLMNKLKHRGLQPECC | 56 |
| VRNGMSLHDLLMNKLKMRGLQPECC | 28 | VRNGMSLHDVLMNKLKHRGLQPECC | 253 |
| VRNGMSLHDVLMNKLKLRGLQPECC | 48 | VRNGMSLHDILMNKLKHRGLQPECC | 116 |
| VRNGMSLHDVLMNKLKMRGLQPECC | 16 | Group80 | |
| VRNGMSLHDLLMNKLKVRGLQPECC | 81 | VRNGMSLHDLLMKKLKLRGLQPECC | 65 |
| VRNGMSLHDLLMNKLKIRGLQPECC | 26 | VRNGMSLHDMLMKKLKLRGLQPECC | 10 |
| VRNGMSLHDMLMNKLKVRGLQPECC | 12 | VRNGMSLHDLLMKKLKMRGLQPECC | 25 |
| VRNGMSLHDVLMNKLKVRGLQPECC | 85 | VRNGMSLHDVLMKKLKLRGLQPECC | 98 |
| VRNGMSLHDVLMNKLKIRGLQPECC | 15 | VRNGMSLHDILMKKLKLRGLQPECC | 15 |
| VRNGMSLHDILMNKLKVRGLQPECC | 15 | VRNGMSLHDVLMKKLKMRGLQPECC | 25 |
| Group14 | | VRNGMSLHDILMKKLKMRGLQPECC | 12 |
| VRNGMSLHDCLMTKLKFRGLQPECC | 59 | VRNGMSLHDLLMKKLKVRGLQPECC | 141 |
| VRNGMSLHDCLMSKLKFRGLQPECC | 53 | VRNGMSLHDLLMKKLKIRGLQPECC | 36 |
| VRNGMSLHDCLMNKLKFRGLQPECC | 268 | VRNGMSLHDMLMKKLKVRGLQPECC | 10 |
| VRNGMSLHDCLMKKLKFRGLQPECC | 163 | VRNGMSLHDILMKKLKVRGLQPECC | 44 |
| VRNGMSLHDCLMRKLKFRGLQPECC | 95 | VRNGMSLHDILMKKLKIRGLQPECC | 10 |
| Group15 | | VRNGMSLHDVLMKKLKVRGLQPECC | 214 |
| VRNGMSLHDLLMKKLKQRGLQPECC | 184 | VRNGMSLHDVLMKKLKIRGLQPECC | 69 |
| VRNGMSLHDMLMKKLKQRGLQPECC | 52 | Group81 | |
| VRNGMSLHDILMKKLKQRGLQPECC | 178 | VRNGMSLHDTLMPKLKTRGLQPECC | 54 |
| VRNGMSLHDVLMKKLKQRGLQPECC | 116 | VRNGMSLHDTLMPKLKSRGLQPECC | 9 |
| VRNGMSLHDTLMKKLKQRGLQPECC | 41 | VRNGMSLHDTLMPKLKARGLQPECC | 23 |
| VRNGMSLHDLLMRKLKQRGLQPECC | 62 | VRNGMSLHDTLMPKLKVRGLQPECC | 28 |
| VRNGMSLHDVLMRKLKQRGLQPECC | 24 | VRNGMSLHDTLMPKLKLRGLQPECC | 10 |
| VRNGMSLHDALMKKLKQRGLQPECC | 45 | Group82 | |
| VRNGMSLHDALMRKLKQRGLQPECC | 22 | VRNGMSLHDTLMRKLKYRGLQPECC | 122 |
| Group16 | | VRNGMSLHDSLMRKLKYRGLQPECC | 20 |
| VRNGMSLHDTLMSKLKYRGLQPECC | 76 | VRNGMSLHDALMRKLKYRGLQPECC | 475 |
| VRNGMSLHDTLMAKLKYRGLQPECC | 56 | VRNGMSLHDTLMKKLKYRGLQPECC | 542 |
| VRNGMSLHDTLMTKLKYRGLQPECC | 99 | VRNGMSLHDSLMKKLKYRGLQPECC | 62 |

| | | | |
|---|---|---|---|
| VRNGMSLHDSLMTKLKYRGLQPECC | 20 | Group83 | |
| VRNGMSLHDALMTKLKYRGLQPECC | 173 | VRNGMSLHDILMPKLKLRGLQPECC | 43 |
| VRNGMSLHDALMSKLKYRGLQPECC | 117 | VRNGMSLHDVLMPKLKLRGLQPECC | 15 |
| VRNGMSLHDALMAKLKYRGLQPECC | 170 | VRNGMSLHDILMPKLKMRGLQPECC | 13 |
| Group17 | | VRNGMSLHDLLMPKLKLRGLQPECC | 46 |
| VRNGMSLHDALMSKLKTRGLQPECC | 13 | VRNGMSLHDLLMPKLKMRGLQPECC | 14 |
| VRNGMSLHDALMTKLKTRGLQPECC | 9 | VRNGMSLHDLLMPKLKVRGLQPECC | 80 |
| VRNGMSLHDALMNKLKTRGLQPECC | 42 | VRNGMSLHDLLMPKLKIRGLQPECC | 23 |
| VRNGMSLHDALMVKLKTRGLQPECC | 16 | VRNGMSLHDMLMPKLKVRGLQPECC | 15 |
| VRNGMSLHDALMIKLKTRGLQPECC | 13 | VRNGMSLHDILMPKLKIRGLQPECC | 23 |
| VRNGMSLHDALMMKLKTRGLQPECC | 13 | VRNGMSLHDVLMPKLKIRGLQPECC | 12 |
| Group18 | | VRNGMSLHDILMPKLKVRGLQPECC | 90 |
| VRNGMSLHDCLMKPLKYRGLQPECC | 209 | VRNGMSLHDVLMPKLKVRGLQPECC | 60 |
| VRNGMSLHDCLMRPLKYRGLQPECC | 58 | Group84 | |
| VRNGMSLHDCLMKPLKFRGLQPECC | 37 | VRNGMSLHDLLMKKLKNRGLQPECC | 338 |
| VRNGMSLHDCLMRPLKFRGLQPECC | 10 | VRNGMSLHDMLMKKLKNRGLQPECC | 166 |
| Group19 | | VRNGMSLHDVLMKKLKNRGLQPECC | 352 |
| VRNGMSLHDLLMKALKYRGLQPECC | 224 | VRNGMSLHDILMKKLKNRGLQPECC | 216 |
| VRNGMSLHDMLMKALKYRGLQPECC | 24 | VRNGMSLHDLLMRKLKNRGLQPECC | 158 |
| VRNGMSLHDVLMKALKYRGLQPECC | 44 | VRNGMSLHDMLMRKLKNRGLQPECC | 29 |
| VRNGMSLHDILMKALKYRGLQPECC | 19 | VRNGMSLHDVLMRKLKNRGLQPECC | 177 |
| VRNGMSLHDTLMKALKYRGLQPECC | 59 | VRNGMSLHDILMRKLKNRGLQPECC | 58 |
| VRNGMSLHDSLMKALKYRGLQPECC | 25 | Group85 | |
| Group20 | | VRNGMSLHDLLMKKLKYRGLQPECC | 395 |
| VRNGMSLHDVLMFKLKIRGLQPECC | 16 | VRNGMSLHDMLMKKLKYRGLQPECC | 249 |
| VRNGMSLHDVLMFKLKVRGLQPECC | 13 | VRNGMSLHDVLMKKLKYRGLQPECC | 426 |
| VRNGMSLHDVLMYKLKVRGLQPECC | 10 | VRNGMSLHDILMKKLKYRGLQPECC | 246 |
| VRNGMSLHDLLMFKLKVRGLQPECC | 56 | VRNGMSLHDLLMRKLKYRGLQPECC | 446 |
| Group21 | | VRNGMSLHDMLMRKLKYRGLQPECC | 383 |
| VRNGMSLHDTLMMKLKYRGLQPECC | 430 | VRNGMSLHDVLMRKLKYRGLQPECC | 310 |
| VRNGMSLHDTLMLKLKYRGLQPECC | 188 | VRNGMSLHDILMRKLKYRGLQPECC | 106 |
| VRNGMSLHDTLMVKLKYRGLQPECC | 139 | Group86 | |
| VRNGMSLHDTLMIKLKYRGLQPECC | 26 | VRNGMSLHDALMQKLKNRGLQPECC | 79 |
| VRNGMSLHDSLMLKLKYRGLQPECC | 33 | VRNGMSLHDTLMQKLKNRGLQPECC | 41 |
| VRNGMSLHDSLMMKLKYRGLQPECC | 17 | VRNGMSLHDLLMQKLKNRGLQPECC | 102 |
| Group22 | | VRNGMSLHDMLMQKLKNRGLQPECC | 14 |
| VRNGMSLHDVLMMKLKTRGLQPECC | 25 | VRNGMSLHDVLMQKLKNRGLQPECC | 93 |
| VRNGMSLHDVLMLKLKTRGLQPECC | 15 | VRNGMSLHDILMQKLKNRGLQPECC | 31 |
| VRNGMSLHDVLMVKLKTRGLQPECC | 14 | Group87 | |
| VRNGMSLHDVLMIKLKTRGLQPECC | 12 | VRNGMSLHDALMKKLKLRGLQPECC | 64 |
| Group23 | | VRNGMSLHDALMKKLKMRGLQPECC | 15 |
| VRNGMSLHDVLMFKLKYRGLQPECC | 105 | VRNGMSLHDALMKKLKVRGLQPECC | 111 |
| VRNGMSLHDILMFKLKYRGLQPECC | 11 | VRNGMSLHDALMKKLKIRGLQPECC | 33 |
| VRNGMSLHDLLMFKLKYRGLQPECC | 186 | VRNGMSLHDALMRKLKVRGLQPECC | 33 |
| VRNGMSLHDALMFKLKYRGLQPECC | 130 | VRNGMSLHDALMRKLKIRGLQPECC | 15 |
| VRNGMSLHDLLMYKLKYRGLQPECC | 133 | VRNGMSLHDALMRKLKLRGLQPECC | 13 |
| VRNGMSLHDMLMYKLKYRGLQPECC | 16 | Group88 | |
| VRNGMSLHDVLMYKLKYRGLQPECC | 169 | VRNGMSLHDLLMKPLKYRGLQPECC | 334 |
| VRNGMSLHDILMYKLKYRGLQPECC | 108 | VRNGMSLHDMLMKPLKYRGLQPECC | 332 |
| Group24 | | VRNGMSLHDVLMKPLKYRGLQPECC | 541 |
| VRNGMSLHDLLMTKLKYRGLQPECC | 307 | VRNGMSLHDILMKPLKYRGLQPECC | 286 |
| VRNGMSLHDMLMTKLKYRGLQPECC | 42 | VRNGMSLHDLLMRPLKYRGLQPECC | 102 |
| VRNGMSLHDLLMSKLKYRGLQPECC | 196 | VRNGMSLHDMLMRPLKYRGLQPECC | 44 |
| VRNGMSLHDMLMSKLKYRGLQPECC | 73 | VRNGMSLHDVLMRPLKYRGLQPECC | 63 |
| VRNGMSLHDVLMTKLKYRGLQPECC | 253 | VRNGMSLHDILMRPLKYRGLQPECC | 57 |
| VRNGMSLHDILMTKLKYRGLQPECC | 125 | Group89 | |
| VRNGMSLHDVLMSKLKYRGLQPECC | 252 | VRNGMSLHDALMHKLKNRGLQPECC | 66 |
| VRNGMSLHDILMSKLKYRGLQPECC | 162 | VRNGMSLHDTLMHKLKNRGLQPECC | 37 |
| Group25 | | VRNGMSLHDLLMHKLKNRGLQPECC | 146 |
| VRNGMSLHDVLMSKLKVRGLQPECC | 40 | VRNGMSLHDMLMHKLKNRGLQPECC | 27 |
| VRNGMSLHDVLMTKLKVRGLQPECC | 36 | VRNGMSLHDVLMHKLKNRGLQPECC | 164 |
| VRNGMSLHDLLMSKLKVRGLQPECC | 35 | VRNGMSLHDILMHKLKNRGLQPECC | 83 |
| VRNGMSLHDLLMTKLKVRGLQPECC | 24 | Group90 | |
| VRNGMSLHDLLMAKLKVRGLQPECC | 16 | VRNGMSLHDALMQKLKHRGLQPECC | 66 |
| VRNGMSLHDVLMAKLKVRGLQPECC | 9 | VRNGMSLHDTLMQKLKHRGLQPECC | 27 |
| Group26 | | VRNGMSLHDLLMQKLKHRGLQPECC | 112 |
| VRNGMSLHDMLMNKLKYRGLQPECC | 421 | VRNGMSLHDMLMQKLKHRGLQPECC | 9 |

| | | | |
|---|---|---|---|
| VRNGMSLHDLLMNKLKYRGLQPECC | 413 | VRNGMSLHDVLMQKLKHRGLQPECC | 62 |
| VRNGMSLHDVLMNKLKYRGLQPECC | 305 | VRNGMSLHDILMQKLKHRGLQPECC | 10 |
| VRNGMSLHDILMNKLKYRGLQPECC | 180 | Group91 | |
| VRNGMSLHDLLMNKLKFRGLQPECC | 266 | VRNGMSLHDLLMYKLKWRGLQPECC | 99 |
| VRNGMSLHDMLMNKLKFRGLQPECC | 34 | VRNGMSLHDLLMFKLKWRGLQPECC | 13 |
| VRNGMSLHDVLMNKLKFRGLQPECC | 271 | VRNGMSLHDVLMYKLKWRGLQPECC | 55 |
| VRNGMSLHDILMNKLKFRGLQPECC | 55 | VRNGMSLHDVLMFKLKWRGLQPECC | 22 |
| Group27 | | Group92 | |
| VRNGMSLHDCLMHKLKTRGLQPECC | 93 | VRNGMSLHDALMHKLKHRGLQPECC | 46 |
| VRNGMSLHDCLMHKLKVRGLQPECC | 20 | VRNGMSLHDTLMHKLKHRGLQPECC | 26 |
| VRNGMSLHDCLMHKLKYRGLQPECC | 154 | VRNGMSLHDLLMHKLKHRGLQPECC | 96 |
| VRNGMSLHDCLMHKLKFRGLQPECC | 10 | VRNGMSLHDMLMHKLKHRGLQPECC | 9 |
| Group28 | | VRNGMSLHDVLMHKLKHRGLQPECC | 71 |
| VRNGMSLHDCLMKKLKTRGLQPECC | 92 | VRNGMSLHDILMHKLKHRGLQPECC | 15 |
| VRNGMSLHDCLMKKLKSRGLQPECC | 17 | Group93 | |
| VRNGMSLHDCLMRKLKTRGLQPECC | 59 | VRNGMSLHDTLMPKLKHRGLQPECC | 112 |
| VRNGMSLHDCLMRKLKSRGLQPECC | 12 | VRNGMSLHDALMPKLKHRGLQPECC | 49 |
| VRNGMSLHDCLMRKLKARGLQPECC | 16 | VRNGMSLHDLLMPKLKHRGLQPECC | 250 |
| VRNGMSLHDCLMKKLKARGLQPECC | 15 | VRNGMSLHDMLMPKLKHRGLQPECC | 227 |
| Group29 | | VRNGMSLHDILMPKLKHRGLQPECC | 177 |
| VRNGMSLHDCLMMKLKYRGLQPECC | 409 | VRNGMSLHDVLMPKLKHRGLQPECC | 143 |
| VRNGMSLHDCLMLKLKYRGLQPECC | 396 | Group94 | |
| VRNGMSLHDCLMVKLKYRGLQPECC | 144 | VRNGMSLHDVLMGKLKHRGLQPECC | 26 |
| VRNGMSLHDCLMIKLKYRGLQPECC | 76 | VRNGMSLHDILMGKLKHRGLQPECC | 10 |
| VRNGMSLHDCLMLKLKFRGLQPECC | 126 | VRNGMSLHDLLMGKLKHRGLQPECC | 49 |
| VRNGMSLHDCLMMKLKFRGLQPECC | 21 | VRNGMSLHDALMGKLKHRGLQPECC | 24 |
| Group30 | | Group95 | |
| VRNGMSLHDALMKKLKTRGLQPECC | 81 | VRNGMSLHDALMNKLKWRGLQPECC | 272 |
| VRNGMSLHDALMKKLKSRGLQPECC | 16 | VRNGMSLHDTLMNKLKWRGLQPECC | 265 |
| VRNGMSLHDALMRKLKTRGLQPECC | 41 | VRNGMSLHDLLMNKLKWRGLQPECC | 260 |
| VRNGMSLHDALMRKLKSRGLQPECC | 12 | VRNGMSLHDMLMNKLKWRGLQPECC | 27 |
| VRNGMSLHDALMKKLKARGLQPECC | 16 | VRNGMSLHDVLMNKLKWRGLQPECC | 256 |
| VRNGMSLHDALMRKLKARGLQPECC | 11 | VRNGMSLHDILMNKLKWRGLQPECC | 48 |
| Group31 | | Group96 | |
| VRNGMSLHDLLMRKLKLRGLQPECC | 16 | VRNGMSLHDCLMKKLKLRGLQPECC | 59 |
| VRNGMSLHDLLMRKLKMRGLQPECC | 12 | VRNGMSLHDCLMKKLKMRGLQPECC | 22 |
| VRNGMSLHDLLMRKLKVRGLQPECC | 34 | VRNGMSLHDCLMKKLKVRGLQPECC | 104 |
| VRNGMSLHDVLMRKLKVRGLQPECC | 52 | VRNGMSLHDCLMKKLKIRGLQPECC | 55 |
| VRNGMSLHDVLMRKLKIRGLQPECC | 12 | VRNGMSLHDCLMRKLKVRGLQPECC | 36 |
| VRNGMSLHDILMRKLKVRGLQPECC | 14 | VRNGMSLHDCLMRKLKIRGLQPECC | 17 |
| VRNGMSLHDVLMRKLKLRGLQPECC | 24 | VRNGMSLHDCLMRKLKLRGLQPECC | 15 |
| Group32 | | Group97 | |
| VRNGMSLHDCLMPKLKTRGLQPECC | 21 | VRNGMSLHDALMGKLKYRGLQPECC | 55 |
| VRNGMSLHDCLMPKLKARGLQPECC | 10 | VRNGMSLHDTLMGKLKYRGLQPECC | 12 |
| VRNGMSLHDCLMPKLKVRGLQPECC | 58 | VRNGMSLHDVLMGKLKYRGLQPECC | 26 |
| VRNGMSLHDCLMPKLKYRGLQPECC | 161 | VRNGMSLHDILMGKLKYRGLQPECC | 9 |
| VRNGMSLHDCLMPKLKFRGLQPECC | 30 | VRNGMSLHDLLMGKLKYRGLQPECC | 109 |
| Group33 | | Group98 | |
| VRNGMSLHDTLMPKLKYRGLQPECC | 141 | VRNGMSLHDLLMMKLKWRGLQPECC | 23 |
| VRNGMSLHDSLMPKLKYRGLQPECC | 14 | VRNGMSLHDMLMMKLKWRGLQPECC | 9 |
| VRNGMSLHDLLMPKLKYRGLQPECC | 242 | VRNGMSLHDLLMLKLKWRGLQPECC | 29 |
| VRNGMSLHDMLMPKLKYRGLQPECC | 156 | VRNGMSLHDVLMLKLKWRGLQPECC | 40 |
| VRNGMSLHDILMPKLKYRGLQPECC | 488 | VRNGMSLHDVLMMKLKWRGLQPECC | 18 |
| VRNGMSLHDVLMPKLKYRGLQPECC | 317 | Group99 | |
| VRNGMSLHDLLMPKLKFRGLQPECC | 144 | VRNGMSLHDALMCKLKNRGLQPECC | 37 |
| VRNGMSLHDMLMPKLKFRGLQPECC | 13 | VRNGMSLHDTLMCKLKNRGLQPECC | 16 |
| VRNGMSLHDVLMPKLKFRGLQPECC | 92 | VRNGMSLHDVLMCKLKNRGLQPECC | 80 |
| VRNGMSLHDILMPKLKFRGLQPECC | 45 | VRNGMSLHDILMCKLKNRGLQPECC | 19 |
| Group34 | | VRNGMSLHDLLMCKLKNRGLQPECC | 65 |
| VRNGMSLHDCLMKKLKNRGLQPECC | 386 | Group100 | |
| VRNGMSLHDCLMRKLKNRGLQPECC | 204 | VRNGMSLHDTLMWKLKHRGLQPECC | 89 |
| VRNGMSLHDCLMQKLKNRGLQPECC | 107 | VRNGMSLHDALMWKLKHRGLQPECC | 21 |
| VRNGMSLHDCLMSKLKNRGLQPECC | 349 | VRNGMSLHDVLMWKLKHRGLQPECC | 46 |
| VRNGMSLHDCLMTKLKNRGLQPECC | 143 | VRNGMSLHDILMWKLKHRGLQPECC | 26 |
| VRNGMSLHDCLMAKLKNRGLQPECC | 130 | VRNGMSLHDLLMWKLKHRGLQPECC | 58 |
| Group35 | | Group101 | |
| VRNGMSLHDCLMQKLKHRGLQPECC | 59 | VRNGMSLHDVLMTKLKFRGLQPECC | 26 |

| | | | |
|---|---|---|---|
| VRNGMSLHDCLMEKLKHRGLQPECC | 56 | VRNGMSLHDILMTKLKFRGLQPECC | 11 |
| VRNGMSLHDCLMKKLKHRGLQPECC | 337 | VRNGMSLHDVLMSKLKFRGLQPECC | 40 |
| VRNGMSLHDCLMRKLKHRGLQPECC | 103 | VRNGMSLHDLLMTKLKFRGLQPECC | 42 |
| VRNGMSLHDCLMSKLKHRGLQPECC | 148 | VRNGMSLHDLLMSKLKFRGLQPECC | 40 |
| VRNGMSLHDCLMTKLKHRGLQPECC | 142 | Group102 | |
| VRNGMSLHDCLMAKLKHRGLQPECC | 63 | VRNGMSLHDALMWKLKWRGLQPECC | 45 |
| VRNGMSLHDCLMNKLKHRGLQPECC | 205 | VRNGMSLHDTLMWKLKWRGLQPECC | 35 |
| VRNGMSLHDCLMDKLKHRGLQPECC | 11 | VRNGMSLHDLLMWKLKWRGLQPECC | 16 |
| Group36 | | VRNGMSLHDVLMWKLKWRGLQPECC | 16 |
| VRNGMSLHDLLMMKLKVRGLQPECC | 27 | Group103 | |
| VRNGMSLHDLLMMKLKIRGLQPECC | 13 | VRNGMSLHDCLMMKLKVRGLQPECC | 108 |
| VRNGMSLHDLLMMKLKLRGLQPECC | 29 | VRNGMSLHDCLMMKLKIRGLQPECC | 15 |
| VRNGMSLHDLLMMKLKTRGLQPECC | 38 | VRNGMSLHDCLMLKLKIRGLQPECC | 14 |
| VRNGMSLHDVLMMKLKVRGLQPECC | 23 | VRNGMSLHDCLMMKLKLRGLQPECC | 10 |
| VRNGMSLHDVLMMKLKLRGLQPECC | 15 | Group104 | |
| Group37 | | VRNGMSLHDLLMKPLKRRGLQPECC | 43 |
| VRNGMSLHDILMKPLKLRGLQPECC | 62 | VRNGMSLHDMLMKPLKRRGLQPECC | 11 |
| VRNGMSLHDVLMKPLKLRGLQPECC | 15 | VRNGMSLHDILMKPLKRRGLQPECC | 66 |
| VRNGMSLHDMLMKPLKLRGLQPECC | 14 | VRNGMSLHDVLMKPLKRRGLQPECC | 23 |
| VRNGMSLHDLLMKPLKLRGLQPECC | 36 | Group105 | |
| VRNGMSLHDLLMKPLKMRGLQPECC | 13 | VRNGMSLHDCLMMKLKNRGLQPECC | 185 |
| VRNGMSLHDLLMKPLKVRGLQPECC | 14 | VRNGMSLHDCLMLKLKNRGLQPECC | 89 |
| VRNGMSLHDILMKPLKVRGLQPECC | 11 | VRNGMSLHDCLMIKLKNRGLQPECC | 78 |
| VRNGMSLHDMLMRPLKLRGLQPECC | 15 | VRNGMSLHDCLMVKLKNRGLQPECC | 75 |
| VRNGMSLHDLLMRPLKLRGLQPECC | 11 | Group106 | |
| VRNGMSLHDILMRPLKLRGLQPECC | 12 | VRNGMSLHDCLMMKLKHRGLQPECC | 305 |
| Group38 | | VRNGMSLHDCLMLKLKHRGLQPECC | 243 |
| VRNGMSLHDLLMKPLKHRGLQPECC | 173 | VRNGMSLHDCLMIKLKHRGLQPECC | 136 |
| VRNGMSLHDMLMKPLKHRGLQPECC | 51 | VRNGMSLHDCLMVKLKHRGLQPECC | 107 |
| VRNGMSLHDVLMKPLKHRGLQPECC | 204 | Group107 | |
| VRNGMSLHDILMKPLKHRGLQPECC | 131 | VRNGMSLHDCLMLKLKWRGLQPECC | 57 |
| VRNGMSLHDLLMRPLKHRGLQPECC | 47 | VRNGMSLHDCLMMKLKWRGLQPECC | 43 |
| VRNGMSLHDMLMRPLKHRGLQPECC | 10 | VRNGMSLHDCLMVKLKWRGLQPECC | 16 |
| VRNGMSLHDILMRPLKHRGLQPECC | 24 | VRNGMSLHDCLMIKLKWRGLQPECC | 13 |
| VRNGMSLHDVLMRPLKHRGLQPECC | 17 | Group108 | |
| VRNGMSLHDTLMKPLKHRGLQPECC | 38 | VRNGMSLHDALMKALKYRGLQPECC | 41 |
| VRNGMSLHDSLMKPLKHRGLQPECC | 33 | VRNGMSLHDALMKSLKYRGLQPECC | 36 |
| VRNGMSLHDALMKPLKHRGLQPECC | 40 | VRNGMSLHDALMKGLKYRGLQPECC | 14 |
| Group39 | | VRNGMSLHDALMKKLKYRGLQPECC | 528 |
| VRNGMSLHDALMKKLKDRGLQPECC | 14 | VRNGMSLHDALMKRLKYRGLQPECC | 15 |
| VRNGMSLHDTLMKKLKDRGLQPECC | 10 | Group109 | |
| VRNGMSLHDVLMKKLKDRGLQPECC | 19 | VRNGMSLHDTLMKKLKTRGLQPECC | 25 |
| VRNGMSLHDILMKKLKDRGLQPECC | 10 | VRNGMSLHDTLMKKLKARGLQPECC | 9 |
| VRNGMSLHDLLMKKLKDRGLQPECC | 32 | VRNGMSLHDTLMKKLKFRGLQPECC | 46 |
| VRNGMSLHDLLMRKLKDRGLQPECC | 15 | VRNGMSLHDTLMKKLKLRGLQPECC | 16 |
| VRNGMSLHDMLMRKLKDRGLQPECC | 10 | Group110 | |
| Group40 | | VRNGMSLHDALMHKLKTRGLQPECC | 12 |
| VRNGMSLHDTLMKPLKNRGLQPECC | 50 | VRNGMSLHDALMHKLKVRGLQPECC | 10 |
| VRNGMSLHDSLMKPLKNRGLQPECC | 20 | VRNGMSLHDALMHKLKYRGLQPECC | 180 |
| VRNGMSLHDTLMRPLKNRGLQPECC | 26 | VRNGMSLHDALMHKLKFRGLQPECC | 59 |
| VRNGMSLHDLLMKPLKNRGLQPECC | 190 | Group111 | |
| VRNGMSLHDMLMKPLKNRGLQPECC | 95 | VRNGMSLHDALMMKLKVRGLQPECC | 40 |
| VRNGMSLHDILMKPLKNRGLQPECC | 166 | VRNGMSLHDALMIKLKVRGLQPECC | 17 |
| VRNGMSLHDVLMKPLKNRGLQPECC | 87 | VRNGMSLHDALMMKLKLRGLQPECC | 12 |
| VRNGMSLHDLLMRPLKNRGLQPECC | 80 | VRNGMSLHDALMIKLKMRGLQPECC | 13 |
| VRNGMSLHDMLMRPLKNRGLQPECC | 35 | VRNGMSLHDALMVKLKMRGLQPECC | 9 |
| VRNGMSLHDILMRPLKNRGLQPECC | 111 | Group112 | |
| VRNGMSLHDVLMRPLKNRGLQPECC | 39 | VRNGMSLHDLLMGKLKTRGLQPECC | 48 |
| Group41 | | VRNGMSLHDLLMGKLKSRGLQPECC | 10 |
| VRNGMSLHDILMKALKHRGLQPECC | 13 | VRNGMSLHDLLMGKLKARGLQPECC | 13 |
| VRNGMSLHDVLMKALKHRGLQPECC | 12 | VRNGMSLHDVLMGKLKTRGLQPECC | 23 |
| VRNGMSLHDLLMKALKHRGLQPECC | 49 | Group113 | |
| VRNGMSLHDTLMKALKHRGLQPECC | 16 | VRNGMSLHDVLMDKLKNRGLQPECC | 26 |
| VRNGMSLHDLLMKSLKHRGLQPECC | 36 | VRNGMSLHDILMDKLKNRGLQPECC | 15 |
| VRNGMSLHDILMKSLKHRGLQPECC | 14 | VRNGMSLHDLLMDKLKNRGLQPECC | 90 |
| Group42 | | VRNGMSLHDCLMDKLKNRGLQPECC | 18 |
| VRNGMSLHDTLMYKLKNRGLQPECC | 9 | Group114 | |

| | | | |
|---|---|---|---|
| VRNGMSLHDTLMFKLKNRGLQPECC | 9 | VRNGMSLHDILMPKLKWRGLQPECC | 99 |
| VRNGMSLHDLLMYKLKNRGLQPECC | 150 | VRNGMSLHDVLMPKLKWRGLQPECC | 27 |
| VRNGMSLHDMLMYKLKNRGLQPECC | 9 | VRNGMSLHDLLMPKLKWRGLQPECC | 45 |
| VRNGMSLHDVLMYKLKNRGLQPECC | 76 | VRNGMSLHDCLMPKLKWRGLQPECC | 16 |
| VRNGMSLHDILMYKLKNRGLQPECC | 13 | Group115 | |
| VRNGMSLHDVLMFKLKNRGLQPECC | 33 | VRNGMSLHDALMKALKHRGLQPECC | 26 |
| VRNGMSLHDILMFKLKNRGLQPECC | 17 | VRNGMSLHDALMKSLKHRGLQPECC | 9 |
| VRNGMSLHDLLMFKLKNRGLQPECC | 66 | VRNGMSLHDALMKKLKHRGLQPECC | 276 |
| Group43 | | VRNGMSLHDALMRKLKHRGLQPECC | 178 |
| VRNGMSLHDLLMDKLKYRGLQPECC | 213 | Group116 | |
| VRNGMSLHDMLMDKLKYRGLQPECC | 24 | VRNGMSLHDLLMHKLKVRGLQPECC | 24 |
| VRNGMSLHDVLMDKLKYRGLQPECC | 94 | VRNGMSLHDVLMHKLKVRGLQPECC | 16 |
| VRNGMSLHDILMDKLKYRGLQPECC | 41 | VRNGMSLHDLLMHKLKLRGLQPECC | 10 |
| VRNGMSLHDTLMDKLKYRGLQPECC | 67 | VRNGMSLHDVLMHKLKTRGLQPECC | 72 |
| VRNGMSLHDLLMDKLKFRGLQPECC | 53 | VRNGMSLHDLLMHKLKTRGLQPECC | 26 |
| VRNGMSLHDVLMDKLKFRGLQPECC | 38 | Group117 | |
| Group44 | | VRNGMSLHDALMMKLKWRGLQPECC | 83 |
| VRNGMSLHDLLMQKLKVRGLQPECC | 36 | VRNGMSLHDALMLKLKWRGLQPECC | 42 |
| VRNGMSLHDLLMQKLKIRGLQPECC | 10 | VRNGMSLHDALMVKLKWRGLQPECC | 15 |
| VRNGMSLHDMLMQKLKVRGLQPECC | 54 | VRNGMSLHDALMYKLKWRGLQPECC | 48 |
| VRNGMSLHDVLMQKLKVRGLQPECC | 20 | VRNGMSLHDALMFKLKWRGLQPECC | 26 |
| VRNGMSLHDLLMQKLKLRGLQPECC | 14 | Group118 | |
| VRNGMSLHDVLMQKLKLRGLQPECC | 12 | VRNGMSLHDTLMKKLKNRGLQPECC | 110 |
| Group45 | | VRNGMSLHDSLMKKLKNRGLQPECC | 11 |
| VRNGMSLHDALMKKLKFRGLQPECC | 424 | VRNGMSLHDTLMRKLKNRGLQPECC | 92 |
| VRNGMSLHDALMRKLKFRGLQPECC | 74 | VRNGMSLHDALMRKLKNRGLQPECC | 223 |
| VRNGMSLHDALMSKLKFRGLQPECC | 30 | VRNGMSLHDALMKKLKNRGLQPECC | 187 |
| VRNGMSLHDALMTKLKFRGLQPECC | 27 | GroupMIXED | |
| Group46 | | VRNGMSLHDTLMYKLKYRGLQPECC | 62 |
| VRNGMSLHDALMKKLKWRGLQPECC | 189 | VRNGMSLHDTLMFKLKYRGLQPECC | 48 |
| VRNGMSLHDALMRKLKWRGLQPECC | 159 | VRNGMSLHDYLMAELKIRGLQPECC | 12 |
| VRNGMSLHDALMQKLKWRGLQPECC | 19 | VRNGMSLHDILMTILKCRGLQPECC | 10 |
| VRNGMSLHDALMTKLKWRGLQPECC | 52 | VRNGMSLHDLLMDKLKWRGLQPECC | 15 |
| VRNGMSLHDALMSKLKWRGLQPECC | 13 | VRNGMSLHDVLMDKLKWRGLQPECC | 15 |
| VRNGMSLHDALMAKLKWRGLQPECC | 13 | VRNGMSLHDALMDKLKWRGLQPECC | 13 |
| Group47 | | VRNGMSLHDVLMFKLKTRGLQPECC | 20 |
| VRNGMSLHDCLMSKLKWRGLQPECC | 30 | VRNGMSLHDLLMFKLKTRGLQPECC | 14 |
| VRNGMSLHDCLMTKLKWRGLQPECC | 15 | VRNGMSLHDCLMQKLKYRGLQPECC | 252 |
| VRNGMSLHDCLMNKLKWRGLQPECC | 213 | VRNGMSLHDCLMQKLKFRGLQPECC | 63 |
| VRNGMSLHDCLMKKLKWRGLQPECC | 139 | VRNGMSLHDCLMQKLKVRGLQPECC | 26 |
| VRNGMSLHDCLMRKLKWRGLQPECC | 83 | VRNGMSLHDHLMLKLKYRGLQPECC | 13 |
| VRNGMSLHDCLMHKLKWRGLQPECC | 29 | VRNGMSLHDELMECLKARGLQPECC | 19 |
| Group48 | | VRNGMSLHDQLMKPLKYRGLQPECC | 9 |
| VRNGMSLHDTLMYKLKHRGLQPECC | 20 | VRNGMSLHDLLMCKLKTRGLQPECC | 23 |
| VRNGMSLHDTLMFKLKHRGLQPECC | 17 | VRNGMSLHDCLMFKLKNRGLQPECC | 69 |
| VRNGMSLHDILMFKLKHRGLQPECC | 35 | VRNGMSLHDCLMYKLKNRGLQPECC | 35 |
| VRNGMSLHDVLMFKLKHRGLQPECC | 35 | VRNGMSLHDLLMNPLKYRGLQPECC | 25 |
| VRNGMSLHDLLMFKLKHRGLQPECC | 65 | VRNGMSLHDILMNPLKYRGLQPECC | 14 |
| VRNGMSLHDVLMYKLKHRGLQPECC | 56 | VRNGMSLHDLLMRSLKNRGLQPECC | 15 |
| VRNGMSLHDILMYKLKHRGLQPECC | 12 | VRNGMSLHDLLMRALKNRGLQPECC | 14 |
| VRNGMSLHDLLMYKLKHRGLQPECC | 60 | VRNGMSLHDCLMKPLKNRGLQPECC | 117 |
| Group49 | | VRNGMSLHDCLMRPLKNRGLQPECC | 33 |
| VRNGMSLHDLLMKKLKWRGLQPECC | 526 | VRNGMSLHDCLMWKLKNRGLQPECC | 24 |
| VRNGMSLHDMLMKKLKWRGLQPECC | 9 | VRNGMSLHDCLMWKLKTRGLQPECC | 19 |
| VRNGMSLHDVLMKKLKWRGLQPECC | 287 | VRNGMSLHDCLMWKLKHRGLQPECC | 118 |
| VRNGMSLHDILMKKLKWRGLQPECC | 76 | VRNGMSLHDALMNKLKARGLQPECC | 13 |
| VRNGMSLHDTLMKKLKWRGLQPECC | 22 | VRNGMSLHDVLMNKLKARGLQPECC | 18 |
| VRNGMSLHDVLMRKLKWRGLQPECC | 51 | VRNGMSLHDALMFKLKVRGLQPECC | 20 |
| VRNGMSLHDLLMRKLKWRGLQPECC | 39 | VRNGMSLHDALMFKLKIRGLQPECC | 11 |
| VRNGMSLHDLLMQKLKWRGLQPECC | 72 | VRNGMSLHDALMFKLKTRGLQPECC | 11 |
| VRNGMSLHDVLMQKLKWRGLQPECC | 14 | VRNGMSLHDLLMPKLKQRGLQPECC | 10 |
| Group50 | | VRNGMSLHDVLMKKLKCRGLQPECC | 36 |
| VRNGMSLHDLLMWKLKYRGLQPECC | 191 | VRNGMSLHDLLMKKLKCRGLQPECC | 18 |
| VRNGMSLHDMLMWKLKYRGLQPECC | 52 | VRNGMSLHDALMKKLKCRGLQPECC | 14 |
| VRNGMSLHDVLMWKLKYRGLQPECC | 117 | VRNGMSLHDALMQKLKYRGLQPECC | 216 |
| VRNGMSLHDILMWKLKYRGLQPECC | 42 | VRNGMSLHDALMQKLKFRGLQPECC | 72 |
| VRNGMSLHDTLMWKLKYRGLQPECC | 43 | VRNGMSLHDTLMQKLKYRGLQPECC | 127 |

| | | | |
|---|---|---|---|
| VRNGMSLHDLLMWKLKFRGLQPECC | 28 | VRNGMSLHDCLMVKLKTRGLQPECC | 36 |
| VRNGMSLHDVLMWKLKFRGLQPECC | 26 | VRNGMSLHDCLMMKLKTRGLQPECC | 13 |
| VRNGMSLHDALMWKLKYRGLQPECC | 187 | VRNGMSLHDCLMTKLKTRGLQPECC | 21 |
| VRNGMSLHDALMWKLKFRGLQPECC | 14 | VRNGMSLHDHLMKPLKLRGLQPECC | 37 |
| Group51 | | VRNGMSLHDHLMRPLKLRGLQPECC | 11 |
| VRNGMSLHDLLMIKLKQRGLQPECC | 12 | VRNGMSLHDHLMKPLKVRGLQPECC | 11 |
| VRNGMSLHDLLMVKLKQRGLQPECC | 11 | VRNGMSLHDLLMNRLKYRGLQPECC | 19 |
| VRNGMSLHDLLMLKLKQRGLQPECC | 10 | VRNGMSLHDLLMTRLKYRGLQPECC | 13 |
| VRNGMSLHDVLMVKLKQRGLQPECC | 9 | VRNGMSLHDHLMNKLKYRGLQPECC | 47 |
| VRNGMSLHDLLMAKLKQRGLQPECC | 11 | VRNGMSLHDHLMKKLKYRGLQPECC | 23 |
| VRNGMSLHDLLMTKLKQRGLQPECC | 17 | VRNGMSLHDALMPKLKYRGLQPECC | 124 |
| VRNGMSLHDVLMTKLKQRGLQPECC | 15 | VRNGMSLHDALMPKLKFRGLQPECC | 14 |
| Group52 | | VRNGMSLHDTLMPKLKFRGLQPECC | 21 |
| VRNGMSLHDVLMSKLKTRGLQPECC | 31 | VRNGMSLHDLLMNKLKQRGLQPECC | 42 |
| VRNGMSLHDILMSKLKTRGLQPECC | 13 | VRNGMSLHDVLMNKLKQRGLQPECC | 25 |
| VRNGMSLHDVLMSKLKSRGLQPECC | 10 | VRNGMSLHDALMNKLKQRGLQPECC | 12 |
| VRNGMSLHDVLMTKLKTRGLQPECC | 37 | VRNGMSLHDCLMKALKNRGLQPECC | 63 |
| VRNGMSLHDILMTKLKTRGLQPECC | 11 | VRNGMSLHDCLMKSLKNRGLQPECC | 28 |
| VRNGMSLHDVLMAKLKTRGLQPECC | 10 | VRNGMSLHDTLMMKLKFRGLQPECC | 18 |
| VRNGMSLHDLLMSKLKSRGLQPECC | 21 | VRNGMSLHDTLMLKLKFRGLQPECC | 14 |
| VRNGMSLHDLLMTKLKSRGLQPECC | 13 | VRNGMSLHDHLMPKLKYRGLQPECC | 39 |
| VRNGMSLHDLLMSKLKTRGLQPECC | 64 | VRNGMSLHDHLMPKLKHRGLQPECC | 28 |
| VRNGMSLHDLLMTKLKTRGLQPECC | 31 | VRNGMSLHDCLMGKLKYRGLQPECC | 41 |
| Group53 | | VRNGMSLHDCLMGKLKHRGLQPECC | 21 |
| VRNGMSLHDVLMRKLKTRGLQPECC | 75 | VRNGMSLHDCLMYKLKWRGLQPECC | 32 |
| VRNGMSLHDVLMRKLKSRGLQPECC | 14 | VRNGMSLHDCLMPKLKHRGLQPECC | 88 |
| VRNGMSLHDLLMRKLKTRGLQPECC | 51 | VRNGMSLHDCLMPKLKNRGLQPECC | 152 |
| VRNGMSLHDLLMRKLKSRGLQPECC | 15 | VRNGMSLHDCLMHKLKHRGLQPECC | 57 |
| VRNGMSLHDLLMKKLKTRGLQPECC | 193 | VRNGMSLHDCLMHKLKNRGLQPECC | 74 |
| VRNGMSLHDMLMKKLKTRGLQPECC | 18 | VRNGMSLHDLLMKALKRRGLQPECC | 19 |
| VRNGMSLHDLLMKKLKSRGLQPECC | 65 | VRNGMSLHDLLMKSLKRRGLQPECC | 12 |
| VRNGMSLHDVLMKKLKSRGLQPECC | 34 | VRNGMSLHDLLMKGLKRRGLQPECC | 11 |
| VRNGMSLHDILMKKLKSRGLQPECC | 9 | VRNGMSLHDLLMYKLKFRGLQPECC | 24 |
| VRNGMSLHDVLMKKLKTRGLQPECC | 157 | VRNGMSLHDVLMYKLKFRGLQPECC | 25 |
| VRNGMSLHDILMKKLKTRGLQPECC | 44 | VRNGMSLHDLLMFKLKFRGLQPECC | 11 |
| Group54 | | VRNGMSLHDLLMGKLKVRGLQPECC | 39 |
| VRNGMSLHDLLMKPLKFRGLQPECC | 115 | VRNGMSLHDVLMGKLKVRGLQPECC | 21 |
| VRNGMSLHDMLMKPLKFRGLQPECC | 16 | VRNGMSLHDALMGKLKVRGLQPECC | 9 |
| VRNGMSLHDVLMKPLKFRGLQPECC | 152 | VRNGMSLHDCLMYKLKYRGLQPECC | 205 |
| VRNGMSLHDILMKPLKFRGLQPECC | 43 | VRNGMSLHDCLMFKLKYRGLQPECC | 61 |
| VRNGMSLHDILMRPLKFRGLQPECC | 14 | VRNGMSLHDCLMYKLKFRGLQPECC | 12 |
| VRNGMSLHDVLMRPLKFRGLQPECC | 13 | VRNGMSLHDCLMVKLKVRGLQPECC | 9 |
| VRNGMSLHDLLMRPLKFRGLQPECC | 32 | VRNGMSLHDCLMIKLKARGLQPECC | 10 |
| VRNGMSLHDALMKPLKFRGLQPECC | 23 | VRNGMSLHDLLMCKLKHRGLQPECC | 91 |
| VRNGMSLHDTLMKPLKFRGLQPECC | 21 | VRNGMSLHDVLMCKLKHRGLQPECC | 19 |
| Group55 | | VRNGMSLHDALMCKLKHRGLQPECC | 25 |
| VRNGMSLHDCLMNKLKTRGLQPECC | 54 | VRNGMSLHDLLMHKLKWRGLQPECC | 38 |
| VRNGMSLHDCLMNKLKSRGLQPECC | 18 | VRNGMSLHDVLMHKLKWRGLQPECC | 26 |
| VRNGMSLHDCLMNKLKNRGLQPECC | 257 | VRNGMSLHDALMHKLKWRGLQPECC | 24 |
| VRNGMSLHDCLMNKLKLRGLQPECC | 49 | VRNGMSLHDHLMKPLKYRGLQPECC | 178 |
| VRNGMSLHDCLMNKLKMRGLQPECC | 29 | VRNGMSLHDHLMRPLKYRGLQPECC | 17 |
| VRNGMSLHDCLMNKLKVRGLQPECC | 41 | VRNGMSLHDHLMKPLKFRGLQPECC | 9 |
| VRNGMSLHDCLMNKLKIRGLQPECC | 10 | VRNGMSLHDTLMNKLKYRGLQPECC | 230 |
| Group56 | | VRNGMSLHDSLMNKLKYRGLQPECC | 50 |
| VRNGMSLHDCLMKKLKYRGLQPECC | 357 | VRNGMSLHDTLMNKLKFRGLQPECC | 64 |
| VRNGMSLHDCLMKRLKYRGLQPECC | 18 | VRNGMSLHDLLMKSLKYRGLQPECC | 31 |
| VRNGMSLHDCLMRKLKYRGLQPECC | 310 | VRNGMSLHDVLMKSLKYRGLQPECC | 22 |
| VRNGMSLHDCLMKALKYRGLQPECC | 71 | VRNGMSLHDTLMKSLKYRGLQPECC | 13 |
| VRNGMSLHDCLMKSLKYRGLQPECC | 19 | VRNGMSLHDCLMGKLKLRGLQPECC | 11 |
| Group57 | | VRNGMSLHDCLMGKLKVRGLQPECC | 11 |
| VRNGMSLHDALMGKLKNRGLQPECC | 111 | VRNGMSLHDCLMGKLKTRGLQPECC | 18 |
| VRNGMSLHDTLMGKLKNRGLQPECC | 30 | VRNGMSLHDLLMGKLKLRGLQPECC | 16 |
| VRNGMSLHDLLMGKLKNRGLQPECC | 116 | VRNGMSLHDLLMGKLKFRGLQPECC | 11 |
| VRNGMSLHDMLMGKLKNRGLQPECC | 13 | VRNGMSLHDTLMNKLKTRGLQPECC | 30 |
| VRNGMSLHDVLMGKLKNRGLQPECC | 128 | VRNGMSLHDTLMNKLKVRGLQPECC | 16 |
| VRNGMSLHDILMGKLKNRGLQPECC | 40 | VRNGMSLHDTLMNKLKARGLQPECC | 9 |
| VRNGMSLHDFLMGKLKNRGLQPECC | 10 | VRNGMSLHDALMPKLKTRGLQPECC | 13 |

| | | | |
|---|---|---|---|
| Group58 | | VRNGMSLHDALMPKLKVRGLQPECC | 11 |
| VRNGMSLHDVLMQKLKFRGLQPECC | 96 | VRNGMSLHDCLMFKLKTRGLQPECC | 23 |
| VRNGMSLHDILMQKLKFRGLQPECC | 35 | VRNGMSLHDCLMFKLKVRGLQPECC | 10 |
| VRNGMSLHDLLMQKLKFRGLQPECC | 77 | VRNGMSLHDCLMSKLKTRGLQPECC | 23 |
| VRNGMSLHDVLMEKLKFRGLQPECC | 11 | VRNGMSLHDCLMSKLKSRGLQPECC | 9 |
| VRNGMSLHDVLMKKLKFRGLQPECC | 155 | VRNGMSLHDCLMSKLKARGLQPECC | 10 |
| VRNGMSLHDILMKKLKFRGLQPECC | 46 | VRNGMSLHDMLMKPLKWRGLQPECC | 58 |
| VRNGMSLHDVLMRKLKFRGLQPECC | 49 | VRNGMSLHDLLMKPLKWRGLQPECC | 20 |
| VRNGMSLHDLLMKKLKFRGLQPECC | 127 | VRNGMSLHDVLMKPLKWRGLQPECC | 27 |
| VRNGMSLHDLLMRKLKFRGLQPECC | 38 | VRNGMSLHDVLMTKLKLRGLQPECC | 30 |
| Group59 | | VRNGMSLHDLLMTKLKLRGLQPECC | 28 |
| VRNGMSLHDLLMHKLKYRGLQPECC | 245 | VRNGMSLHDLLMSKLKARGLQPECC | 25 |
| VRNGMSLHDMLMHKLKYRGLQPECC | 38 | VRNGMSLHDVLMSKLKARGLQPECC | 10 |
| VRNGMSLHDVLMHKLKYRGLQPECC | 100 | VRNGMSLHDLLMQKLKTRGLQPECC | 17 |
| VRNGMSLHDILMHKLKYRGLQPECC | 80 | VRNGMSLHDVLMQKLKTRGLQPECC | 12 |
| VRNGMSLHDTLMHKLKYRGLQPECC | 35 | VRNGMSLHDLLMKRLKYRGLQPECC | 58 |
| VRNGMSLHDLLMHKLKFRGLQPECC | 25 | VRNGMSLHDVLMKRLKYRGLQPECC | 13 |
| VRNGMSLHDILMHKLKFRGLQPECC | 9 | VRNGMSLHDALMYKLKNRGLQPECC | 35 |
| Group60 | | VRNGMSLHDALMFKLKNRGLQPECC | 27 |
| VRNGMSLHDVLMEKLKHRGLQPECC | 24 | VRNGMSLHDALMDKLKYRGLQPECC | 54 |
| VRNGMSLHDLLMEKLKHRGLQPECC | 12 | VRNGMSLHDALMDKLKFRGLQPECC | 9 |
| VRNGMSLHDLLMDKLKHRGLQPECC | 78 | VRNGMSLHDALMYKLKYRGLQPECC | 150 |
| VRNGMSLHDVLMDKLKHRGLQPECC | 24 | VRNGMSLHDALMYKLKFRGLQPECC | 20 |
| Group61 | | VRNGMSLHDALMYKLKHRGLQPECC | 58 |
| VRNGMSLHDTLMKALKNRGLQPECC | 24 | VRNGMSLHDALMFKLKHRGLQPECC | 24 |
| VRNGMSLHDSLMKALKNRGLQPECC | 10 | VRNGMSLHDVLMNKLKCRGLQPECC | 16 |
| VRNGMSLHDTLMKSLKNRGLQPECC | 28 | VRNGMSLHDLLMNKLKCRGLQPECC | 14 |
| VRNGMSLHDALMKSLKNRGLQPECC | 37 | VRNGMSLHDCLMAKLKYRGLQPECC | 105 |
| VRNGMSLHDALMKALKNRGLQPECC | 12 | VRNGMSLHDCLMAKLKFRGLQPECC | 15 |
| Group62 | | VRNGMSLHDCLMKALKHRGLQPECC | 15 |
| VRNGMSLHDCLMTKLKYRGLQPECC | 377 | VRNGMSLHDCLMKSLKHRGLQPECC | 12 |
| VRNGMSLHDCLMSKLKYRGLQPECC | 198 | VRNGMSLHDCLMFKLKHRGLQPECC | 537 |
| VRNGMSLHDCLMDKLKYRGLQPECC | 50 | VRNGMSLHDCLMYKLKHRGLQPECC | 70 |
| VRNGMSLHDCLMNKLKYRGLQPECC | 317 | VRNGMSLHDCLMWKLKYRGLQPECC | 73 |
| VRNGMSLHDCLMNRLKYRGLQPECC | 12 | VRNGMSLHDCLMWKLKFRGLQPECC | 38 |
| Group63 | | VRNGMSLHDLLMVKLKTRGLQPECC | 45 |
| VRNGMSLHDLLMTKLKHRGLQPECC | 219 | VRNGMSLHDLLMIKLKTRGLQPECC | 24 |
| VRNGMSLHDMLMTKLKHRGLQPECC | 26 | VRNGMSLHDLLMVKLKSRGLQPECC | 9 |
| VRNGMSLHDLLMSKLKHRGLQPECC | 172 | VRNGMSLHDTLMKKLKVRGLQPECC | 41 |
| VRNGMSLHDMLMSKLKHRGLQPECC | 17 | VRNGMSLHDTLMRKLKVRGLQPECC | 14 |
| VRNGMSLHDVLMSKLKHRGLQPECC | 133 | VRNGMSLHDALMKPLKNRGLQPECC | 78 |
| VRNGMSLHDILMSKLKHRGLQPECC | 83 | VRNGMSLHDALMRPLKNRGLQPECC | 31 |
| VRNGMSLHDVLMTKLKHRGLQPECC | 162 | VRNGMSLHDCLMKKLKQRGLQPECC | 111 |
| VRNGMSLHDILMTKLKHRGLQPECC | 43 | VRNGMSLHDCLMRKLKQRGLQPECC | 24 |
| VRNGMSLHDVLMAKLKHRGLQPECC | 74 | VRNGMSLHDHLMKPLKNRGLQPECC | 87 |
| VRNGMSLHDILMAKLKHRGLQPECC | 25 | VRNGMSLHDHLMRPLKNRGLQPECC | 29 |
| VRNGMSLHDLLMAKLKHRGLQPECC | 106 | VRNGMSLHDCLMKPLKHRGLQPECC | 122 |
| Group64 | | VRNGMSLHDCLMRPLKHRGLQPECC | 13 |
| VRNGMSLHDLLMMKLKFRGLQPECC | 35 | VRNGMSLHDALMTKLKVRGLQPECC | 12 |
| VRNGMSLHDLLMLKLKFRGLQPECC | 19 | VRNGMSLHDALMSKLKVRGLQPECC | 11 |
| VRNGMSLHDLLMVKLKFRGLQPECC | 12 | VRNGMSLHDMLMKGLKYRGLQPECC | 10 |
| VRNGMSLHDVLMLKLKFRGLQPECC | 53 | VRNGMSLHDLLMKGLKYRGLQPECC | 9 |
| VRNGMSLHDVLMMKLKFRGLQPECC | 39 | VRNGMSLHDKLMHCLKVRGLQPECC | 10 |
| VRNGMSLHDVLMVKLKFRGLQPECC | 55 | VRNGMSLHDRLMMFLKIRGLQPECC | 10 |
| VRNGMSLHDVLMIKLKFRGLQPECC | 9 | VRNGMSLHDYLMQVLKIRGLQPECC | 11 |
| Group65 | | VRNGMSLHDGLMVDLKMRGLQPECC | 9 |
| VRNGMSLHDALMNKLKYRGLQPECC | 282 | VRNGMSLHDLLMLPLKYRGLQPECC | 9 |
| VRNGMSLHDALMNKLKFRGLQPECC | 242 | VRNGMSLHDCLMNPLKYRGLQPECC | 11 |
| VRNGMSLHDALMNKLKVRGLQPECC | 42 | VRNGMSLHDHLMKPLKHRGLQPECC | 51 |
| VRNGMSLHDALMNKLKIRGLQPECC | 18 | VRNGMSLHDHLMKPLKRRGLQPECC | 18 |
| VRNGMSLHDALMNKLKLRGLQPECC | 37 | VRNGMSLHDTLMKPLKLRGLQPECC | 10 |
| Group66 | | VRNGMSLHDGLMKPLKYRGLQPECC | 10 |
| VRNGMSLHDTLMLKLKWRGLQPECC | 21 | VRNGMSLHDLLMKGLKNRGLQPECC | 18 |
| VRNGMSLHDTLMMKLKWRGLQPECC | 16 | VRNGMSLHDLLMEKLKYRGLQPECC | 39 |
| VRNGMSLHDTLMVKLKWRGLQPECC | 10 | VRNGMSLHDCLMKPLKRRGLQPECC | 9 |
| VRNGMSLHDTLMYKLKWRGLQPECC | 16 | VRNGMSLHDCLMRDLKWRGLQPECC | 11 |
| VRNGMSLHDTLMFKLKWRGLQPECC | 11 | VRNGMSLHDCLMWKLKWRGLQPECC | 56 |

| | | | |
|---|---|---|---|
| Group67 | | VRNGMSLHDALMKNLKWRGLQPECC | 10 |
| VRNGMSLHDTLMSKLKNRGLQPECC | 75 | VRNGMSLHDLLMKRLKWRGLQPECC | 16 |
| VRNGMSLHDTLMTKLKNRGLQPECC | 67 | VRNGMSLHDHLMPKLKVRGLQPECC | 11 |
| VRNGMSLHDTLMAKLKNRGLQPECC | 35 | VRNGMSLHDHLMPKLKNRGLQPECC | 25 |
| VRNGMSLHDTLMMKLKNRGLQPECC | 83 | VRNGMSLHDHLMKALKYRGLQPECC | 80 |
| VRNGMSLHDTLMLKLKNRGLQPECC | 67 | VRNGMSLHDHLMCKLKYRGLQPECC | 9 |
| VRNGMSLHDTLMIKLKNRGLQPECC | 29 | VRNGMSLHDHLMVKLKFRGLQPECC | 11 |
| VRNGMSLHDTLMVKLKNRGLQPECC | 28 | VRNGMSLHDCLMWALKKRGLQPECC | 10 |
| Group68 | | VRNGMSLHDCLMGKLKNRGLQPECC | 76 |
| VRNGMSLHDALMSKLKNRGLQPECC | 121 | VRNGMSLHDCLMCKLKYRGLQPECC | 43 |
| VRNGMSLHDALMTKLKNRGLQPECC | 104 | VRNGMSLHDCLMIKLKRRGLQPECC | 19 |
| VRNGMSLHDALMAKLKNRGLQPECC | 60 | VRNGMSLHDLLMPKLKCRGLQPECC | 19 |
| VRNGMSLHDALMLKLKNRGLQPECC | 120 | VRNGMSLHDVLMAKLKCRGLQPECC | 27 |
| VRNGMSLHDALMMKLKNRGLQPECC | 86 | VRNGMSLHDCLMKKLKCRGLQPECC | 188 |
| VRNGMSLHDALMIKLKNRGLQPECC | 63 | VRNGMSLHDQLMKILKCRGLQPECC | 9 |
| VRNGMSLHDALMVKLKNRGLQPECC | 42 | VRNGMSLHDLLMCKLKWRGLQPECC | 11 |
| Group69 | | VRNGMSLHDILMCKLKIRGLQPECC | 19 |
| VRNGMSLHDLLMAKLKFRGLQPECC | 19 | VRNGMSLHDALMFKLKCRGLQPECC | 24 |
| VRNGMSLHDVLMAKLKFRGLQPECC | 16 | VRNGMSLHDDLMFKLKYRGLQPECC | 17 |
| VRNGMSLHDLLMAKLKYRGLQPECC | 149 | VRNGMSLHDTLMMKLKVRGLQPECC | 17 |
| VRNGMSLHDMLMAKLKYRGLQPECC | 21 | VRNGMSLHDLLMWKLKTRGLQPECC | 11 |
| VRNGMSLHDVLMAKLKYRGLQPECC | 215 | VRNGMSLHDVLMNKLKRRGLQPECC | 12 |
| VRNGMSLHDILMAKLKYRGLQPECC | 69 | VRNGMSLHDCLMCFLKARGLQPECC | 10 |
| | | VRNGMSLHDILMEYLKPRGLQPECC | 10 |
| | | VRNGMSLHDDLMIYLKDRGLQPECC | 9 |
| | | VRNGMSLHDWLMWCLKDRGLQPECC | 19 |
| | | VRNGMSLHDDLMWILKWRGLQPECC | 19 |
| | | VRNGMSLHDILMWWLKARGLQPECC | 10 |

**Supplementary Table 4. Converged Positions within each cluster.** X indicates lack of 100% convergence. Binary indicates a 0 for each position with an X and a 1 for each position not X.

| Cluster | Binary | 81 | 84 | 85 | 88 |
|---|---|---|---|---|---|
| Group60 | 0001 | X | X | X | Y |
| Group6 | 0010 | X | X | K | X |
| Group20 | 0010 | X | X | K | X |
| Group37 | 0010 | X | X | P | X |
| Group52 | 0010 | X | X | K | X |
| Group53 | 0010 | X | X | K | X |
| Group4 | 0011 | X | X | K | N |
| Group7 | 0011 | X | X | K | W |
| Group11 | 0011 | X | X | K | A |
| Group12 | 0011 | X | X | K | H |
| Group15 | 0011 | X | X | K | Q |
| Group16 | 0011 | X | X | K | Y |
| Group21 | 0011 | X | X | K | Y |
| Group23 | 0011 | X | X | K | Y |
| Group24 | 0011 | X | X | K | Y |
| Group25 | 0011 | X | X | K | V |
| Group38 | 0011 | X | X | P | H |
| Group39 | 0011 | X | X | K | D |
| Group40 | 0011 | X | X | P | N |
| Group42 | 0011 | X | X | K | N |
| Group48 | 0011 | X | X | K | H |
| Group49 | 0011 | X | X | K | W |
| Group54 | 0011 | X | X | P | F |
| Group58 | 0011 | X | X | K | F |
| Group63 | 0011 | X | X | K | H |
| Group64 | 0011 | X | X | K | F |
| Group72 | 0011 | X | X | K | Y |
| Group74 | 0011 | X | X | K | N |
| Group77 | 0011 | X | X | K | H |
| Group82 | 0011 | X | X | K | Y |
| Group84 | 0011 | X | X | K | N |
| Group85 | 0011 | X | X | K | Y |
| Group88 | 0011 | X | X | P | Y |
| Group91 | 0011 | X | X | K | W |
| Group98 | 0011 | X | X | K | W |
| Group101 | 0011 | X | X | K | F |
| Group118 | 0011 | X | X | K | N |
| Group5 | 0101 | X | K | X | N |
| Group41 | 0101 | X | K | X | H |
| Group61 | 0101 | X | K | X | N |
| Group8 | 0110 | X | P | K | X |
| Group10 | 0110 | X | C | K | X |
| Group13 | 0110 | X | N | K | X |
| Group26 | 0110 | X | N | K | X |
| Group31 | 0110 | X | R | K | X |
| Group33 | 0110 | X | P | K | X |
| Group36 | 0110 | X | M | K | X |
| Group43 | 0110 | X | D | K | X |
| Group44 | 0110 | X | Q | K | X |
| Group50 | 0110 | X | W | K | X |
| Group59 | 0110 | X | H | K | X |
| Group69 | 0110 | X | A | K | X |
| Group71 | 0110 | X | N | K | X |
| Group80 | 0110 | X | K | K | X |
| Group83 | 0110 | X | P | K | X |
| Group112 | 0110 | X | G | K | X |
| Group116 | 0110 | X | H | K | X |
| Group1 | 0111 | X | K | P | Y |
| Group2 | 0111 | X | Q | K | Y |

| Cluster | Binary | 81 | 84 | 85 | 88 |
|---|---|---|---|---|---|
| Group3 | 0111 | X | W | K | N |
| Group19 | 0111 | X | K | A | Y |
| Group57 | 0111 | X | G | K | N |
| Group70 | 0111 | X | N | K | N |
| Group78 | 0111 | X | P | K | N |
| Group79 | 0111 | X | N | K | H |
| Group86 | 0111 | X | Q | K | N |
| Group89 | 0111 | X | H | K | N |
| Group90 | 0111 | X | Q | K | H |
| Group92 | 0111 | X | H | K | H |
| Group93 | 0111 | X | P | K | H |
| Group94 | 0111 | X | G | K | H |
| Group95 | 0111 | X | N | K | W |
| Group97 | 0111 | X | G | K | Y |
| Group99 | 0111 | X | C | K | N |
| Group100 | 0111 | X | W | K | H |
| Group102 | 0111 | X | W | K | W |
| Group104 | 0111 | X | K | P | R |
| Group113 | 0111 | X | D | K | N |
| Group114 | 0111 | X | P | K | W |
| Group62 | 1001 | C | X | X | Y |
| Group9 | 1010 | C | X | K | X |
| Group18 | 1010 | C | X | P | X |
| Group28 | 1010 | C | X | K | X |
| Group29 | 1010 | C | X | K | X |
| Group30 | 1010 | A | X | K | X |
| Group73 | 1010 | A | X | K | X |
| Group87 | 1010 | A | X | K | X |
| Group96 | 1010 | C | X | K | X |
| Group103 | 1010 | C | X | K | X |
| Group111 | 1010 | A | X | K | X |
| Group14 | 1011 | C | X | K | F |
| Group17 | 1011 | A | X | K | T |
| Group22 | 1011 | V | X | K | T |
| Group34 | 1011 | C | X | K | N |
| Group35 | 1011 | C | X | K | H |
| Group45 | 1011 | A | X | K | F |
| Group46 | 1011 | A | X | K | W |
| Group47 | 1011 | C | X | K | W |
| Group51 | 1011 | L | X | K | Q |
| Group66 | 1011 | T | X | K | W |
| Group67 | 1011 | T | X | K | N |
| Group68 | 1011 | A | X | K | N |
| Group75 | 1011 | A | X | K | H |
| Group76 | 1011 | T | X | K | H |
| Group105 | 1011 | C | X | K | N |
| Group106 | 1011 | C | X | K | H |
| Group107 | 1011 | C | X | K | W |
| Group117 | 1011 | A | X | K | W |
| Group56 | 1101 | C | K | X | Y |
| Group108 | 1101 | A | K | X | Y |
| Group115 | 1101 | A | K | X | H |
| Group27 | 1110 | C | H | K | X |
| Group32 | 1110 | C | P | K | X |
| Group55 | 1110 | C | N | K | X |
| Group65 | 1110 | A | N | K | X |
| Group81 | 1110 | T | P | K | X |
| Group109 | 1110 | T | K | K | X |
| Group110 | 1110 | A | H | K | X |

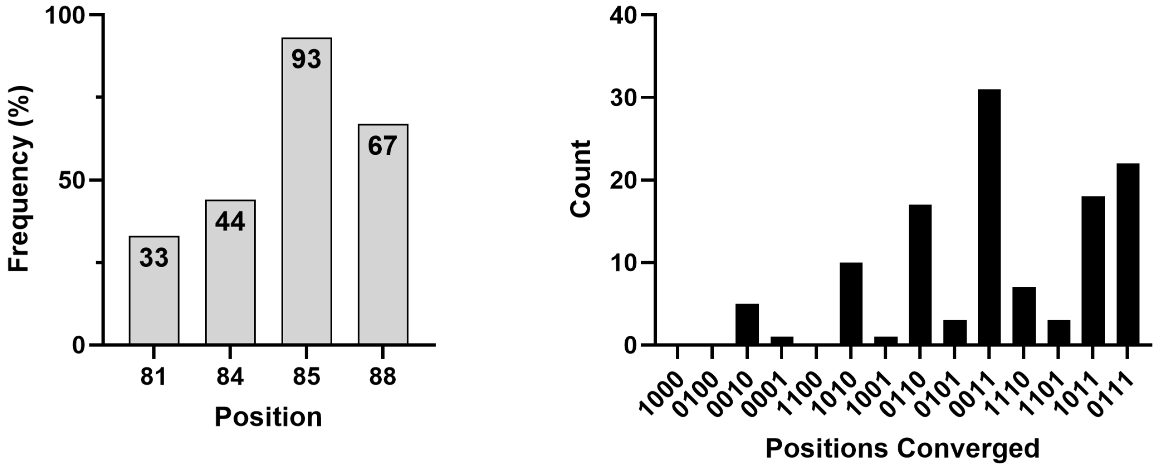

**Supplementary Figure 17. Convergence in Clusters by randomized position (Supplementary Table 4).** Frequency of convergence for each position (left). Number of clusters with each of the possible convergence patterns (right).

**Supplementary Table 5. Algorithms for navigating the pen-Raf fitness landscape.** (example of 1 and 2 shown diagrammatically below)

**1. Greedy Algorithm – Select the best option for each possible order of specification.**

| Position 1 | Position 2 | Position 3 | Position 4 | 81 | 84 | 85 | 88 | Slope |
|---|---|---|---|---|---|---|---|---|
| 81 | 84 | 85 | 88 | L | K | K | W | 109 |
| 81 | 84 | 88 | 85 | L | K | P | Y | 56 |
| 81 | 85 | 88 | 84 | L | R | P | Y | 65 |
| 81 | 85 | 84 | 88 | L | K | K | W | 109 |
| 81 | 88 | 84 | 85 | L | K | P | Y | 56 |
| 81 | 88 | 85 | 84 | L | R | K | Y | 65 |
| 84 | 81 | 85 | 88 | L | K | K | W | 109 |
| 84 | 81 | 88 | 85 | L | K | P | Y | 56 |
| 84 | 85 | 81 | 88 | L | K | K | W | 109 |
| 84 | 85 | 88 | 81 | T | K | K | Y | 86 |
| 84 | 88 | 81 | 85 | L | K | P | Y | 56 |
| 84 | 88 | 85 | 81 | M | K | P | Y | 69 |
| 85 | 81 | 84 | 88 | L | K | K | W | 109 |
| 85 | 81 | 88 | 84 | L | R | K | Y | 65 |
| 85 | 84 | 81 | 88 | L | K | K | W | 109 |
| 85 | 84 | 88 | 81 | T | K | K | Y | 86 |
| 85 | 88 | 81 | 84 | L | R | K | Y | 65 |
| 85 | 88 | 84 | 81 | T | K | K | Y | 86 |
| 88 | 81 | 84 | 85 | L | K | P | Y | 56 |
| 88 | 81 | 85 | 84 | L | R | K | Y | 65 |
| 88 | 84 | 81 | 85 | L | K | P | Y | 56 |
| 88 | 84 | 85 | 81 | M | K | P | Y | 69 |
| 88 | 85 | 81 | 84 | L | R | K | Y | 65 |
| 88 | 85 | 84 | 81 | T | K | K | Y | 86 |

**2. Mutations from WT – Select the best point mutation for each possible order of specification.**

| Position 1 | Position 2 | Position 3 | Position 4 | 81 | 84 | 85 | 88 | Slope |
|---|---|---|---|---|---|---|---|---|
| 81 | 84 | 85 | 88 | L | K | K | W | 109 |
| 81 | 84 | 88 | 85 | L | K | K | N | 59 |
| 81 | 85 | 88 | 84 | L | K | K | W | 109 |
| 81 | 85 | 84 | 88 | L | K | K | W | 109 |
| 81 | 88 | 84 | 85 | L | K | K | N | 59 |
| 81 | 88 | 85 | 84 | L | K | K | N | 59 |
| 84 | 81 | 85 | 88 | L | K | K | W | 109 |
| 84 | 81 | 88 | 85 | L | K | K | N | 59 |
| 84 | 85 | 81 | 88 | V | K | K | H | 82 |
| 84 | 85 | 88 | 81 | C | K | K | C | 108 |
| 84 | 88 | 81 | 85 | H | K | P | Y | 44 |
| 84 | 88 | 85 | 81 | T | K | K | Y | 86 |
| 85 | 81 | 84 | 88 | V | K | K | H | 82 |
| 85 | 81 | 88 | 84 | V | K | K | H | 82 |
| 85 | 84 | 81 | 88 | V | K | K | H | 82 |
| 85 | 84 | 88 | 81 | C | K | K | C | 108 |
| 85 | 88 | 81 | 84 | C | K | K | C | 108 |
| 85 | 88 | 84 | 81 | C | K | K | C | 108 |
| 88 | 81 | 84 | 85 | H | K | P | Y | 44 |
| 88 | 81 | 85 | 84 | H | K | P | Y | 44 |
| 88 | 84 | 81 | 85 | H | K | P | Y | 44 |
| 88 | 84 | 85 | 81 | T | K | K | Y | 86 |
| 88 | 85 | 81 | 84 | T | K | K | Y | 86 |
| 88 | 85 | 84 | 81 | C | M | K | Y | 87 |

**3. Substantial (+6 slope) improvement mutational pathways from WT** (mutation is bolded)

| Mutation 1 | Mutation 2 | Mutation 3 | Mutation 4 | Mutation 5 | Mutation 6 | 81-84-85-88 | Slope |
|---|---|---|---|---|---|---|---|
| CK**K**V - 26 | CKK**C** - 108 | | | | | CKKC | 108 |
| CK**K**V - 26 | CKK**N** - 67 | CKK**C** -108 | | | | CKKC | 108 |
| CK**K**V - 26 | CKK**N** - 67 | C**S**KN -75 | | | | CSKN | 75 |
| CK**K**V - 26 | CKK**Y** - 52 | CKK**C** -108 | | | | CKKC | 108 |
| CK**K**V - 26 | CKK**Y** - 52 | C**M**KY - 87 | | | | CMKY | 87 |
| CK**K**V - 26 | CKK**Y** - 52 | **T**KKY – 86 | | | | TKKY | 86 |
| CK**K**V - 26 | CKK**Y** - 52 | C**T**KY – 83 | | | | CTKY | 83 |

| | | | | | | | |
|---|---|---|---|---|---|---|---|
| CK**K**V - 26 | CKK**Y** - 52 | C**L**KY – 78 | | | | CLKY | 78 |
| CK**K**V - 26 | CKK**Y** - 52 | C**Q**KY - 64 | | | | CQKY | 64 |
| CK**K**V - 26 | CKK**H** - 49 | CKK**C** -108 | | | | CKKC | 108 |
| CK**K**V - 26 | CKK**H** - 49 | **V**KKH - 82 | | | | VKKH | 82 |
| CK**K**V - 26 | CKK**H** - 49 | C**M**KH - 75 | CMK**Y** - 87 | | | CMKY | 87 |
| CK**K**V - 26 | CKK**H** - 49 | C**F**KH - 55 | | | | CFKH | 55 |
| CK**K**V - 26 | **V**KKV - 38 | VKK**H** - 82 | | | | VKKH | 82 |
| CK**K**V - 26 | **V**KKV - 38 | VKK**W** - 59 | **L**KKW - 109 | | | LKKW | 109 |
| CK**K**V - 26 | **V**KKV - 38 | VKK**N** - 58 | **C**KKN - 67 | CKK**C** -108 | | CKKC | 108 |
| CK**K**V - 26 | CKK**F** - 38 | CKK**C** - 108 | | | | CKKC | 108 |
| CK**K**V - 26 | CKK**F** - 38 | **A**KKF - 87 | | | | AKKF | 87 |
| CK**K**V - 26 | CKK**F** - 38 | C**N**KF - 68 | | | | CNKF | 68 |
| CK**K**V - 26 | CKK**Q** - 37 | CKK**C** -108 | | | | CKKC | 108 |
| CK**K**V - 26 | CKK**Q** - 37 | **I**KKQ -68 | | | | IKKQ | 68 |
| CKA**Y** - 23 | CK**K**Y - 52 | CKK**C** -108 | | | | CKKC | 108 |
| CKA**Y** - 23 | CK**K**Y - 52 | C**M**KY - 87 | | | | CMKY | 87 |
| CKA**Y** - 23 | CK**K**Y - 52 | **T**KKY – 86 | | | | TKKY | 86 |
| CKA**Y** - 23 | CK**K**Y - 52 | C**T**KY – 83 | | | | CTKY | 83 |
| CKA**Y** - 23 | CK**K**Y - 52 | C**L**KY – 78 | | | | CLKY | 78 |
| CKA**Y** - 23 | CK**K**Y - 52 | C**Q**KY - 64 | | | | CQKY | 64 |
| CKA**Y** - 23 | CK**P**Y - 50 | **M**KPY - 69 | | | | MKPY | 69 |
| CKA**Y** - 23 | CK**P**Y - 50 | **T**KPY - 62 | TK**K**Y - 86 | | | TKKY | 86 |
| CKA**Y** - 23 | **H**KAY - 38 | HK**P**Y - 44 | **M**KPY - 69 | | | MKPY | 69 |
| CKA**Y** - 23 | **H**KAY - 38 | HK**P**Y - 44 | **T**KPY - 62 | TK**K**Y - 86 | | TKKY | 86 |
| CKA**Y** - 23 | **H**KAY - 38 | HK**P**Y - 44 | **L**KPY - 56 | **M**KPY - 69 | | MKPY | 69 |
| CKA**N** - 20 | **L**KAN - 30 | LK**K**N – 59 | LKK**W** – 109 | | | LKKW | 109 |
| CKA**N** - 20 | **L**KAN - 30 | LK**K**N – 59 | **C**KKN - 67 | CKK**C** – 108 | | CKKC | 108 |
| CKA**N** - 20 | **L**KAN - 30 | LK**P**N - 46 | LKP**Y** - 56 | **M**KPY - 69 | | MKPY | 69 |

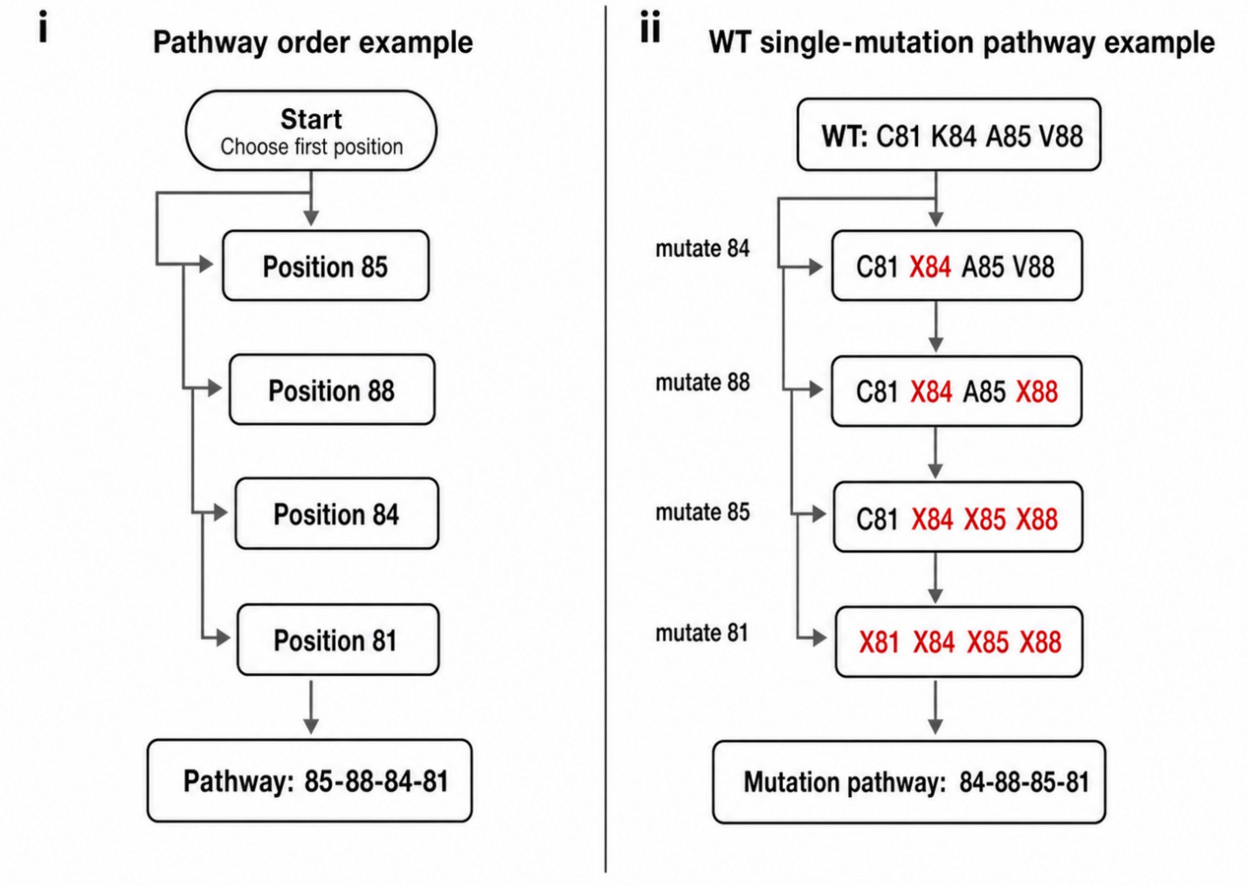

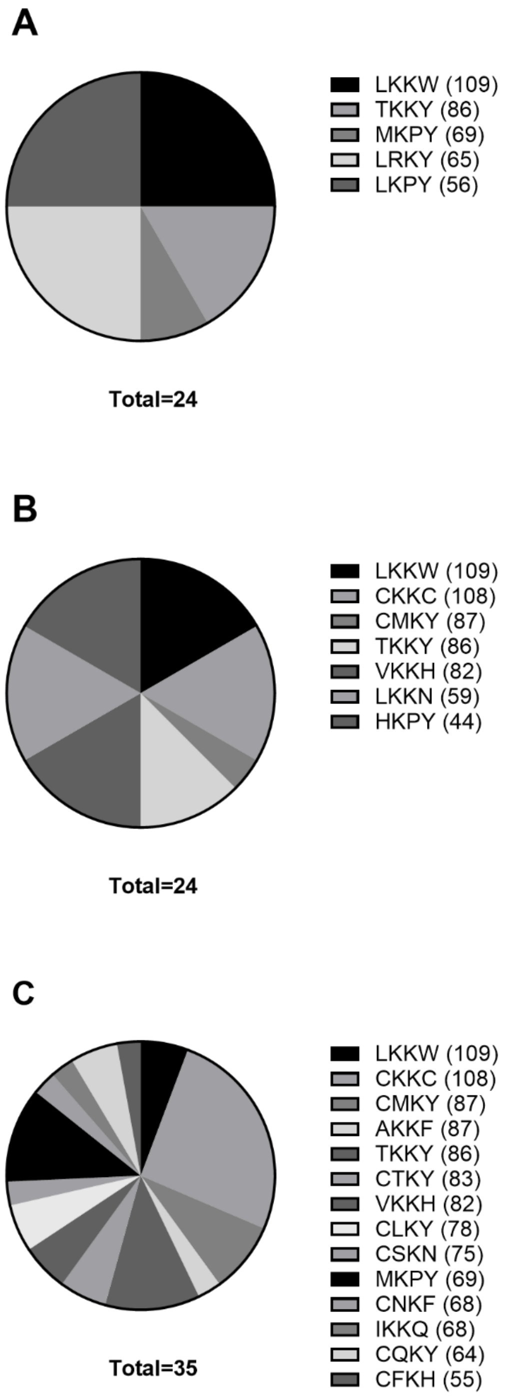

**Supplementary Figure 18. Summary of algorithmic specification of variant results (Supplementary Table 5): A)** Greedy algorithm, **B)** best point mutation for all possible routes from WT**, and C)** point mutation paths from WT that must have > +6 in slope from prior variant. Normalized slopes listed next to variant (81/84/85/88).

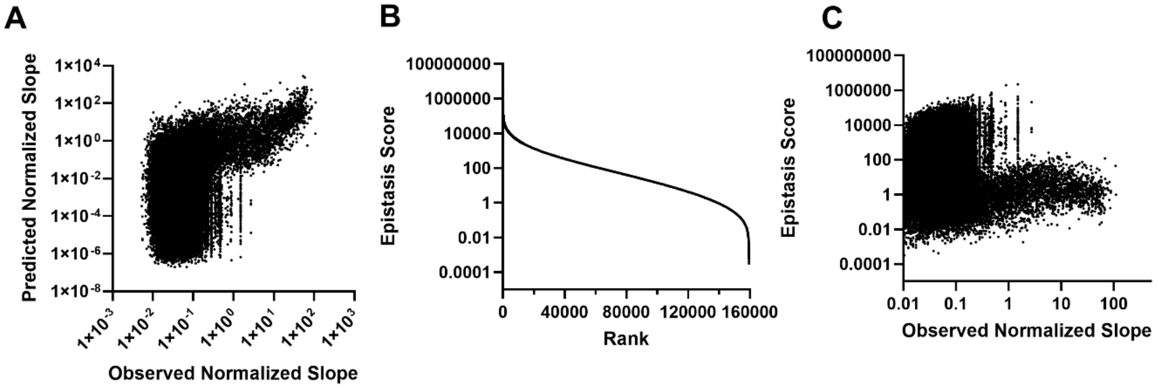

**Supplementary Figure 19. Predicting Fitness without a lower bound. A)** Predicted fitness based only on single point mutant fitness data for all possible variants (without any lower bound on predicted fitness). Pearson's Correlation, r = 0.282, p = <0.001. **B**) Epistasis Score (Observed Fitness/Predicted Fitness) for all variants from **A**. **C**) Epistasis Score vs the Observed Normalized Slope for all variants in **A** shows that nearly all variants >3 have epistasis values <1000.

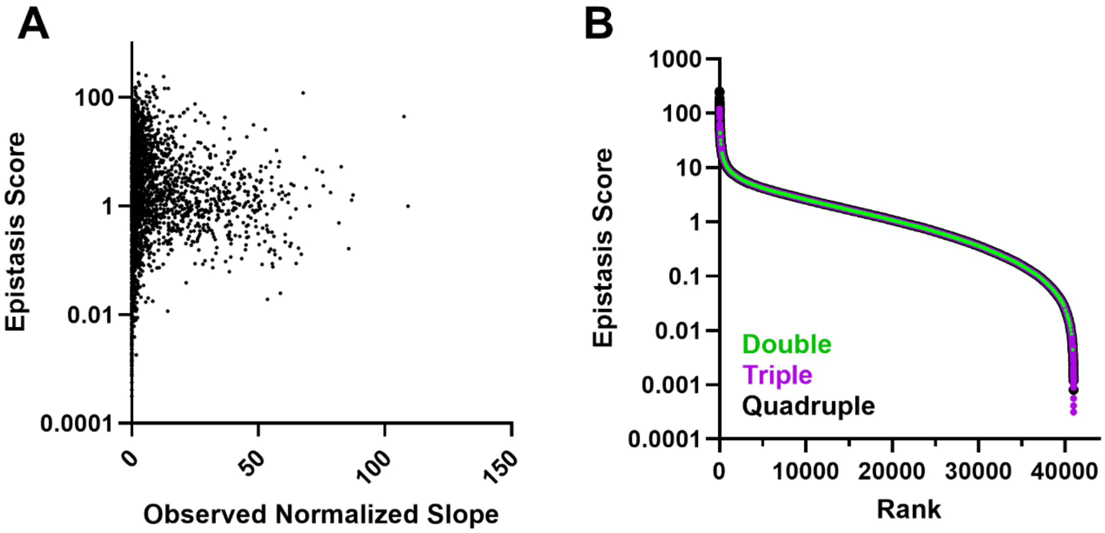

**Supplementary Figure 20. Epistasis using single mutants with a 0.01 Predicted Slope floor. A)** Epistasis vs Observed Normalized Slope. **B)** Epistasis (only variants with Predicted Slopes >0.01) separated by mutant type – double, triple, or quadruple.

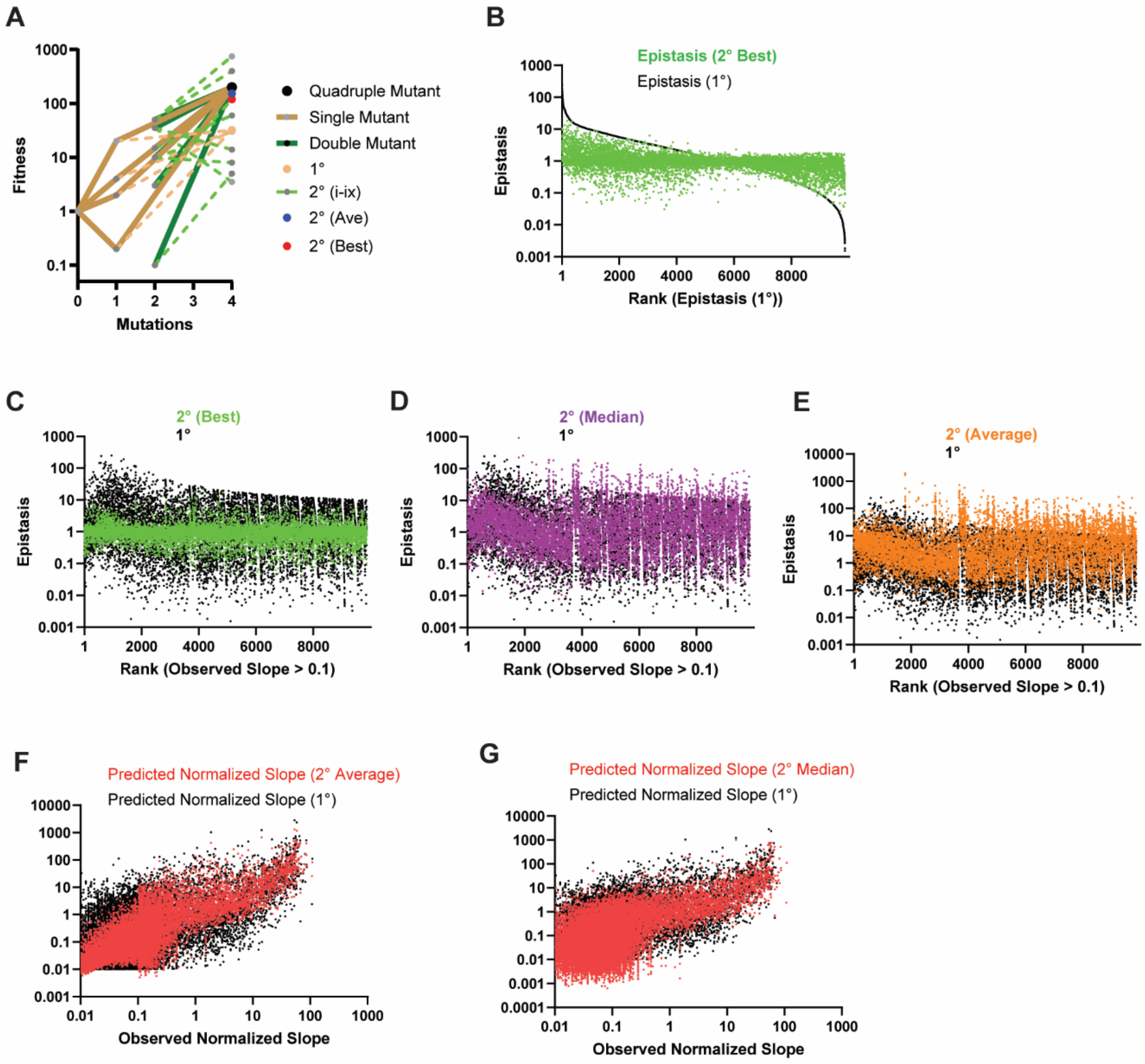

**Supplementary Figure 21. Predicting Fitness with single and double mutant data. A)** Predicting the fitness (normalized slope) of quadruple mutant result in a unique fitness prediction (tan lines): for each of the four single mutations away from WT (normalized slope = 1), each point mutant has a different fitness (grey points at 1 mutation) and only one way of combining the set of four point mutants to predict a single 1° fitness (tan point with dashed tan lines). The difference between the actual fitness of the quadruple mutant and the 1° fitness is the 1° epistasis score. For double mutations (grey points at the origin of green lines), there are 6 combinations (ab, ac, ad, bc, bd, cd) and 9 possible combinations for combining (ab+cd, ab+c+d, ac+bd, ac+b+d, ad+bc, ad+b+c, bc+a+d, bd+a+c, cd+a+b) this data to make a 2° prediction (dashed green lines leading to 9 different predictions of fitness). 2° epistasis scores are the difference between the observed fitness and each of these predicted fitness values. The average, median, and "best" 2° predictions/epistasis scores are determined from these 9 predictions. "Best" is determined by the predicted fitness value that is closest to the observed fitness value. **B)** The 1° and best 2° epistasis scores for all variants with > 0.1 observed normalized slope are shown with the rank-order of the 1° epistasis scores. The "best" 2° epistasis scores are much closer to 1 at both tails in the 1° epistasis scores. **C-E**) The 1° epistasis scores (black) and 2° epistasis scores (colored, **C** – Best, **D** – Median, **E** – Average) for all variants with > 0.1 observed normalized slope (in the rank order of observed slope). Error is minimized throughout the "best" 2° epistasis scores and for the most active variants in the median and average 2° epistasis scores. **F-G**) Observed vs predicted slope for all variants with an observed normalized slope >0.01 comparing (black) 1° predicted slope to (red) 2° predicted slope (**F**- average; **G** – median).

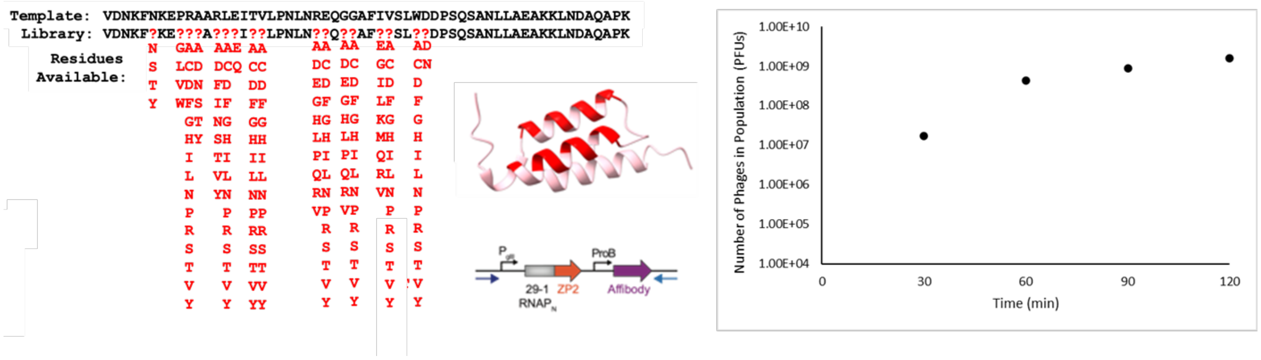

**Supplementary Figure 22. Affibody Library.** The design (left) of the affibody library used in this work is identical to our previously reported affibody library design (randomized residues shown on the structure in red)[2], but with the affibody in the PANCS-Inhibitor SP design (middle). The total PFU of the transformed library was 2x10$^7$ at 30 minutes and 4x10$^8$ at 60 minutes, and 2x10$^9$ at 120 minutes; the 30 and 120 minute titers are lower and upper bounds on library size, while the 60 minute value is likely on the order of magnitude of the diversity encoded in this library (~10$^8$ unique variants).

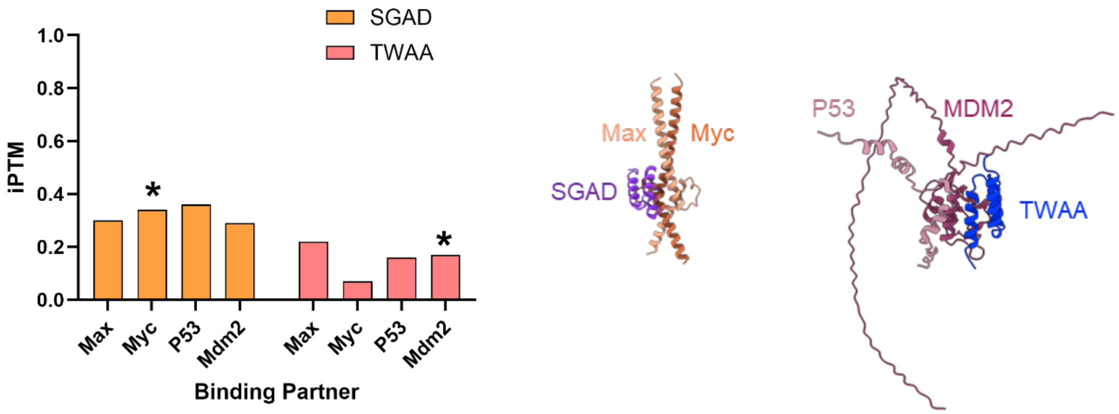

**Supplementary Figure 23. AlphaFold 3 predictions of *de novo* inhibitors.** *(Left)* The PPI between each inhibitor with each target was predicted using AlphaFold 3 and the iPTM values (all low confidence, <0.5) are shown. * indicates PPIs that actually form; those structures are shown *(right)*.

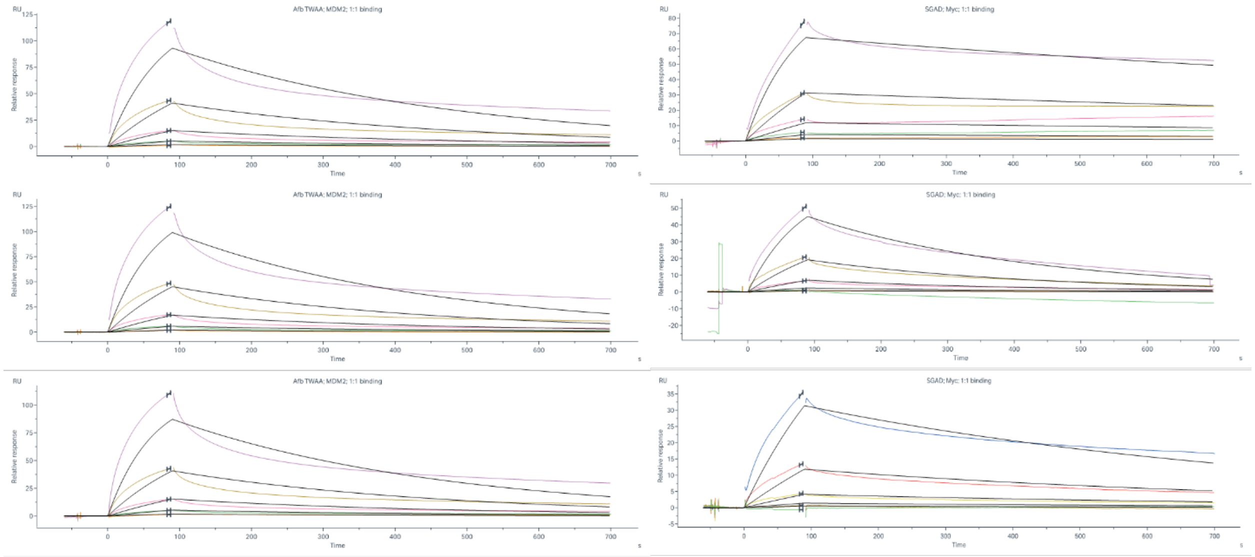

| Interaction | $k_a$ (1/Ms) | $k_d$ (1/s) | $K_d$ (nM) |
|---|---|---|---|
| TWAA:Mdm2 | 3.73E+03 | 2.55E-03 | 684 |
| | 4.23E+03 | 2.81E-03 | 664 |
| | 4.65E+03 | 2.68E-03 | 576 |
| **Average** | **4.20E+03** | **2.68E-03** | **638** |

| Interaction | $k_a$ (1/Ms) | $k_d$ (1/s) | $K_d$ (nM) |
|---|---|---|---|
| SGAD:Myc | 4.28E+03 | 5.13E-04 | 120 |
| | 3.16E+03 | 2.93E-03 | 927 |
| | 1.49E+03 | 1.36E-03 | 913 |
| **Average** | **2.98E+03** | **1.60E-03** | **538** |

**Supplementary Figure 24. SPR results.** Triplicate (left) GST-TWAA dose response curves (3000, 1000, 333, 111, 37, 0 nM) on Mdm2-His6 and (right) GST-SGAD dose response curves (3000, 1000, 333, 111, 37, 0 nM) on Myc-His6 loaded NTA chip. Tables below each set indicate kinetic (1:1) binding fits.

**Supplementary Table 6. Plasmids.**

| Figure | Plasmid ID | Description | Benchling Link |
|---|---|---|---|
| 1D/S2 | 37-66 | p15A CGG SD8 lux 2xVSV pKat SD8 KRas(wt)-C(CGG) | https://benchling.com/s/seq-LP4ry6CVER4NvNTjE0Kt?m=slm-THZPiyZwIOgaLC3Dy14N |
| 1D/S2 | 66-144 | p15A CGG SD8 lux 2xVSV pKat SD8 MDM2-C(CGG) | https://benchling.com/s/seq-TNg7uIsbUFU8xLocAB8k?m=slm-XmExXPcYYIxwfBiAaajA |
| 1D/S2 | 66-176 | p15A CGG SD8 lux 2xVSV pKat SD8 Max-C(CGG) | https://benchling.com/s/seq-tya4t3Bi5bl7V0U0hEEu?m=slm-83OkugpJTZTaRgNPyYtx |
| 1D/S2 | 38-50 | pBR322 origin sd8 P(J23117) N(29-1)-Raf | https://benchling.com/s/seq-HYFMs3vBUTEE7bVBiScF?m=slm-Q0hJa5k2pYCetntXrCih |
| 1D/S2 | 66-135 | pBR322 origin sd8 P(J23117) N(29-1)-P53 | https://benchling.com/s/seq-AQxHv2ElTKnS8cHP9WaJ?m=slm-5BSWjhaSt3Cie42qJ1Tu |
| 1D/S2 | 66-124 | pBR322 origin sd8 P(J23117) N(29-1)-Myc | https://benchling.com/s/seq-tdi9EJRWWEgj4xYtZEt9?m=slm-95Xa9PUWs8qYyhXOqvvC |
| 1F/1G/S2 | 19-30 | cloDF13 IPTG inducible inhibitor - RAF (general design for all inhibitors in inhibitor E. coli lux) | https://benchling.com/s/seq-pgkatsKNqeB6jynJ3BjX?m=slm-C32H40nA3czfbRdXdoQ3 |
| 1F/1G/S2 | De Novo 1 | cloDF13 IPTG inducible inhibitor(general design for all de novo binders/inhibitors) | https://benchling.com/s/seq-55yylpEoOl66ueJgAF75?m=slm-0fxUM81LtkbJgiuLTTdW |
| 2B | 74-185 | pBR322 origin sd8 P(J23117) N(29-1)-synzip1 | https://benchling.com/s/seq-PLDqnZXtJMFNg68RikYd?m=slm-X3wexMGwVhde6O9ZvHI5 |
| 2B/5F/G | 44-63 | p15A CGG SD8 Lux 2xVSV pKat sd8 ZB-C(CGG) | https://benchling.com/s/seq-7rrUOjIEN87XgKlEwC2T?m=slm-ns3tCfWcqgFpDpAqfQFi |
| 2B | 74-184 | cloDF13 IPTG inducible synzip2-ZA-Raf (general deisng for adaptor) | https://benchling.com/s/seq-J3BJcytj2BOkzU6Ib6W5?m=slm-jQwhwnzyrv1eDwAfW1P0 |
| 2B | 74-191 | cloDF13 IPTG inducible synzip1 | https://benchling.com/s/seq-BSaIu1UbKror5W5DlcpF?m=slm-w2M2X8wrlCmkux8Yhrva |
| 2B | 74-192 | cloDF13 IPTG inducible synzip2-ZA | https://benchling.com/s/seq-SUIr8xXHsU4RDQ52C09s?m=slm-RhgGuDgx2HMtcD90CaNY |
| 2B | 74-193 | cloDF13 IPTG inducible synzip2-Raf | https://benchling.com/s/seq-mp5B2Fc9ALhziQBmViDb?m=slm-89fszrXyB7wNHCHBpNOr |
| 2B | 74-194 | cloDF13 IPTG inducible ZA-Raf | https://benchling.com/s/seq-3OytbYUf970s7gRUm9a9?m=slm-N1EtgJ3n7YE0m1bceq3k |
| 2D | 73-119 | Empty phage: pgIII 29-1N-term-synzip1-CE-SP gen2 | https://benchling.com/s/seq-1VpK2dY2XFc4rwhXFsC2?m=slm-gaw9oMmyNJOdFcJvhvnp |
| 2D | 53-78 | Control phage: pgIII 29-1N-term-ZA-SP | https://benchling.com/s/seq-4RS1WHVqO3j7t3mx8320?m=slm-2amdnQRn2voGPzQJIeFj |

| | | | |
|---|---|---|---|
| 2D | 71-158 | pgIII 29-1N-term-synzip1 ProB Affibody(Raf), gen2 | https://benchling.com/s/seq-OhxGgVosvGO9nhv55J6g?m=slm-7fSZwVe56R9tYKC2Qpc7 |
| 2D | 71-156 | pgIII 29-1N-term-synzip1 ProB CI2(MDM2), gen2 | https://benchling.com/s/seq-7Xh7qqJsKVihNQbJSt9U?m=slm-EBzTlnkrbBL6m6V0mYS3 |
| 2D | 71-151 | pgIII 29-1N-term-synzip1 ProB MAX gen2 | https://benchling.com/s/seq-44yC030HCBLubj49h9OO?m=slm-tNX7IBklJ6xzv7U1JDbv |
| 2F/3B/S6 | 73-120 | p15A, J23114 SD8 synzip2-ZA-Raf, CGG-driven SD8 gIII recoded, 2xVSV, pKat SD8 ZB-RNAPC-CGG | https://benchling.com/s/seq-2VUO83wnNKTeuLCcbDvA?m=slm-PVqNov6NZyfln4IXuAKO |
| 2F/3B/S6 | 59-42 | negAP, pBR322, pT7 SD8 gIIIneg, 2xVSV terminator, pKat sd8 KRas-T7-C-term-RNAP | https://benchling.com/s/seq-PhvD6E3TyypVs2jyEJTp?m=slm-wuwqyPS7XyS6vFsXbjgT |
| 2F/5C/S6 | 71-111 | p15A, J23114 SD8 synzip2-ZA-P53, CGG-driven SD8 gIII recoded, 2xVSV, pKat SD8 ZB-RNAPC-CGG | https://benchling.com/s/seq-RDsJFiub9DAO1ak8z7lO?m=slm-vrBuflkA84buOLZV32fU |
| 2F/5C/S6 | 71-141 | negAP, pBR322, pT7 sd8 gIIIneg, 2xVSV terminator, pKat sd5 MDM2-T7-C-term-RNAP | https://benchling.com/s/seq-mdgXNuo9kmGOzUwjuvQB?m=slm-pk9J1zhLgkEgym9mlvHL |
| 2F/5B/S6 | 71-143 | p15A, J23114 SD8 synzip2-ZA-MYC_full length, CGG-driven SD8 gIII recoded, 2xVSV, pKat SD8 ZB-RNAPC-CGG | https://benchling.com/s/seq-1rnWT6wOlkrf1vn2vvOY?m=slm-PytJCE2xSPpB4XECwUcO |
| 2F/5B/S6 | 71-186 | negAP, pBR322, pT7 SD8 gIIIneg, 2xVSV terminator, pKat SD8 MAX_new-T7-C-term-RNAP | https://benchling.com/s/seq-PZBeSfcxgFxAQQE7FC5o?m=slm-NOK5Ee9rB8O9ZQ9BOdBQ |
| 3B | SL-7 | pen-RAF 4 NNK library SP | https://benchling.com/s/seq-ZxwHh150GP4Dw8Att5mI?m=slm-DTNRqBqLXEanlio92zg8 |
| 3D | 66-155 | cloDF13 IPTG inducible Pen-Raf (general inducible inhibitor design) | https://benchling.com/s/seq-wXSwIuChYJAhJRL5vPIR?m=slm-X4o8kwaq1O00rxYYDwog |
| 3F/3G | 74-107 | CMV Nluc11S-Raf CMV KRas(WT)-Nluc114 | https://benchling.com/s/seq-fuJ8SOECowq9s4TXbDFc?m=slm-EeB0oeRR2SuFGEdqwh1X |
| 3F/5H/5I | 74-108 | CMV Raf (general design for mammailan inhibitor expression) | https://benchling.com/s/seq-yrHyk59ELh706bWrOKjQ?m=slm-l0JWn4RNESdGGqnp7JUx |
| 3G/3I | 84-40 | pET28a His-penRAF (general design for all pen-inhibitors for purification) | https://benchling.com/s/seq-7Saa4pypEzZ7hbMVwoHs?m=slm-gHiRgO79KBtwwPPUE8RC |
| 5B/5C | SL-9 | Affibody inhibitor library SP | https://benchling.com/s/seq-1zs8gU1OtlLFnGoZHXqm?m=slm-QRzz01MlSmOg3E8v1Odt |
| 5D/E | De Novo 1 | cloDF13 IPTG inducible inhibitor(general design for all de novo binders/inhibitors) | https://benchling.com/s/seq-55yylpEoOl66ueJgAF75?m=slm-0fxUM81LtkbJgiuLTTdW |
| 5D | 66-177 | p15A CGG SD8 lux 2xVSV pKat SD8 Myc-C(CGG) | https://benchling.com/s/seq-lUCZcBhbavUDA1CtT6kh?m=slm-dwjjKQR7UPRTBvmASVCk |

| | | | |
|---|---|---|---|
| 5E | 66-147 | p15A CGG SD8 lux 2xVSV pKat SD8 P53_short-C(CGG) | https://benchling.com/s/seq-N5K6z4NRCQLeddbnLTmu?m=slm-MKRQ3f8W5H8KfJZjuR7R |
| 5F/G | 84-43 | SD8 P(J23114) N(29-1)-pen-RAF (general design for binder lux) | https://benchling.com/s/seq-GoNuNuriY3ZuLcIxme0y?m=slm-qlltf76COBEPUP1uofck |
| 5F/G | De Novo 3 | RNAPNwt-inhibitor (general) (expression lux) | https://benchling.com/s/seq-TsSoF6j5rIpJzEFnMtf1?m=slm-1qWOEKzxJQQmj4oll9wm |
| 5H | 84-41 | CMV Nluc11S-MYC_long CMV MAX_new-Nluc114 | https://benchling.com/s/seq-l0urMpeMilFtzXRlQH2O?m=slm-nVA8Aw917PZjb8MSZ0l9 |
| 5I | 84-42 | CMV Nluc11S-P53_short CMV MDM2-Nluc114 | https://benchling.com/s/seq-JFOWYAvhR56zfGHWwZXL?m=slm-uJzSU9Mehym3Qj79dCsm |
| S6 | 71-123 | p15A, J23114 SD8 synzip2-Noxa-Raf, CGG-driven SD8 gIII recoded, 2xVSV, pKat SD8 ZB-RNAPC-CGG | https://benchling.com/s/seq-gZL9joN3RL37RQzbzxaE?m=slm-220GPVeR43E4AHsQVnJD |
| S6 | 71-125 | p15A, J23114 SD8 synzip2-ZA-Raf, CGG-driven SD8 gIII recoded, 2xVSV, pKat SD8 Noxa-RNAPC-CGG | https://benchling.com/s/seq-rlgGrVqMnVE77D5XArbs?m=slm-usI3qzNzRR2PB8qgOrt9 |
| S6 | 59-43 | negAP, pBR322, pT7 sd8 gIIIneg, 2xVSV terminator, pKat sd8 KRas-T7-C-term-RNAP | https://benchling.com/s/seq-jwLYtCXHwNWMoecN5OSe?m=slm-W4NyHUWMgDx7UGWzTHqJ |
| S6 | 66-82 | negAP, pBR322, pT7 SD8 gIIIneg, 2xVSV terminator, pKat SD8 KRas-T7-C-term-RNAP | https://benchling.com/s/seq-7SbKuXTDmwMLYnGOlO45?m=slm-IbUGvX5GeofqRr5UgVlj |
| S6 | 71-114 | pgIII 29-1N-term-synzip1 ProB MDM2 gen2 | https://benchling.com/s/seq-MLbHRAWYu1PXKXl6Y2qh?m=slm-TePuttKl1rXf7DJHGShK |
| S6 | 73-121 | pgIII 29-1N-term-synzip1 ProB Pen-Raf gen2 | https://benchling.com/s/seq-ZkVnAtQ6NQClBfOSzxcF?m=slm-ku15P2obM4oTv6mjNhgv |
| S6 | 23-49 | negAP, pBR322 T7 sd8 gIIIneg, 2xVSV terminator, pKat sd8 NhNoxa-C(T7)RNAP | https://benchling.com/s/seq-C85LhxB4si9zNCtuDwMf?m=slm-sKb5EGxjK81LN8dnje9V |
| S6 | jin 487 | negAP, PBR322, terminator-NhNOXA neg C term T7-SD8-sd8 | https://benchling.com/s/seq-PkjVpHuRp0qRcRZxVo5Y?m=slm-gUiHuIqJKMZkkhR8RQBX |
| S6 | 28-67 | negAP, pBR322, terminator-NhNOXA neg C term T7-SD8-SD8 | https://benchling.com/s/seq-dUvkwb4t6enS4CPg7MEK?m=slm-j25S2heQXd8dMxx3pL7x |
| S6 | 59-55 | negAP, pBR322, pT7 sd5 gIIIneg, 2xVSV terminator, pKat sd2 KRas-T7-C-term-RNAP | https://benchling.com/s/seq-9LF4EfsQuE83QYbcyg1F?m=slm-pwM8nmGPebGjkMdu7ire |
| S6 | 66-95 | negAP, pBR322, pT7 sd5 gIIIneg, 2xVSV terminator, pKat sd5 KRas-T7-C-term-RNAP | https://benchling.com/s/seq-4NKc4qDEKZ3kdOAcHEeO?m=slm-J30ui564TTCyqzDEcPvu |
| S6 | 66-94 | negAP, pBR322, pT7 sd8 gIIIneg, 2xVSV terminator, pKat sd5 KRas-T7-C-term-RNAP | https://benchling.com/s/seq-TTOHla5REb4p6kzH42eg?m=slm-dEtpECXGS751OavDPqzZ |

| | | | |
|---|---|---|---|
| S6 | 73-122 | pgIII 29-1N-term-synzip1 ProB Pen-cRaf-v1 gen2 | https://benchling.com/s/seq-HD7AZATb0T1n1i8eUPAa?m=slm-mHky3bnG0z0poxQMvXoI |
| S6 | 73-123 | pgIII 29-1N-term-synzip1 ProB Pen-Raf(Q66A) gen2 | https://benchling.com/s/seq-2fm1O2akkQzjqDWyIwR9?m=slm-k9FK6r2GxfbJDziVSafB |
| S6 | 73-118 | pgIII 29-1N-term-synzip1 ProB Raf(R89L) gen2 | https://benchling.com/s/seq-5kAn5dwdPrQqKc9qHH6r?m=slm-7HzzAYB5M990BuzUXIe6 |
| S6 | 71-157 | pgIII 29-1N-term-synzip1 ProB Monobody(KRas), gen2 | https://benchling.com/s/seq-BAPyvXD2j1DWtb33pcRe?m=slm-zyoGbcLV5JcYyQ6wQvu3 |
| S6 | 71-158 | pgIII 29-1N-term-synzip1 ProB Affibody(Raf), gen2 | https://benchling.com/s/seq-NV5W3AvIHxA1eylNEEuD?m=slm-VuBGGZ8K3Mu7x4BYYl4r |
| S6 | 71-134 | negAP, pBR322, pT7 sd5 gIIIneg, 2xVSV terminator, pKat sd2 MDM2-T7-C-term-RNAP | https://benchling.com/s/seq-35FvQt9NM4el4TTB457H?m=slm-lY9rM3eOufk6yospf0i3 |
| S6 | 71-135 | negAP, pBR322, pT7 sd8 gIIIneg, 2xVSV terminator, pKat sd2 MDM2-T7-C-term-RNAP | https://benchling.com/s/seq-Y6erxxkDYQgdp4LRORS7?m=slm-Lic9Y6Jso4wHXmfEQMcB |
| S6 | 71-136 | negAP, pBR322, pT7 SD8 gIIIneg, 2xVSV terminator, pKat sd2 MDM2-T7-C-term-RNAP | https://benchling.com/s/seq-ImHPV2g8SUKw2wQA57Mw?m=slm-NKhS5b9i3wQnaoEQ8Lem |
| S6 | 71-140 | negAP, pBR322, pT7 sd5 gIIIneg, 2xVSV terminator, pKat sd5 MDM2-T7-C-term-RNAP | https://benchling.com/s/seq-T4EtpjdFsKaTjwCHyRWL?m=slm-eVmHb83bVfaljvbaTCuX |
| S6 | 66-169 | negAP, pBR322, pT7 sd8 gIIIneg, 2xVSV terminator, pKat sd8 MDM2-T7-C-term-RNAP | https://benchling.com/s/seq-hG3ug4ktvzYiRYhTnjof?m=slm-ioG5bOFwk2A5oeEMRWzW |
| S6 | 66-168 | negAP, pBR322, pT7 SD8 gIIIneg, 2xVSV terminator, pKat sd8 MDM2-T7-C-term-RNAP | https://benchling.com/s/seq-ZivHZfa0QmRNMK8nvbVZ?m=slm-N4EF1cPhtbKsdtdw9kJo |
| S6 | 66-167 | negAP, pBR322, pT7 SD8 gIIIneg, 2xVSV terminator, pKat SD8 MDM2-T7-C-term-RNAP | https://benchling.com/s/seq-DKmaVzseCO6WBo0Iu6zi?m=slm-bntj1oh8V9ANfEz3Hbqi |
| S6 | 71-115 | pgIII 29-1N-term-synzip1 ProB P53short gen2 | https://benchling.com/s/seq-zOOs3Vap256J6I7QIHzI?m=slm-p5SRBo7Y2CrhqJCED4yh |
| S6 | 71-156 | pgIII 29-1N-term-synzip1 ProB CI2(MDM2), gen2 | https://benchling.com/s/seq-Mq3eg5g2LqIEbj7VK4NH?m=slm-7JWzuO0vwv6E9njb531C |
| S6 | 71-144 | p15A, J23114 SD8 synzip2-ZA-MYC_DBD, CGG-driven SD8 gIII recoded, 2xVSV, pKat SD8 ZB-RNAPC-CGG | https://benchling.com/s/seq-TUMW6bwGSyWrjooVUHIV?m=slm-7O80bsGdGfIZZavPR0Lq |
| S6 | 71-145 | p15A, J23114 SD8 synzip2-ZA-MAX_new, CGG-driven SD8 gIII recoded, 2xVSV, pKat SD8 ZB-RNAPC-CGG | https://benchling.com/s/seq-NtVxmfPk5cb1uU1UDjEU?m=slm-PuBM2CiO8EM0I8TNrYac |
| S6 | 71-192 | negAP, pBR322, pT7 sd8 gIIIneg, 2xVSV terminator, pKat sd8 MAX-T7-C-term-RNAP | https://benchling.com/s/seq-clFQ5zifVivozqZT3r1M?m=slm-AU9LA3LuL2oiPk4bhz2U |

| | | | |
|---|---|---|---|
| S6 | 71-189 | negAP, pBR322, pT7 SD8 gIIIneg, 2xVSV terminator, pKat sd8 MAX_new-T7-C-term-RNAP | https://benchling.com/s/seq-5aM8djKccHTJ16MuBKZH?m=slm-ceMiYIUpOvhQ5JYy5hwr |
| S6 | 71-149 | pgIII 29-1N-term-synzip1 ProB MYC full length gen2 | https://benchling.com/s/seq-HpPQoRU2fAtCya0sf36E?m=slm-0r0v2ADyetGdg9hCIZkU |
| S6 | 71-150 | pgIII 29-1N-term-synzip1 ProB MYC gen2 | https://benchling.com/s/seq-d8HqK9pYE4meA1VopEbs?m=slm-NuoGtivMOG4BnOiDp9RK |
| S6 | 71-155 | pgIII 29-1N-term-synzip1 ProB Telobody(myc), gen2 | https://benchling.com/s/seq-e8WM2TWEF8kn5VkjTLmH?m=slm-bvrMvB7d6tBWw7SJz2kn |
| S24 | 73-72 | pET Myc-H6 | https://benchling.com/s/seq-fWmpFHCQ7b5RAfyJ68ae?m=slm-jIfeGZ6JVDpxlz64zNbg |
| S24 | 73-73 | pET Mdm2-H6 | https://benchling.com/s/seq-gmnUxnm2bj8rhyPwSktd?m=slm-xxHJ9rdh3gMwZobFgjPa |
| S24 | De Novo 2 | pET30-3xFLAG-GST-TEV-Affibody variant (general design) | https://benchling.com/s/seq-HJ2Zn5GkjjbZDvszxeGB?m=slm-loELToDk8iQaILZt6G7g |

### Supplementary Table 7. Primers.

| ID | Use | Sequence |
|---|---|---|
| BR-76 | Monitoring selection de-enrichment | atgaacacgattaacatcgctaagaacgacttc |
| JD-1060 | | ccgattgagggagcatgttgaaaatctcca |
| MS-617 | Primers for randomizing Affibody for library construction | CATCATgctagcGTAGATAATAAGTTTAACAAAGAANNKNNKNNKGCANNKNNKGAAATTNNKNNKCTGCCGAACTTAAACNNKNNKCAGNNKNNKGCCTTCATCNNKAGCCTGNNKGATGACCCGTCCCAAAGCGCTAATTTGCTGG |
| MS-619 | | CATCATgctagcCATATGTATATCTCCTTCTTAAAGTTAAAAAAGTTAAACAAAATTATTTGTAGAGGGAAACCGTTGTGGTCTCCC |
| SL-89 | Primers for randomizing pen-Raf for library construction | catcatctgcagccggagtgttgtgcggtctttcgtctgc |
| SL-95 | | catcatctgcagaccacgMNNtttcagMNNMNNcatcagMNNgtcatgcaggctcataccgttgcgcacattaacgacggtacg |
| SL-0124 | HT NGS Forward primers to add random barcode from pen-Raf PCR template (first round PCR for sample prep) | GTGACTGGAGTTCAGACGTGtgctcttccgatctNNNNNNGTGCGCAACGGTATGAGC |
| SL-0125 | | GTGACTGGAGTTCAGACGTGtgctcttccgatctNNNNNNNGTGCGCAACGGTATGAGC |
| SL-0126 | | GTGACTGGAGTTCAGACGTGtgctcttccgatctNNNNNNNNGTGCGCAACGGTATGAGC |
| SL-0127 | | GTGACTGGAGTTCAGACGTGtgctcttccgatctNNNNNNNNNGTGCGCAACGGTATGAGC |
| SL-0128 | HT NGS Reverse primers to add random barcode from pen-Raf PCR template (first round PCR for sample prep) | ACACTCTTTCCCTACACGACgctcttccgatctNNNNNNACAACACTCCGGCTGCAG |
| SL-0129 | | ACACTCTTTCCCTACACGACgctcttccgatctNNNNNNNACAACACTCCGGCTGCAG |
| SL-0130 | | ACACTCTTTCCCTACACGACgctcttccgatctNNNNNNNNACAACACTCCGGCTGCAG |
| SL-0131 | | ACACTCTTTCCCTACACGACgctcttccgatctNNNNNNNNNACAACACTCCGGCTGCAG |
| D701 | Forward (second round PCR) with unique index and Illumina adaptor for HT NGS (Fig. 3) | CAAGCAGAAGACGGCATACGAGATcgagtaatGTGACTGGAGTTCAGACGTG |
| D702 | | CAAGCAGAAGACGGCATACGAGATtctccggaGTGACTGGAGTTCAGACGTG |
| D703 | | CAAGCAGAAGACGGCATACGAGATaatgagcgGTGACTGGAGTTCAGACGTG |
| D704 | | CAAGCAGAAGACGGCATACGAGATggaatctcGTGACTGGAGTTCAGACGTG |
| D705 | | CAAGCAGAAGACGGCATACGAGATttctgaatGTGACTGGAGTTCAGACGTG |
| D706 | | CAAGCAGAAGACGGCATACGAGATacgaattcGTGACTGGAGTTCAGACGTG |
| D707 | | CAAGCAGAAGACGGCATACGAGATagcttcagGTGACTGGAGTTCAGACGTG |
| D708 | | CAAGCAGAAGACGGCATACGAGATgcgcattaGTGACTGGAGTTCAGACGTG |
| D709 | | CAAGCAGAAGACGGCATACGAGATcatagccgGTGACTGGAGTTCAGACGTG |
| D501 | Reverse (second round PCR) for HT NGS | AATGATACGGCGACCACCGAGATCTACACtatagcctACACTCTTTCCCTACACGAC |
| MS-1479 | Forward barcoded primers for NGS of de novo PANCS-Inhibitor selections (Fig. 5) | acactctttccctacacgacgctcttccgatctAAGGTTgagaccacaacggtttccctctac |
| MS-1480 | | acactctttccctacacgacgctcttccgatctTAGGATgagaccacaacggtttccctctac |
| MS-1481 | | acactctttccctacacgacgctcttccgatctTGAGATgagaccacaacggtttccctctac |
| MS-1482 | | acactctttccctacacgacgctcttccgatctAGTGATgagaccacaacggtttccctctac |
| MS-1483 | | acactctttccctacacgacgctcttccgatctAAGTGTgagaccacaacggtttccctctac |
| MS-1484 | | acactctttccctacacgacgctcttccgatctAAGTTGgagaccacaacggtttccctctac |
| MS-1485 | | acactctttccctacacgacgctcttccgatctAGATGTgagaccacaacggtttccctctac |
| MS-1486 | | acactctttccctacacgacgctcttccgatctAGTAGTgagaccacaacggtttccctctac |
| MS-1487 | | acactctttccctacacgacgctcttccgatctGAATTGgagaccacaacggtttccctctac |
| MS-1488 | | acactctttccctacacgacgctcttccgatctGATATGgagaccacaacggtttccctctac |
| MS-1489 | | acactctttccctacacgacgctcttccgatctGTAATGgagaccacaacggtttccctctac |
| MS-1490 | | acactctttccctacacgacgctcttccgatctGTATAGgagaccacaacggtttccctctac |
| MS-1491 | Reverse of NGS | gactggagttcagacgtgtgctcttccgatctgggcgacattcaaccgattgag |

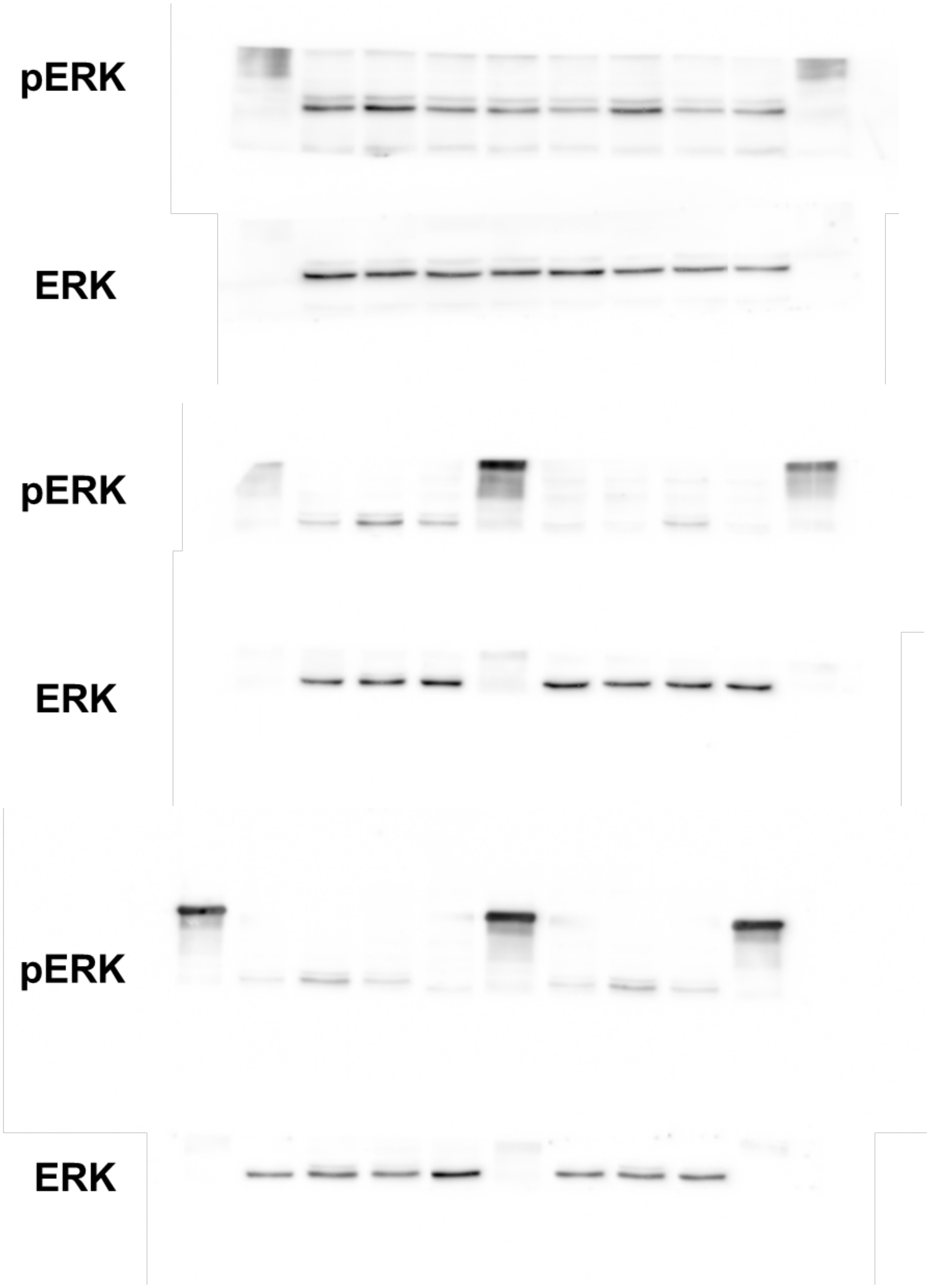

**Supplementary Fig. 25: Full WB Images from Fig. S13.** The pERK antibody stains the ladder.
